# Mutational robustness changes during long-term adaptation in laboratory budding yeast populations

**DOI:** 10.1101/2021.12.17.473185

**Authors:** Milo S. Johnson, Michael M. Desai

**Affiliations:** Department of Organismic and Evolutionary Biology, Center for Mathematical and Statistical Analysis of Biology, Harvard University, Cambridge MA 02138; Quantitative Biology Initiative, Center for Mathematical and Statistical Analysis of Biology, Harvard University, Cambridge MA 02138; NSF-Simons Center for Mathematical and Statistical Analysis of Biology, Harvard University, Cambridge MA 02138; Department of Physics, Harvard University, Cambridge MA 02138

## Abstract

As an adapting population traverses the fitness landscape, its local neighborhood (i.e., the collection of fitness effects of single-step mutations) can change shape because of interactions with mutations acquired during evolution. These changes to the distribution of fitness effects can affect both the rate of adaptation and the accumulation of deleterious mutations. However, while numerous models of fitness landscapes have been proposed in the literature, empirical data on how this distribution changes during evolution remains limited. In this study, we directly measure how the fitness landscape neighborhood changes during laboratory adaptation. Using a barcode-based mutagenesis system, we measure the fitness effects of 91 specific gene disruption mutations in genetic backgrounds spanning 8,000-10,000 generations of evolution in two constant environments. We find that the mean of the distribution of fitness effects decreases in one environment, indicating a reduction in mutational robustness, but does not change in the other. We show that these distribution-level patterns result from biases in variable patterns of epistasis at the level of individual mutations, including fitness-correlated and idiosyncratic epistasis.

## Introduction

Evolutionary adaptation relies on recombination and spontaneous mutagenesis to constantly introduce variation into populations, upon which natural selection can act. The fate of a single mutation - and its impacts on the dynamics of adaptation - depends on how it affects organismal fitness, which we know depends in complex ways on the rest of the genetic background (reviewed in de Visser et al., 2011; Domingo et al., 2019; Lehner, 2011). Understanding this genotype-dependence, or epistasis, involves analyzing key features of the fitness landscape, the high-dimensional map between genotype and fitness (Wright, 1932).

To investigate these questions, numerous studies have surveyed epistatic interactions among a variety of different types of mutations. These studies have found that beneficial mutations isolated from laboratory evolution experiments tend to exhibit negative epistasis. That is, they are less beneficial in combination than would be expected from the combination of their effects in isolation (Karkare et al., 2021; MacLean et al., 2010; Ono et al., 2017; Rokyta et al., 2011; Schenk et al., 2013; Schoustra et al., 2016; Zee and Velicer, 2017). However, some examples of positive epistasis among beneficial mutations have also been observed (Chou et al., 2009; Fumasoni and Murray, 2020; Hsieh et al., 2020; Khan et al., 2011; Levin-Reisman et al., 2019). Surveys of interactions between deleterious mutations have produced numerous examples of both positive and negative epistasis (Costanzo et al., 2016; Elena and Lenski, 1997; Hall and MacLean, 2011; Jasnos and Korona, 2007; Lalić and Elena, 2012; Sanjuán et al., 2004; Van Leuven et al., 2021).

While this earlier work has identified a wide range of epistatic interactions among specific combinations of mutations, several recent studies of epistasis in laboratory microbial systems have found that general patterns of *fitness-correlated epistasis* often emerge. These fitness-correlated patterns typically favor less-fit genotypes: for both beneficial and deleterious mutations, we usually find that the fitness effect of a mutation is negatively (rather than positively) correlated with the fitness of the genetic background on which it occurs (Chou et al., 2011; Johnson et al., 2019; Khan et al., 2011; Kryazhimskiy et al., 2014). These patterns have been termed *diminishing returns* and *increasing costs* for beneficial and deleterious mutations respectively. These trends are particularly intriguing to those interested in modeling the dynamics of adaptation for two reasons. The first is the analytical promise of a simple, linear relationship between a set of difficult-to-model parameters (the fitness effects of mutations) and a single, centrally-important variable, fitness. The second is the potential for these patterns of epistasis to explain declining adaptability, a commonly observed phenomenon in laboratory evolution experiments in which the rate of fitness increase slows as evolution proceeds (reviewed in Couce and Tenaillon, 2015, see Wünsche et al., 2017 for an analysis of the link between diminishing returns and declining adaptability).

The initial studies that identified patterns of fitness-mediated epistasis did so in the context of a relatively small number of beneficial mutations, finding that these mutations became systematically less beneficial in more-fit genetic backgrounds. More recent work has shown that as populations evolve, the average fitness effect of a spontaneous beneficial mutation decreases (Aggeli et al., 2021; Wünsche et al., 2017). Much less work has been done to characterize how the effects of deleterious mutations interact with the beneficial mutations that fix during evolution (Remold and Lenski, 2004). We recently showed that the fitness effects of larger panels of ∼100 to ∼1000 random insertion mutations (both beneficial and deleterious) tend to be systematically less beneficial or more deleterious in more-fit backgrounds (Johnson et al., 2019). However, the genetic backgrounds in that study were derived from a cross between two diverged yeast strains. It remains unclear whether similar patterns would hold across genetic backgrounds that are the result of long-term laboratory evolution, because the mutations that drive evolutionary adaptation are selected along the line of descent, which in principle could bias their epistatic interactions.

Here, we directly address this question by measuring the fitness effects of a panel of insertion mutations during the course of a long-term laboratory evolution experiment in budding yeast. Specifically, we isolated random clones from 6 timepoints spanning 8,000 to 10,000 generations of adaptation to each of two constant laboratory environments from the ongoing evolution experiment we have recently described (Johnson et al., 2021). While the yeast strains used in our prior experiment differed at tens of thousands of segregating loci, the strains in this experiment differ by only tens or hundreds of mutations that fixed during evolution. By looking at the effects of insertion mutations in these strains, we are measuring a panel of hidden phenotypes that may change predictably or stochastically during evolution. The widespread presence of epistasis observed in biological systems suggests that these hidden phenotypes may be important to long-term evolutionary outcomes as the fitness effects of mutations effectively open and close doors to unique pathways for evolution (Johnson et al., 2021; Karkare et al., 2021; Kvitek and Sherlock, 2011).

Our approach complements several recent studies that have analyzed changes in mutational robustness during evolution by conducting mutagenesis followed by phenotypic measurements. For example, Novella et al. (2013) found that evolved vesicular stomatitis virus strains gained robustness, measured as resistance to mutagenesis, after laboratory adaptation. In contrast, Butković et al. (2020) found that a strain of turnip mosaic potyvirus evolved in *Arabidopsis thaliana* lost robustness over time, measured as the change in the ability of the virus to retain its level of plant infectivity after mutagenesis. Our barcode-based mutagenesis system makes it possible to dissect these overall changes in robustness, by measuring how they emerge as a result of changes in the effects of individual mutations.

In this study, we aim to leverage this mutation-level data to better understand the structure of epistasis in evolving populations. First, we analyze the overall distribution of fitness effects to measure how mutational robustness changes during adaptation in each environment. Next, we ask whether the distribution-level changes we observe can be explained by patterns of fitness-correlated epistasis among individual mutations. Finally, we examine how the effects of these mutations change in each of the evolving populations, asking whether epistasis is more often driven by predictable adaptations common across populations or by specific mutations fixed in a single population.

## Results

### Changes in the distribution of fitness effects during evolution

We isolated two clones from each of six timepoints from six haploid populations evolved in rich media at a permissive temperature (YPD 30°C) and from six haploid populations evolved in a defined media environment at a high temperature (SC 37°C), a total of 144 clones. We measured the fitness of each of these clones in the environment to which they adapted, finding that fitness increases steadily through time in both the YPD 30°C and SC 37°C environments, and displays a general pattern of declining adaptability (Figure 1A).

**Figure 1:**
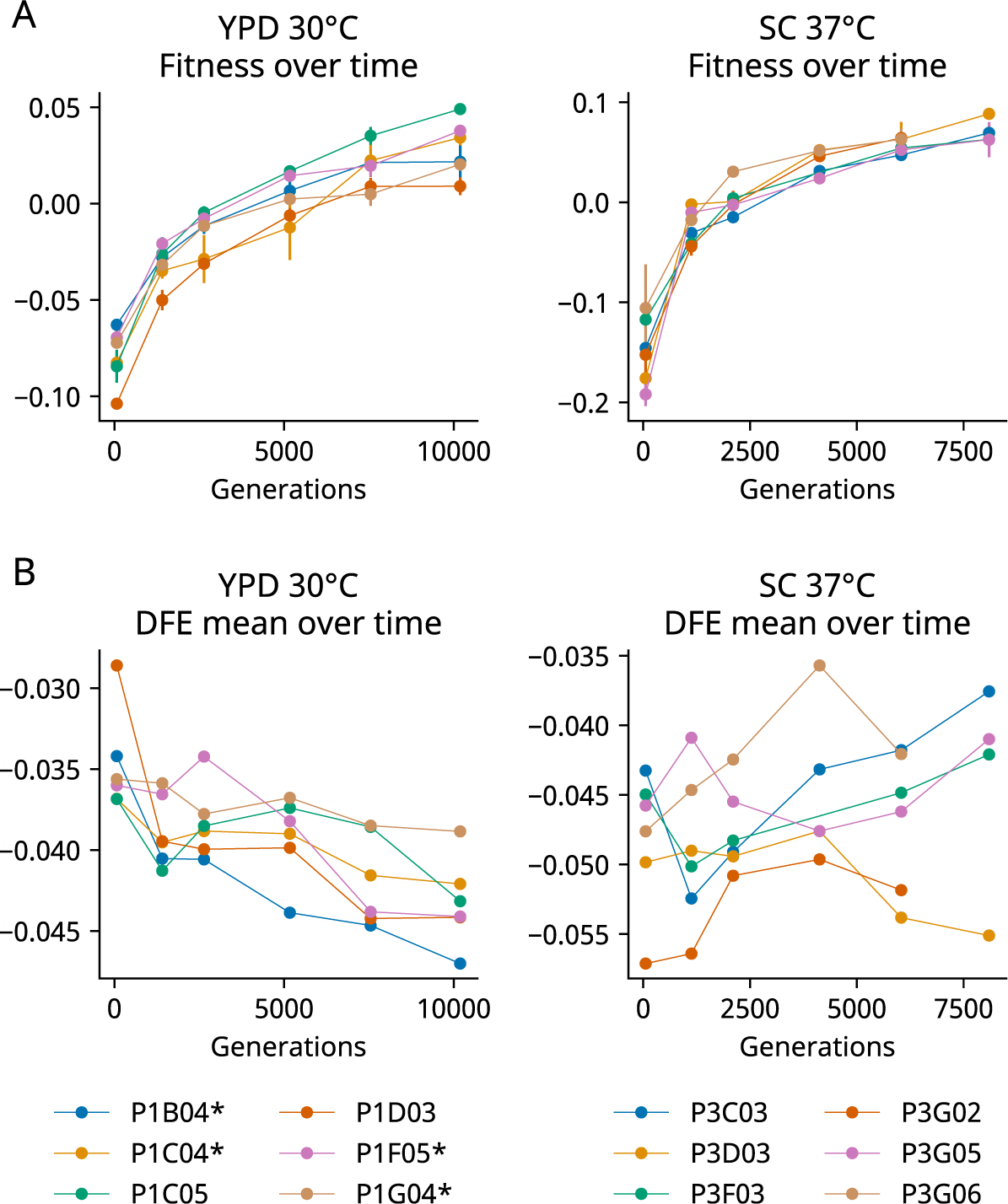
The DFE mean declines in one of two environments during evolution. **(A)** Changes in fitness during the evolution experiment, measured as the average fitness of two clones isolated from six timepoints in each population. In each graph, zero is the fitness of a fluorescent reference used in that environment. Error bars represent the standard deviation of the two replicates (points without error bars have errors smaller than the point size). **(B)** The mean fitness effect of the insertion mutations measured in the clones isolated from each timepoint. Asterisks in the legend represent a significant correlation (P < 0.05) between generation and DFE mean in that population alone. In both (A) and (B), the six populations are indicated by color.

We next created a library containing each of 91 insertion mutations in each of our 144 clones. We measured the fitness effect of each mutation in each clone using barcode-based competition assays as described in Johnson et al., 2019. While there are likely minor genetic differences between the two clones isolated from each population-timepoint that reflect intra-population diversity, we generally observed strong correlations between their fitness effect measurements (Figure 1 – figure supplements 1-3), so we averaged across the two clones to obtain one measure of the fitness effect of each mutation at each population-timepoint.

We find that the distribution of the fitness effects (DFE) of our 91 insertion mutations changes over the course of the evolution experiment. In the YPD 30°C environment, the mean of the DFE decreases over time as fitness increases during evolution (Figure 1B), consistent with a general pattern of both diminishing returns and increasing costs. The negative relationship between generations evolved and the mean of the DFE is significant in the entire set of population-timepoints (P = 2.0×10^-6^) and in four of our six individual populations (P < 0.05). In contrast, although the DFE does shift in individual populations evolved in our SC 37°C environment, we do not see any consistent patterns (Figure 1B).

In both environments, we find that the changes in the mean of the DFE are modest compared to our previous experiment (Johnson et al., 2019). The reasons for this are unclear, but may reflect the fact that fitness differences between the clones we study here are caused by a smaller number of mutations. We note that because of the modest differences in the DFE between clones, the noise in our measurements contributes significantly to the variation. One potentially biased source of this noise is missing measurements: not all mutations have fitness effects measured in each population-timepoint, due to differences in transformation efficiency or, most commonly, mutations that had too few read counts in the first two timepoints. As in Johnson et al., 2019, missing measurements are more common in more-fit strains (clones from later timepoints) in YPD 30°C, suggesting that the negative correlation between generations evolved and the mean of the DFE would be stronger with complete data (Figure 1 – figure supplement 5).

### Epistasis at the level of individual mutations

We next turn to the components of these distribution-level patterns: epistatic patterns for individual mutations. To look at these patterns, we focus on mutations that have fitness measurements in at least 20 population-timepoints in each environment (77 mutations in YPD 30°C, 74 in SC 37°C, 70 in both). We start by looking for patterns of fitness-correlated epistasis. We classify each mutation in each environment as correlated negatively or positively with background fitness if the correlation is significant at the P<0.05 level and the absolute value of the slope is greater than 0.05, and classify them as not significantly correlated if they do not meet these criteria. For both environments, we find numerous examples of both negative and positive correlations, corresponding to increasing costs or decreasing costs respectively for deleterious mutations (and to diminishing returns or increasing returns respectively for beneficial mutations).

In Figure 2, we show examples illustrating how the fitness effect of specific mutations vary across clones isolated from each environment, plotted as a function of the fitness of each clone in that environment (i.e. the background fitness). We find numerous examples of mutations which exhibit negative, positive, and non-significant correlations with background fitness in both environments (panels along the diagonal). We also find examples where a specific mutation exhibits a non-significant correlation in one environment and either a positive or negative correlation in the other (off-diagonal panels). Overall, consistent with our previous results, we observe more negative than positive correlations. As we would expect from the DFE-level results, this imbalance is more pronounced in the YPD 30°C environment: 33/77 mutations have fitness effects that decline significantly as background fitness increases in YPC 30°C, but only 17/74 in SC 37°C. We also find that only 9/77 mutations display the opposite pattern (fitness effects increase significantly as background fitness increases) in YPD 30°C, while 13/74 display this pattern in SC 37°C.

**Figure 2.**
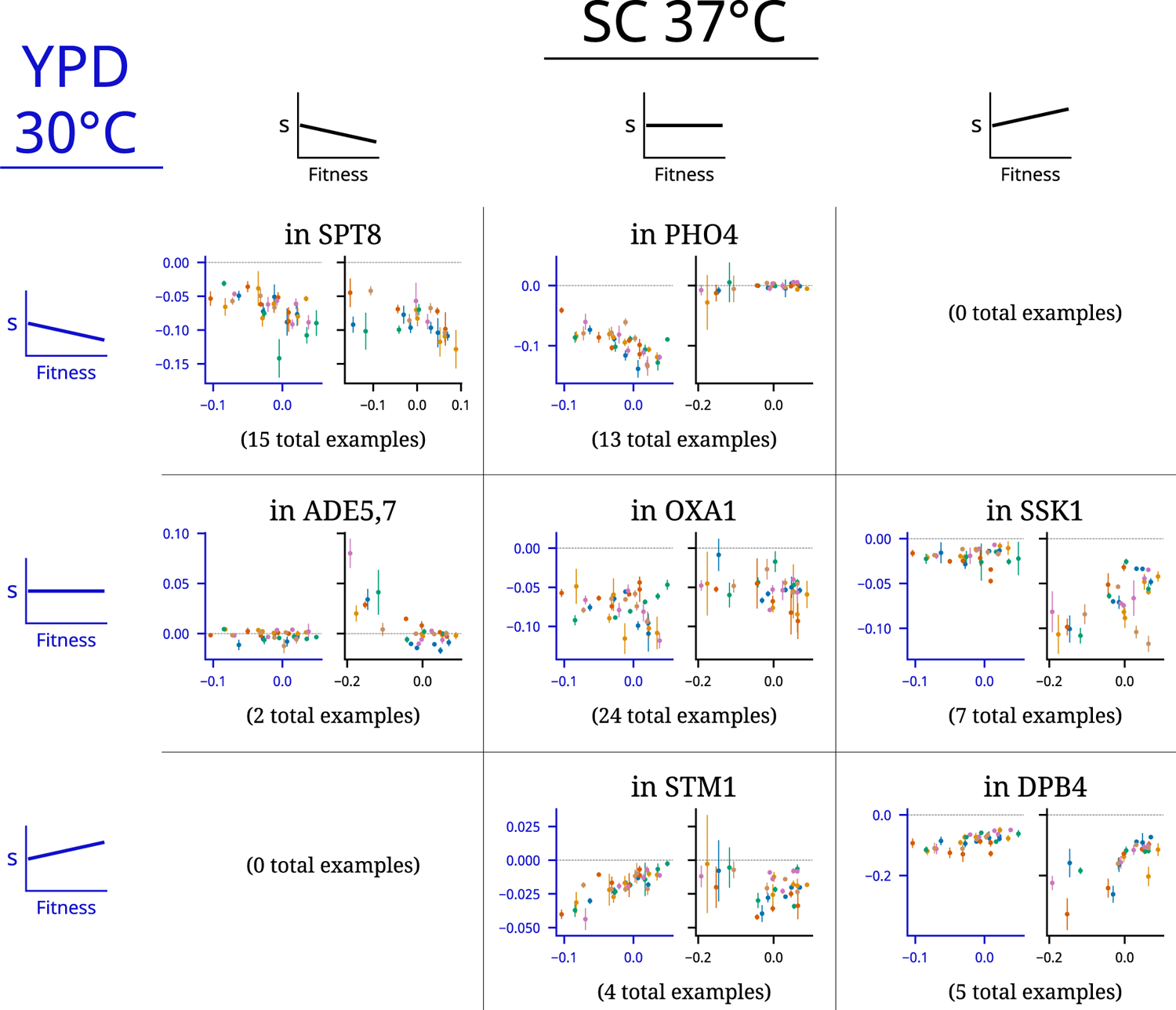
Patterns of fitness-correlated epistasis. Each panel shows an example of a specific mutation with a particular combination of relationships (negative, positive, or non-significant correlation between fitness effect of the mutation, *s,* and background fitness) in the two environments; numbers indicate the total number of mutations displaying each pair of relationships. Graphs depict fitness effect (y-axis) as a function of background fitness (x-axis). Axes are colored to identify the environment: in each panel the blue axes on the left is data from YPD 30°C and the black axes on the right is data from SC 37°C. Points are colored by population, as in Figure 1. Each set of example plots is labeled by where the mutation is in the genome (i.e. which gene it disrupts).

### Modeling the determinants of fitness effects

By definition, the fitness-correlated patterns we observe are the result of interactions between our insertion mutations and mutations that fix during the evolution experiment. If these interactions are all strictly “fitness-mediated,” the fitness effects of mutations will be fully explained by a background fitness effect. Alternately, fitness-correlated epistasis could arise based on the average effect of a number of idiosyncratic effects that are biased to produce a negative correlation with fitness. To understand the relative contribution of these determinants of epistasis, we compare 3 linear models used to explain the fitness effects of a single mutation in each of our two environments:

1. The fitness model (XM): the fitness effect of the mutation is a linear function of background fitness.
2. The idiosyncratic model (IM): the fitness effect of the mutation can change in any population at any timepoint when an interacting mutation fixes in that population (see below for our constraints on fitting these parameters).
3. The full model (FM): the fitness effect of the mutation is affected by both a linear effect of background fitness and the idiosyncratic interactions of fixed mutations, as in the idiosyncratic model.

We fit each model by ordinary least squares. We define the fitness effect of each mutation in the ancestral strain as the mean fitness effect measured among clones from the first timepoint, and fix the intercept of each of our models accordingly. For the IM and FM models, we add idiosyncratic parameters iteratively, choosing the parameter that improves the Bayesian Information Criteria (BIC) the most at each step. We do not allow parameters that fit a single point (e.g., a parameter for an effect at the final timepoint), and we only allow one parameter per population. We stop this iterative process of adding parameters if the BIC improves by less than 2 during a step (or when there is one parameter per population).

Figure 3A shows how well each model explains the fitness effect data for each mutation in each environment. We find that the fitness model often explains a large amount of variance, in agreement with our earlier analysis, but the idiosyncratic model and the full model usually offer more explanatory power. This indicates that epistasis is not strictly fitness-mediated; we commonly observe what appear to be stepwise changes in the fitness effect of a mutation unique to one population (Figure 3B), despite very similar patterns of fitness increase among populations (Figure 1A).

**Figure 3.**
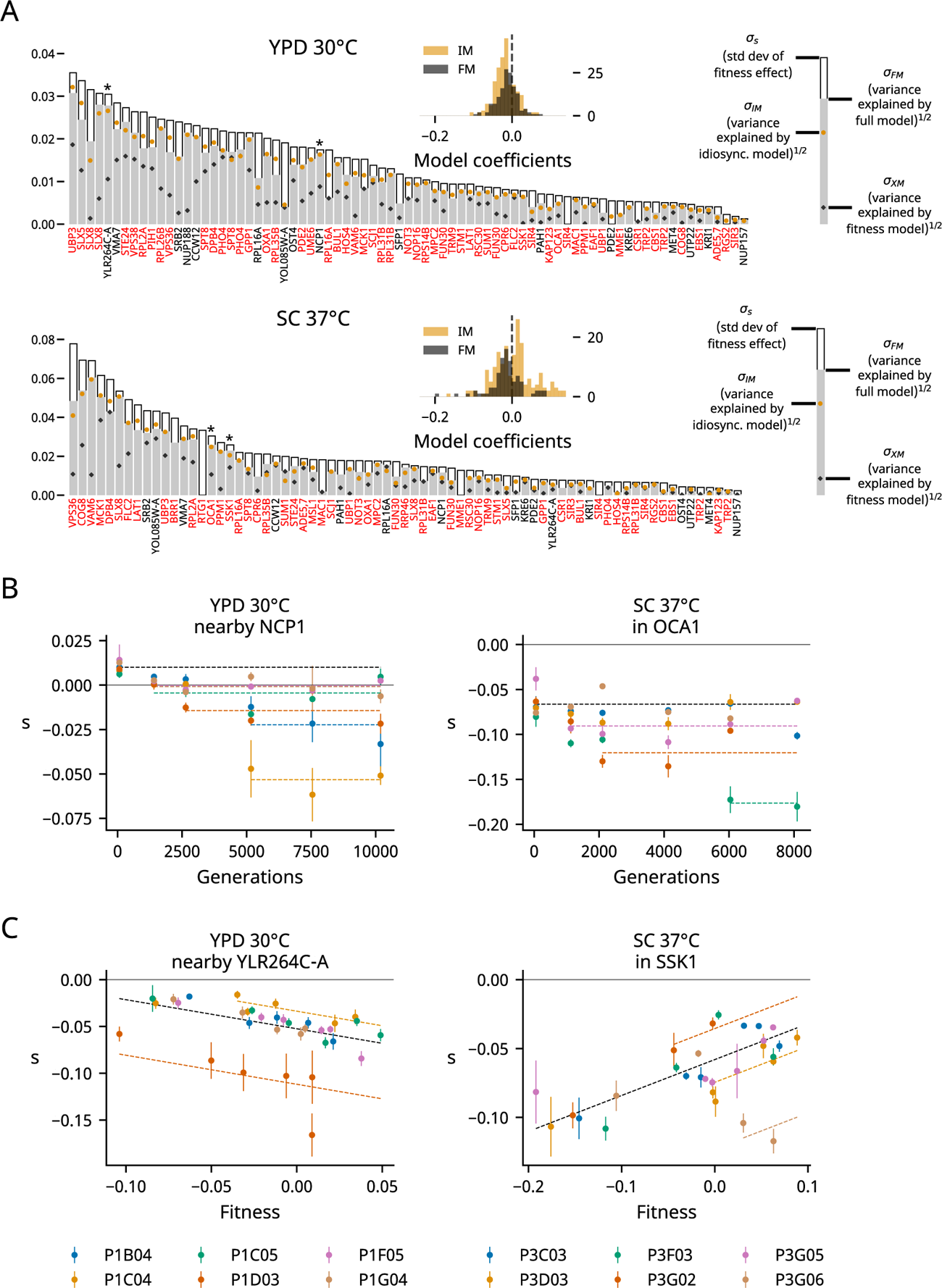
Determinants of fitness effects. **(A)** For each environment, we plot the standard deviation of the fitness effect across all population-timepoints and the square root of the variance explained by each of our three models. Mutations shown in red or black are insertions in or near the corresponding gene, respectively; stars indicate the mutations shown in panels B and C. Only mutations with fitness-effect measurements in at least 20 population-timepoints are shown. The insets show the distribution of all coefficients in the idiosyncratic model (IM) and full model (FM), pooled across all mutations. **(B)** Examples of idiosyncratic model fits. Model predictions are shown by dashed lines, and lines with contributions from indicator variables associated with a particular population are the same color as the points from that population (colors are the same as in Figure 1). **(C)** Examples of full model fits.

Positive and negative coefficients in the idiosyncratic model represent positive and negative epistasis between mutations that fix during evolution and our insertion mutations. The distribution of these coefficients in YPD 30°C is biased towards negative epistasis, while both positive and negative epistasis are common in our idiosyncratic model in SC 37°C (Figure 3A insets). Many of the positive epistatic terms in our idiosyncratic model in SC 37°C are the result of parallel reduction in the fitness costs of some deleterious mutations in the first 2,000 generations of evolution (e.g., see the mutation in SSK1 in Figure 3B and others in Figure 3 – figure supplements 4-5). In the full model, these parallel changes are often well explained by a background fitness parameter, and the remaining epistatic effects are more likely to be negative (Figure 3A, bottom inset).

The differences we observe in epistatic patterns could be caused by interactions between the environment and epistasis (“GxGxE” effects), differences in the adaptive targets in each environment, or a combination of the two. To tease apart these possibilities, we measured the fitness effects of mutations in clones evolved in YPD 30°C in the SC 37°C environment. We assayed the background fitness of each of these clones in SC 37°C and found that populations evolved in YPD 30°C sometimes experience large fitness declines in the SC 37°C environment (Figure 4C). We also observe widespread epistasis in this alternate environment, but do not observe any overall trends in the mean of the DFE in SC 37°C over the course of evolution in YPD 30°C. Instead, the variance in our data is dominated by one particularly low fitness clone in which several insertion mutations were strongly beneficial (Figure 4C).

**Figure 4.**
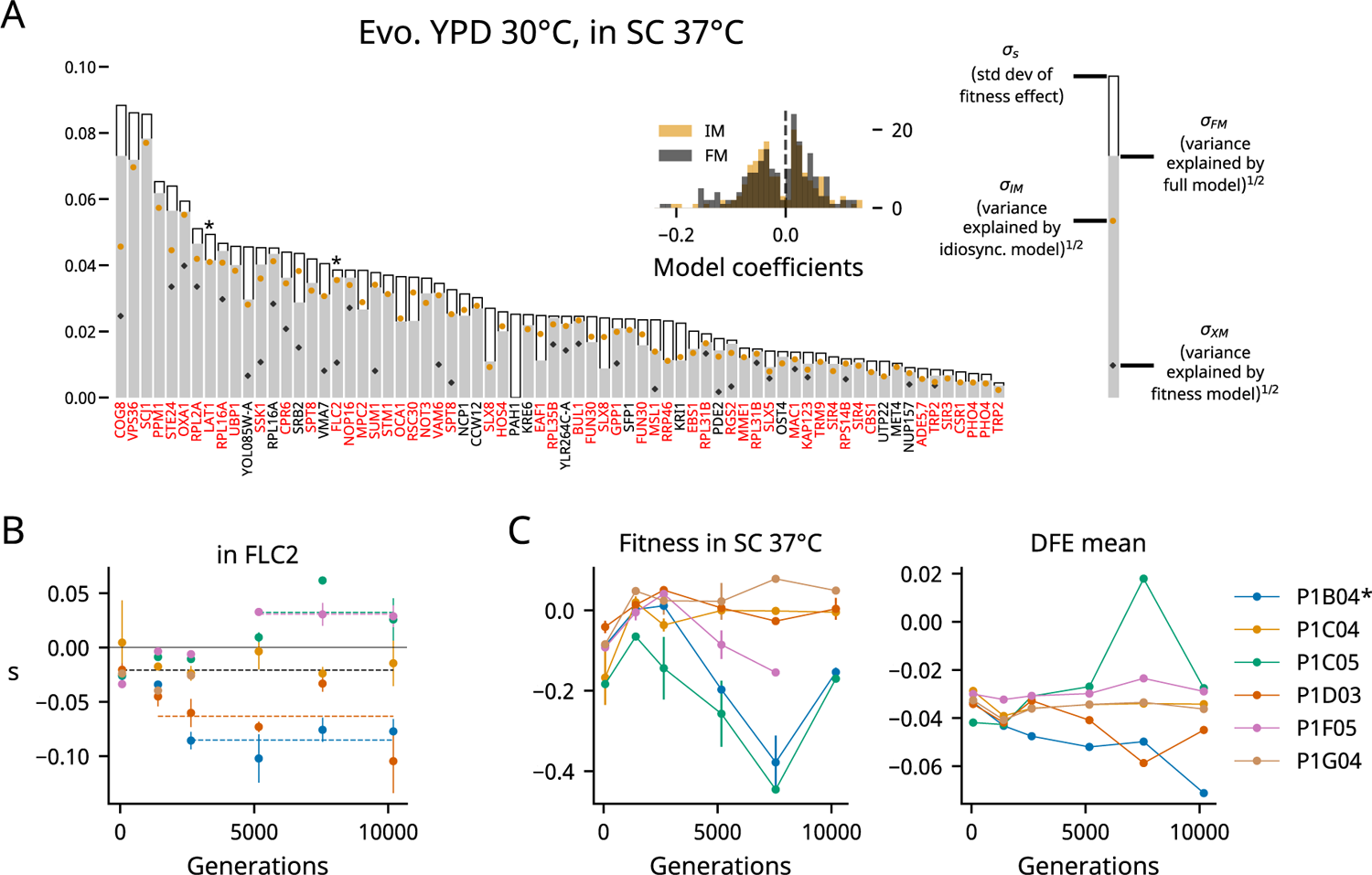
Patterns of epistasis in a non-evolution environment. **(A)** Same as Figure 3A, but for clones from YPD 30°C assayed in SC 37°C. We plot the standard deviation of the fitness effect across all population-timepoints and the square root of the variance explained by each of our three models. Mutations shown in red or black are insertions in or near the corresponding gene, respectively; stars indicate the mutations shown in panel B. Only mutations with fitness-effect measurements in at least 20 population-timepoints are shown. The inset shows the distribution of all coefficients in the idiosyncratic model (IM) and full model (FM), pooled across all mutations. **(B)** Example IM model fit, as in Figure 3B. The model predictions are shown by bold dashed lines, and lines with contributions from indicator variables associated with a particular population are the same color as the points from that population (colors are the same as in Figure 1 and panel C). **(C)** The fitness and DFE mean over time in YPD 30°C populations assayed in SC 37°C. The asterisk indicates a significant correlation (P<0.05). Error bars represent the standard deviation of the fitnesses measured for the two clones, but note that we were only able to measure the fitness of one clone at several population-timepoints due to low fitnesses relative to our reference; the corresponding points here have no error bars.

When we apply the same set of models to this dataset, we again find that both background fitness and idiosyncratic effects can explain how many mutations’ fitness effects vary, with the latter again outperforming the former (Figure 4A). Notably, we observe many more positive epistatic coefficients than in YPD 30°C, though we also observe a heavy tail of strongly negative coefficients. These patterns of idiosyncratic epistasis are not due to outsized contributions from a few populations; the distributions of IM coefficients for YPD 30°C populations are different in the two assay environments across populations (Figure 3 – figure supplement 2). Overall, these results support the hypothesis that GxGxE effects underlie the differences we observe between environments (see also Hall et al. 2019), though they do not rule out the possibility that differences in adaptive targets between environments may also contribute.

## Discussion

### Shifting distributions of fitness effects

By using our barcode-based mutagenesis system to assay the fitness effects of 91 specific gene disruption mutations across numerous genetic backgrounds spanning 8,000-10,000 generations of laboratory evolution, we have described how overall mutational robustness (defined in terms of the average effect of this type of insertion mutation) changes during evolution. We then dissected these overall effects in terms of the ways in which fitness effects of individual mutations change during evolution. We find that populations adapting to our YPD 30°C environment become less robust to deleterious mutations over time. This shift in the mean of the DFE in YPD 30°C is not caused by strictly fitness-mediated shifts in the fitness effects of mutations, but is instead the result of a bias towards negative idiosyncratic epistatic effects (Figure 3). In contrast, in clones isolated from populations evolved in SC 37°C, epistatic interactions are more evenly divided between negative and positive effects in our idiosyncratic model, though we did observe a bias towards negative idiosyncratic epistatic effects in a model that included a background fitness term.

The fact that populations evolved in YPD 30°C lost robustness as they increased in fitness over time is broadly consistent with our earlier work showing that the DFE becomes more strongly deleterious in more-fit genetic backgrounds (Johnson et al., 2019). However, the loss of robustness we observe here was not as strong as in this previous work (see Figure 1 – figure supplement 6 for a comparison to the effect size in Johnson et al., 2019), and we did not observe any predictable change in the DFE mean in SC 37°C. One potential explanation for the weaker patterns we observe at the DFE level in this experiment is that there are simply less mutations involved compared to the genetic backgrounds from our earlier work: the genotypes in our previous experiment were derived from a cross between two yeast strains that differed at tens of thousands of loci, so the pool of mutations was both larger and more balanced (each mutation present in ∼50% of clones) than in this study. If the overall patterns of fitness-correlated epistasis arise due to the collective effect of numerous idiosyncratic interactions, rather than a genuine fitness-mediated effect, we would therefore expect weaker trends here (Lyons et al., 2020; Reddy and Desai, 2021).

Our results also illustrate a form of hidden evolutionary unpredictability, despite the fact that our populations increased in fitness over time along a predictable trajectory (Johnson et al., 2021). As these populations adapted, they accumulated mutations that carry with them epistatic interactions with potential future mutations across the rest of the genome. The patterns of epistasis we observe demonstrate that the predictability of these second-order effects vary widely: some potential future mutations show strong fitness-correlated effects in every population, while others are affected by a small number of idiosyncratic interactions with mutations that fix during evolution. These unpredictable patterns of epistasis could lead to fundamentally unpredictable evolutionary outcomes, with changes in the fitness effects of mutations dynamically closing off or opening up evolutionary pathways during adaptation.

### Patterns of epistasis among individual mutations

Most of the insertion mutations we analyze in this experiment are deleterious across most or all genetic backgrounds. We find that these deleterious mutations tend to become more strongly deleterious over time in populations evolved in the YPD 30°C environment. This pattern results from an overabundance of negative epistatic interactions, which could involve either the beneficial mutations that drive fixation events or the neutral or weakly deleterious mutations that hitchhike to fixation during selective sweeps (McDonald et al., 2016). The strong predictive power of background fitness for the fitness effects of some mutations suggests that interactions with beneficial mutations are driving these patterns of epistasis, but idiosyncratic effects that deviate from these relationships hint at a role for interactions with hitchhiker mutations as well (Figure 3). Systematic backcrossing and mutagenesis experiments would be required to disentangle these patterns, but we suspect both types of interactions contribute to the epistasis we observe.

Among the relatively small number of beneficial insertion mutations we analyze, we find that the beneficial effects tend to decline over time, and almost universally shift to neutral or deleterious effects later in the experiment (Figure 3 – figure supplement 1). These results provide additional examples of diminishing-returns epistasis among beneficial mutations, which can at least partially explain the pattern of declining adaptability often observed in microbial evolution experiments (Chou et al., 2011; Khan et al., 2011; Kryazhimskiy et al., 2014; Wünsche et al., 2017). One of these mutations has a clear functional story behind it: the beneficial effect of an insertion mutation in ADE5,7 is the result of breaking the adenine synthesis pathway upstream of a toxic intermediate, so when populations in SC 37°C fix other loss-of-function mutations in this pathway between the first two sampled timepoints in the evolution experiment, this mutation becomes neutral (we do not see this effect in YPD 30°C because most populations have fixed a mutation in the adenine pathway by the first sampled timepoint (Johnson et al., 2021)).

The picture that emerges from our data is one in which idiosyncratic epistatic effects are largely unpredictable but have biases that often lead to correlations with background fitness. In both our own work and other studies of microbial evolution, these biases are more often towards negative epistasis: beneficial mutations tend to have negative epistatic interactions with both deleterious (Johnson et al., 2019, this study) and beneficial mutations (Chou et al., 2011; Hall et al., 2019; Karkare et al., 2021; Khan et al., 2011; Kryazhimskiy et al., 2014; Ono et al., 2017; Pearson et al., 2012; Perfeito et al., 2014; Rokyta et al., 2011; Wünsche et al., 2017).

What causes these biases in epistasis? At the broadest level, we can distinguish two sources of bias: the structure of biological systems and the set of mutations that fix during evolution. Both theoretical and experimental work have shown that genes within functional modules tend to have similar interaction profiles with other genes (Costanzo et al., 2016; Segrè et al., 2005). Given this kind of “monochromatic” epistasis, all that is necessary for fitness-correlated epistasis to appear during evolution is for beneficial mutations to be clustered in a few modules. The strength and direction of fitness-correlated epistasis will then depend on the particular modules targeted by selection and how those modules interact with the rest of the cell. For example, the targets of adaptation in SC 37°C may be more related to heat stress than general components of growth (e.g. mutations in LCB3, which have been shown to reduce killing in certain heat stress conditions, are enriched in populations in SC 37°C (Ferguson-Yankey et al., 2002; Johnson et al., 2021)). If the strongly deleterious effects of some of our insertion mutations are exacerbated by heat stress, but a beneficial mutation reduces heat stress, we can expect a positive interaction between the two mutations. In contrast, we hypothesize that the deleterious effects of some insertion mutations become more pronounced when growth rate is increased by mutations in YPD 30°C (Johnson et al., 2019). In order to understand or predict these differences in epistasis specific to the evolution environment, we will need to better understand the functional structure of biological systems.

### Higher-order epistasis and the evolution of robustness

Each mutation that fixes during evolution has an immediate first-order effect on fitness, but also carries with it a second-order set of pairwise interactions whose strength and direction is determined by the structure of functional relationships between genes and biological modules. Through higher-order interactions, a mutation can also change the structure of these functional relationships, altering the complexity, redundancy, robustness, and evolvability of biological systems (reviewed in de Visser et al., 2003, Masel and Siegal, 2009, Masel and Trotter, 2010, and Payne and Wagner, 2019). Our work here suggests that mutational robustness tends to decrease during evolution in some environments, but our data is essentially limited to second-order interactions. We expect these effects to dominate during rapid adaptation, but over longer evolutionary timescales, robustness may be more dependent on changes in the higher-order structure of biological systems.

### Ideas and speculation

We find that in one evolution environment, a bias towards negative epistasis leads to a shift in the DFE towards more deleterious mutations over time, while in another evolution environment no such shift occurs. Clearly, the details matter, and it is difficult to draw general conclusions. In this section, we speculate on how work in metabolic control theory (MCT) may help explain the functional underpinnings of our results and fitness-correlated epistasis more generally.

MCT describes how mutations in idealized metabolic pathways change the *control coefficients* of other mutations (Szathmáry, 1993). In a strictly serial pathway, a mutation reducing the activity of one enzyme will decrease the control other enzymes have over flux through the pathway, but a mutation increasing the activity of one of the enzymes will increase the control of others. If two enzymes act instead in parallel, these patterns are reversed: a mutation that increases activity of one will decrease the control of the other on flux, and vice versa.

If flux through the pathway is correlated with fitness, these patterns of interactions predict epistasis between mutations in different enzymes in the pathway. For example, if flux is *negatively* correlated with fitness in a strictly serial pathway, beneficial mutations will be those that reduce flux, so they will reduce the control of other enzymes in the pathway, such that these beneficial mutations exhibit negative epistasis. This is the case for beneficial loss-of-function mutations in the broken adenine-synthesis pathway present in the ancestors of our evolution experiment (Johnson et al., 2021): one beneficial loss-of-function mutation in this pathway (e.g. in ADE4) lowers the control coefficients of the rest of the enzymes in that pathway, such that loss-of-function mutations in these enzymes become less beneficial (e.g. our insertion mutation in ADE5,7 in this experiment, Figure 3 – figure supplement 1).

MCT is a mathematical framework for describing these interactions, but the general qualitative principles can be applied beyond enzymes in metabolic pathways to understand patterns of fitness-correlated epistasis (MacLean, 2010). One hypothesis for the functional underpinnings of increasing-costs epistasis can be framed in this way. First, we assert that the large-scale components of *growth* in the yeast cell *functionally* act in serial. While these components of growth (e.g. cell wall production, ribosome production, and DNA replication) do not belong to an actual serial pathway, they clearly do not act in parallel: these components generally cannot “fill in” for each other. That is, they are *non-redundant*. We therefore propose that an increase in the function of one of these components (in terms of growth rate) will increase the control coefficients of the rest of the components. Similarly, a decrease in the function of one component will decrease the control coefficients of the rest. We can intuitively understand this based on the idea of limitation: if DNA replication slows to a halt, growth rate will become less sensitive to changes in the speed of cell wall production. In contrast, if a population in which growth is limited by DNA replication fixes a mutation that improves DNA replication, we expect the control coefficient of cell wall production to increase, meaning deleterious mutations that slow cell well production will become more deleterious. We believe this effect can explain much of the increasing-costs epistasis we have observed.

Consider the following metaphor for this effect. You work for a car manufacturer, and your factory’s goal is to produce cars as quickly as possible. You work with a small team that builds the wheels. Your team is excellent, but the engine team is much slower. Because the engine team is limiting production, you don’t feel under pressure from the boss at all – honestly, you could skip work one week and the company would hardly suffer (read: your control coefficient is low, deleterious mutations have small effects). One day the engine team purchases a new robot and dramatically speeds up their process. Suddenly cars are waiting for wheels, and the pressure on your team increases dramatically – you can no longer slack off (read: your control coefficient is high, deleterious mutations have larger effects, costs have increased).

In our discussion, we speculated that we see increasing costs more frequently in YPD 30°C because adaptation in that environment is more focused on improving core components of growth, compared to adaptation in SC 37°C where selection for improvement in heat tolerance or survival may be more common. The phenotype of survival does not fit as neatly into our car manufacturing metaphor: we have no strong hypotheses for how control coefficients should change as populations increase heat tolerance, or for how large-scale phenotypes such as growth and survival integrate in terms of competitive fitness. However, we speculate that in SC 37°C, adaptive mutations are less likely to be affecting the core components of growth, and therefore cause increasing costs less often than mutations selected in YPD 30°C. To move closer to predictive models of epistasis, we will need to better understand the functional relationships between these large-scale cellular phenotypes.

We ended our discussion by noting that changes in robustness over longer evolutionary timescales may depend not on the type of changes we observe in our evolution experiment, but on changes to the higher-order structure of biological systems. Within our metaphor, this is the difference between changes to each team’s effectiveness and changes to the overall assembly line system. Defining what constitutes a change to the structure of the system itself is a difficult problem, analogous to defining “novelty” in evolution (Murray, 2020), but most would agree that there are fundamental differences in the complexity and redundancy of the genetic systems of viruses and humans, for example.

Both experimental and theoretical work have suggested that higher genome complexity and redundancy is associated with more negative epistasis between deleterious mutations (Macía et al., 2012; Sanjuán and Elena, 2006; Sanjuán and Nebot, 2008; though see Agrawal and Whitlock, 2010). Negative epistasis between deleterious mutations implies that deleterious mutations increase the control coefficients of other mutations and that beneficial mutations will decrease control coefficients. Therefore, in organisms with higher functional redundancy, we expect to observe more positive interactions between beneficial and deleterious mutations and less increasing-costs epistasis. In other words, the second-order loss of robustness we observe during adaptation should be stronger in organisms that are already less robust. While there may be more unseen biases in the types of interactions beneficial mutations participate in, present data suggests that increasing-costs epistasis may be specific to organisms or environments where the functional redundancy of genes or biological modules is low.

### Epistasis between beneficial mutations

An apparent contradiction emerges from our explanation for increasing-costs epistasis: if adaptation increases the control coefficients of core components of growth, why do we not see more positive epistasis between beneficial mutations in evolution experiments? Why do we instead usually see negative, diminishing-returns epistasis and declining adaptability? We propose that this discrepancy arises due to differences in the availability and form of beneficial and deleterious mutations. Based on previous work, we expect beneficial mutations in laboratory evolution experiments to be primarily loss-of-function mutations (Murray, 2020). In contrast to deleterious mutations, which can be spread across the genome, these types of beneficial mutations will rarely exist in well-adapted core components of growth, and will instead be clustered in a few adaptive targets. Within these targets, we believe loss-of-function beneficial mutations are often *functionally redundant*, meaning that they tend to *decrease* each other’s control coefficients for fitness. Beneficial mutations can be redundant by inactivating the same deleterious pathway (e.g., ADE pathway mutations discussed above), solving the same general problem (e.g., mutations shortening lag in Karkare et al., 2021), or changing a phenotype with a nonlinear fitness function (Chiu et al., 2012; Chou et al., 2014; Keren et al., 2016; Lunzer et al., 2005; Otwinowski et al., 2018).

Nonmonotonic fitness functions can arise from phenotypes with both potential benefits and costs (Dekel and Alon, 2005), such that negative interactions between beneficial mutations and the benefit can lead to fitness-correlated epistasis that crosses neutrality, exhibiting diminishing returns, increasing costs, and sign epistasis (Figure 3 – figure supplement 1).

This explanation provides a prediction: we will be more likely to see synergistic epistasis during evolution experiments when we observe beneficial *gain-of-function* mutations. Chou et al., 2009 provides a particularly strong example of a beneficial gain-of-function (promoter capture) mutation that is more beneficial in more-fit genetic backgrounds. In the long-term *Escherichia coli* evolution experiment, potentiating mutations acquired during adaptation in one population interacted positively with a beneficial gain-of-function mutation (also a promoter capture), enabling aerobic citrate utilization (Blount et al., 2012). Studies of evolutionary repair also provide examples of synergistic interactions between apparently non-redundant beneficial mutations (Fumasoni and Murray, 2020; Hsieh et al., 2020). These counter examples underscore the fact that diminishing returns epistasis is not a rule; it is a pattern that is overrepresented in evolution experiments due to biases for beneficial mutations to be loss-of-function mutations and to be clustered in a few adaptive targets. These biases may be weaker later in evolution experiments when mutations are spread more evenly across cellular modules, such that a period of declining adaptability caused by diminishing returns epistasis early in an experiment gives way to a period of relatively constant fitness gains (Good and Desai, 2015).

### A final note on terminology for epistasis

Very few papers discuss epistasis between beneficial and deleterious mutations – most theoretical and experimental work has focused on epistasis between two beneficial mutations or two deleterious mutations. With these same-signed pairs of mutations, the terms negative and positive epistasis are consistent. Two beneficial mutations that interact negatively imply two deleterious reversions that also interact negatively. However, when we consider one of these beneficial mutations and the deleterious reversion of the other, they interact positively. We provide this note to clarify that increasing costs epistasis, in which deleterious mutations exhibit negative epistasis with beneficial mutations, should not be associated with previous results demonstrating negative epistasis – instead we should expect it in systems where more positive epistasis is observed. While epistasis is already a concept overladen with terminology, we submit that in some cases it may be more useful to classify interactions between mutations or cellular components as being functionally redundant or non-redundant in terms of fitness.

## Materials and Methods

### Strains

All strains used for this study were isolated from the evolution experiment described in Johnson et al., 2021. We isolated 2 clones from each of our focal populations at each sequencing timepoint. For this experiment, we used clones from 12 MATa populations, 6 from the YPD 30°C environment and 6 from the SC 37°C environment. We decided to include population P1B04, which exhibits a cell-clumping phenotype in preliminary imaging data, and to exclude population P1B03, which diploidized during evolution, and populations P3C04, P3F05, P3D05, and P3E02, which lost G-418 resistance during evolution (not being able to select on G-418 during transformation could allow the HygMX cassette to replace the KanMX cassette, leading to leakage during the selection step). Otherwise we chose populations randomly. The ancestor of these populations is MJM361 (MATa, YCR043C:KanMX, STE5pr-URA3, ade2-1, his3Δ::3xHA, leu2Δ::3xHA, trp1-1, can1::STE2pr-HIS3 STE3pr-LEU2, HML::NATMX, rad5-535).

### Barcoded Tn7 libraries

We used a previously created set of Tn7-based plasmid libraries to introduce the same set of ∼100 mutations into each of our strains (Johnson et al., 2019). These plasmids contain a section of the yeast genome corresponding to one of these ∼100 locations, interrupted by a Tn7 insertion containing a random DNA barcode and a HygMX cassette. Each barcode uniquely identifies the mutation and the plasmid library via a mapping established in earlier work (Johnson et al., 2019).

### Transformation

Our yeast transformation protocol is a scaled-up version of that used in Johnson et al. 2019, based on the method described in (Gietz and Schiestl, 2007). We grew strains from freezer stocks overnight, diluted 750 ul into 15 mL YPD + Ampicillin (100 g/mL), grew for 4 hours, pelleted the cells, and resuspended in 900 uL transformation mix and 100 uL plasmid DNA cut with NotI-HF (corresponds to 2 ug of plasmid; cut at 37°C for 3 hours, then heat inactivated at 65°C for 10 min). We then heat shocked this mixture at 42°C for 1 hour, recovered in 3 mL YPD + Ampicillin for 2 hours, plated 25 ul on antibiotic selection plates to check efficiency, and then combined the rest with 40 mL YPD supplemented with antibiotics. For both agar and liquid selective media we included Hygromycin (300 μg/ml), clonNat (20 μg/ml), and G-418 (200 μg/ml). We made frozen glycerol stocks of each transformation after ∼48 hours of growth. All growth was conducted at 30°C, either in a test tube on a roller drum (recovery), or in a baffled flask on an orbital shaker (all other steps).

We transformed 2 clones from 12 populations at 6 timepoints for a total of 144 transformations. We organized these transformations into three “VTn assays,” each associated with 48 transformations using our 48 unique barcoded libraries.

### Fitness Assays

Again, we followed the protocols established in Johnson et al., 2019 for our fitness assays. We assayed our transformed libraries of clones from the YPD 30°C environment in both their evolution environment (YPD 30°C) and the SC 37°C environment, and clones from the SC 37°C environment in their evolution environment (SC 37°C). We first arrayed our transformation glycerol stocks into 2 96-well plates corresponding to the 2 evolution environments, and then inoculated 8 ul from each well of these plates into 4 (for SC 37°C assays) or 8 (for YPD 30°C assays) flat-bottom polypropylene 96-well plates containing 126 ul of media, supplemented with the same antibiotics as during the initial selection. To ensure efficacy of the antibiotics in the SC 37°C environment, we used media with MSG instead of ammonium sulfate (1.71 g/L YNB without amino acids or ammonium sulfate, 2 g/L SC, 1 g/L MSG). After this period of growth, we used YPD and SC supplemented with ampicillin (100 μg/ml) and tetracycline (25 μg/ml), matching the conditions of the evolution experiment. After 40 hours of growth in these plates, we started daily transfers.

At each daily transfer, we diluted YPD 30°C cultures 1/2^10^ and SC 37°C cultures 1/2^8^. During these transfers, we combine and mix cultures from each well corresponding to the same clone/transformation to increase population size and reduce bottleneck noise. In the first (T0) transfer, we combined cultures from the 8 plates that were initially inoculated from the freezer stock and diluted them into 20 96-well plates. In all subsequent transfers (T1-4), we combined cultures from all 20 plates and diluted them into 20 new plates. Specifically, for YPD 30°C, we diluted 3 μl from each well of 20 plates into 60 ul YPD (60 μl total, 1/2 dilution), mixed, then diluted 16 μl into 112 ul YPD (1/2^3^ dilution), mixed, and distributed 2 μl into 126 μl YPD in 20 plates (1/2^6^ dilution). For SC 37°C, we diluted 3 μl from each well of 20 plates into 60 ul SC (60 μl total, 1/2 dilution), mixed, then diluted 60 μl into 60 ul SC (1/2 dilution), mixed, and distributed 2 μl into 126 μl SC in 20 plates (1/2^6^ dilution).

### Barcode sequencing

Our fitness assays in YPD 30°C were originally performed alongside assays in SC 37°C that were later abandoned due to an issue with expired reagents (and repeated with appropriate reagents in our second round of assays). During these assays, we combined equal volumes of culture at the end of each transfer from every well corresponding to each of the three VTn assays in each environment. We can pool the cultures corresponding to each VTn assay because we know which barcodes correspond to which plasmid library/clone, so we can divide our barcode count data appropriately during sequencing analysis. We performed DNA extractions from two 1.5 ml pellets for each assay-timepoint from our YPD 30°C fitness assays and from four 1.5 ml pellets from our SC 37°C fitness assays using Protocol I from the Yeastar Genomic DNA Kit (Zymo Research), as described previously (Johnson et al., 2019). We then amplified barcodes using a two-step PCR protocol. We performed 4 first round PCRs with 19 µl gDNA, 25 l 2X Kapa Hotstart Hifi MM, 3 µl 10 M TnRS1 primer, and 3 µl 10 M TnFX primer, and ran the PCR protocol: 1) 95°C 3:00, 2) 98°C 0:20, 3) 60°C 0:30, 4) 72°C 0:30, GO TO step 2 3 times, 5) 72°C 1:00. We purified these PCRs with PCRClean DX Magnetic Beads (Aline), using a 0.85X ratio. We then set up two second round PCRs per sample by combining 25 µl purified PCR µ1 product, 1.5 µl ddH2O, 10 µl Kapa Hifi Buffer, 1 µl KAPA HiFi HotStart DNA Polymerase, 5 µl 5 M N7XX primer (Nextera), and 5 µl 5 M S5XX primer (Nextera), and ran the PCR protocol: 1) 95°C 3:00, 2) 98°C 0:20, 3) 61°C 0:30, 4) 72°C 0:30, GO TO step 2 19 times, 5) 72°C 2:00. We purified the resulting libraries with Aline beads, using a 0.7X ratio, then repeated the purification with a 0.65X ratio, and finally sequenced our pooled libraries on a NextSeq 550 (Illumina).

### From reads to barcode counts

We process our sequencing data as described previously (Johnson et al., 2019). We first filter reads based on inline indices and quality scores, use regular expressions to extract barcode sequences, and combine barcode counts across timepoints for each VTn assay. Next, we use a single-bp-deletion-neighborhood method to correct errors in raw barcodes, assigning them to the set of known barcodes from each of our plasmid libraries. By associating barcodes with plasmid libraries, we associate them both with a fitness assay for a particular clone and with a particular insertion mutation, and we divide our barcode count data accordingly.

### Estimating fitness effects from barcode counts

Again, we follow Johnson et al., 2019, with minor differences. First, we convert barcode counts to log-frequencies at each timepoint. After this preliminary step, we noticed a large number of log-frequency spikes, restricted largely to one timepoint in one of our VTn assays in SC 37C. These spikes in frequency very likely represent low-level sequencing library contamination from another timepoint due to primer cross-contamination. In Figure 1 – figure supplement 6, we show that this is confined to this single timepoint and demonstrate how we can use a simple heuristic (excluding lineages whose log-frequency at timepoint 2 is 0.5 greater than both timepoint 1 and timepoint 3) to remove the barcoded lineages affected by this sequencing library contamination. This step excludes, on average, less than 1% of the reads from timepoint 2 in these assays. Next, we calculate fitness effects for each barcode, as described in Johnson et al., 2019. After excluding timepoints with less than 5,000 total barcode counts, we measure the log-frequency slope for each barcode at each consecutive pair of timepoints, excluding timepoints in which the barcode has less than 10 counts. We scale each of these log-frequency slopes by the median log-frequency slope of barcodes associated with five neutral reference mutations, and then average these scaled values to get one fitness measurement for each barcode. As in Johnson et al., 2019, we observe a small fraction of outlier barcodes, which follow starkly different log-frequency trajectories than the other barcodes associated with the same insertion mutation, presumably due either to pre-existing mutations in the transformed culture or transformation artifacts (including two mutations being transformed together). We use a log-likelihood ratio test to identify barcodes whose read counts are inconsistent with barcodes near the median fitness measured for one insertion mutation. Based on iterative exclusion and exploration of frequency trajectories for this experiment, we chose a heuristic cutoff of 40 for the log-likelihood ratio required to exclude barcodes (for a detailed description of this method see Johnson et al., 2019). Finally, to decrease noise from low-frequency barcode lineages while retaining the independent measurements unique barcodes provide, we randomly combine counts from individual barcodes into a maximum of 5 combined barcodes (“cBCs”) per insertion mutation. Next, we repeat the fitness measurement process described above to get final fitness measurements for each cBC.

When we compare the average fitness effect measurement for each insertion mutation between the two clones isolated from each population at each timepoint, we see strong agreement (Figure 1 – figure supplements 1-3). Given these strong correlations, we compute fitness effects for each population-timepoint using measurements from cBCs from both clones, treating them as we treated biological replicates in Johnson et al., 2019. We compute the fitness effect (*s*) and the standard error of *s* for each mutation as described in Johnson et al., 2019.

### Measuring changes in the DFE and accounting for missing measurements

For each population-timepoint, we calculated the mean of the fitness effects of all mutations with at least 3 cBCs across the two clones, excluding population-timepoints with less than 20 mutations with fitness effect measurements. To account for missing fitness effect measurements, we first examined the number of deleterious mutations (defined as having a mean fitness effect < −0.05 across all strains in a given condition) that were not measured in each population-timepoint. We next examined the DFE mean of sets of mutations shared across the set of population-timepoints we had assayed successfully in each condition. Finally, we created “filled-in” DFEs for each population timepoint in which missing measurements were replaced with the mean fitness effect measured across all strains in a given condition. The results of these analyses are plotted in Figure 1 – figure supplement 5.

### Modeling the determinants of epistasis

Because our modeling approach can be strongly influenced by outliers, we only consider fitness effect measurements for mutations with at least 5 cBCs across the two clones for these analyses. We describe our basic modeling approach in the main text. To fix the intercepts of our models correctly, we first transform the fitness effect of each mutation by subtracting the mean fitness effect measured across all populations at the first timepoint of the evolution experiment. We similarly transform our background fitness variable by subtracting the average fitness measurement across all populations at the first timepoint. Then we fix the intercept at (0, 0). Note that our plots showing these model fits use the natural scales for fitness and fitness effects, not these transformed scales. All model fitting was performed using the python package statsmodels (Seabold and Perktold, 2010).

### Measuring background fitness

We measured the background fitness of clones with fluorescence-based competitive fitness assays in duplicate for each clone in each environment, using the reference strains strain 2490A-GFP1 and 11470A-GFP1 for the YPD 30°C and SC 37°C clones, respectively. We used the 2490A-GFP1 reference when we assayed the YPD-30°C-evolved clones in SC 37°C because come of these clones have very low fitness and 2490A-GFP1 has a lower fitness than 11470A-GFP1. We used the fitness difference measured between these two references in Johnson et al., 2021 to standardize the fitness measurements YPD-30°C-evolved clones in SC 37°C so that all fitness measurements in SC 37°C are on the same scale. Fitness assays were performed and data was analyzed as described in Johnson et al., 2021. Briefly, we maintained mixed cultures of our clones and fluorescent references for 3 daily growth cycles, as described above, and measured the frequency of fluorescent cells at each transfer using flow cytometry. We then calculated the fitness of each clone as the slope of the natural log of the ratio between the frequencies of the non-reference and reference cell populations over time.

## Acknowledgements

We thank Sergey Kryazhimskiy, Alena Martsul, Andrew Murray and members of the Desai lab for useful discussions about experimental design and analysis. We thank Shreyas Gopalakrishnan, Juhee Goyal, and Megan E. Dillingham for their help with isolating the clones used in this experiment. This work was supported by an NSF Graduate Research Fellowship (to M.S.J.), the NSF (PHY-1914916), and the NIH (GM104239). Computational work was performed on the Cannon cluster supported by the Research Computing Group at Harvard University.

## Data and code accessibility

Raw sequencing data has been deposited in the GenBank SRA (accession: SRP351176). All code used in this project is available on GitHub (https://github.com/mjohnson11/VTn_pipeline).

## Additional files

### Supplementary files

Supplementary File 1. Column-annotated underlying data for this project. Includes background fitness, fitness effect, and modeling data from this experiment and Johnson et al., 2019.

Supplementary File 2. Oligos used in this study.

## Figure supplements

**Figure 1 – figure supplement 1.**
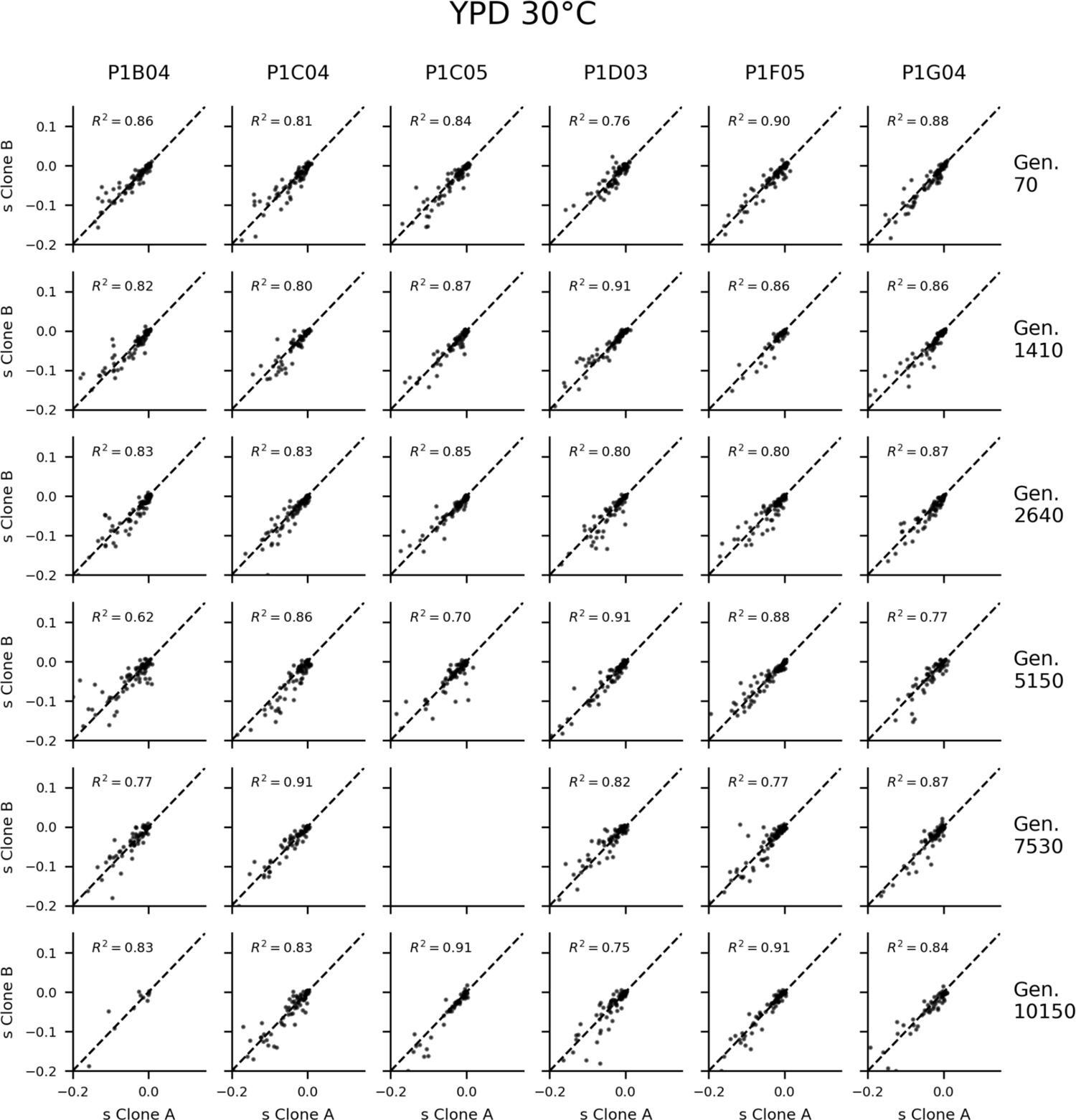
Fitness effect measurement correlations in YPD 30°C. Only mutations with at least three barcodes with fitness effect measurements in each clone are included. Values are graphically pinned to the −0.2 and 0.15 if they fall outside that range.

**Figure 1 – figure supplement 2.**
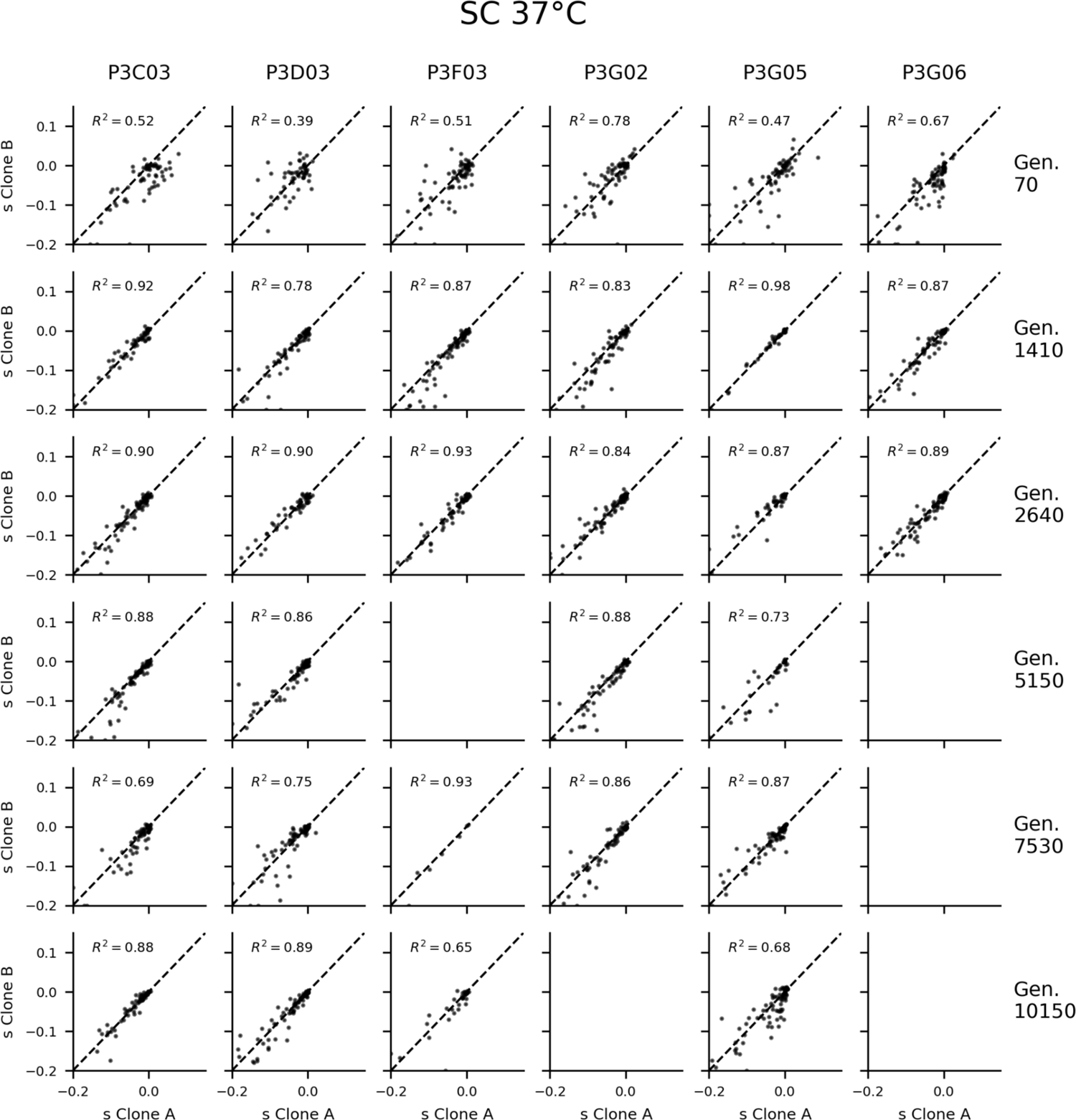
Fitness effect measurement correlations in SC 37°C. Only mutations with at least three barcodes with fitness effect measurements in each clone are included. Values are graphically pinned to the −0.2 and 0.15 if they fall outside that range.

**Figure 1 – figure supplement 3.**
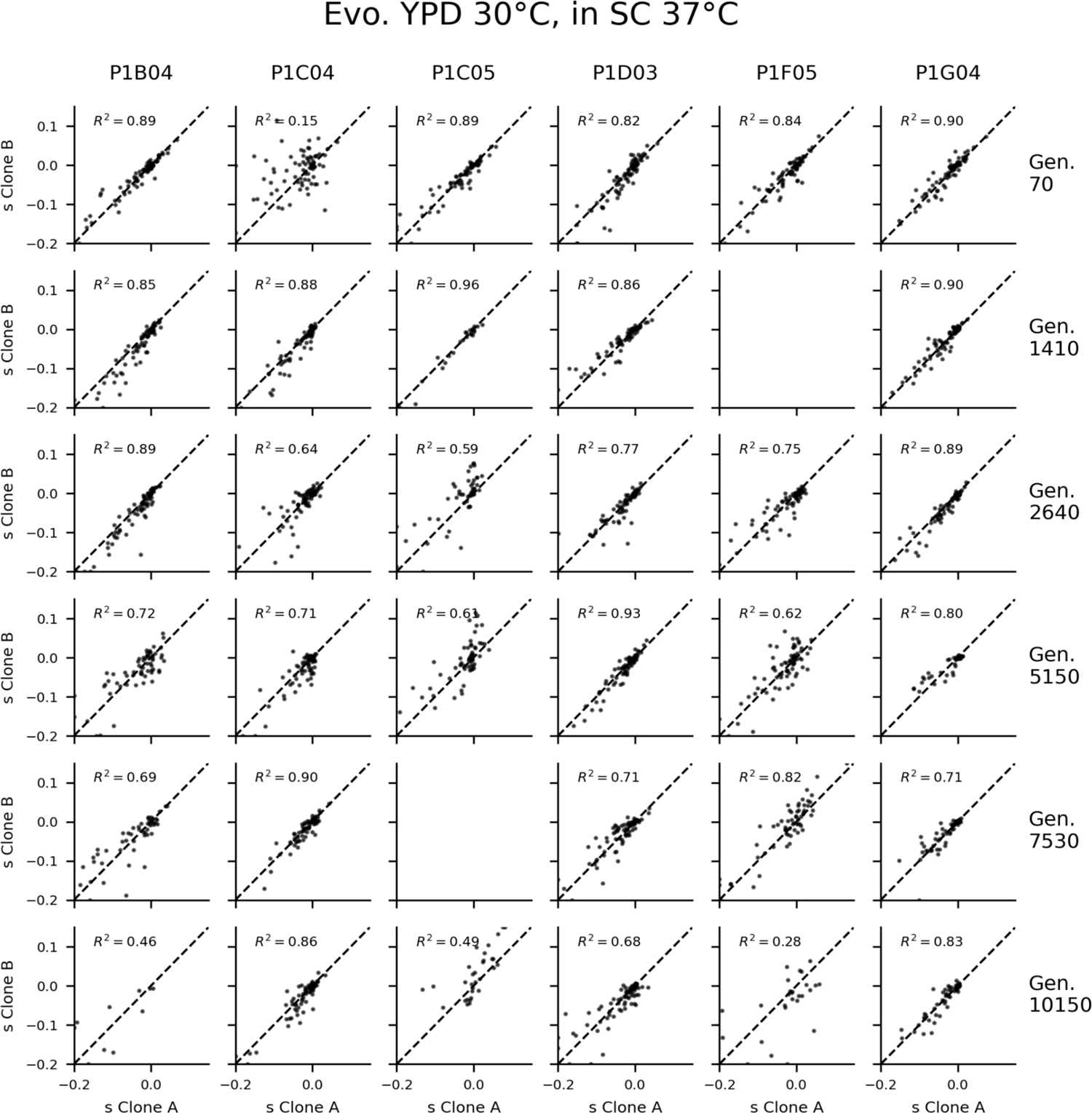
Fitness effect measurement correlations in clones evolved in YPD 30°C, assayed in SC 37°C. Only mutations with at least three barcodes with fitness effect measurements in each clone are included. Values are graphically pinned to the −0.2 and 0.15 if they fall outside that range.

**Figure 1 – figure supplement 4.**
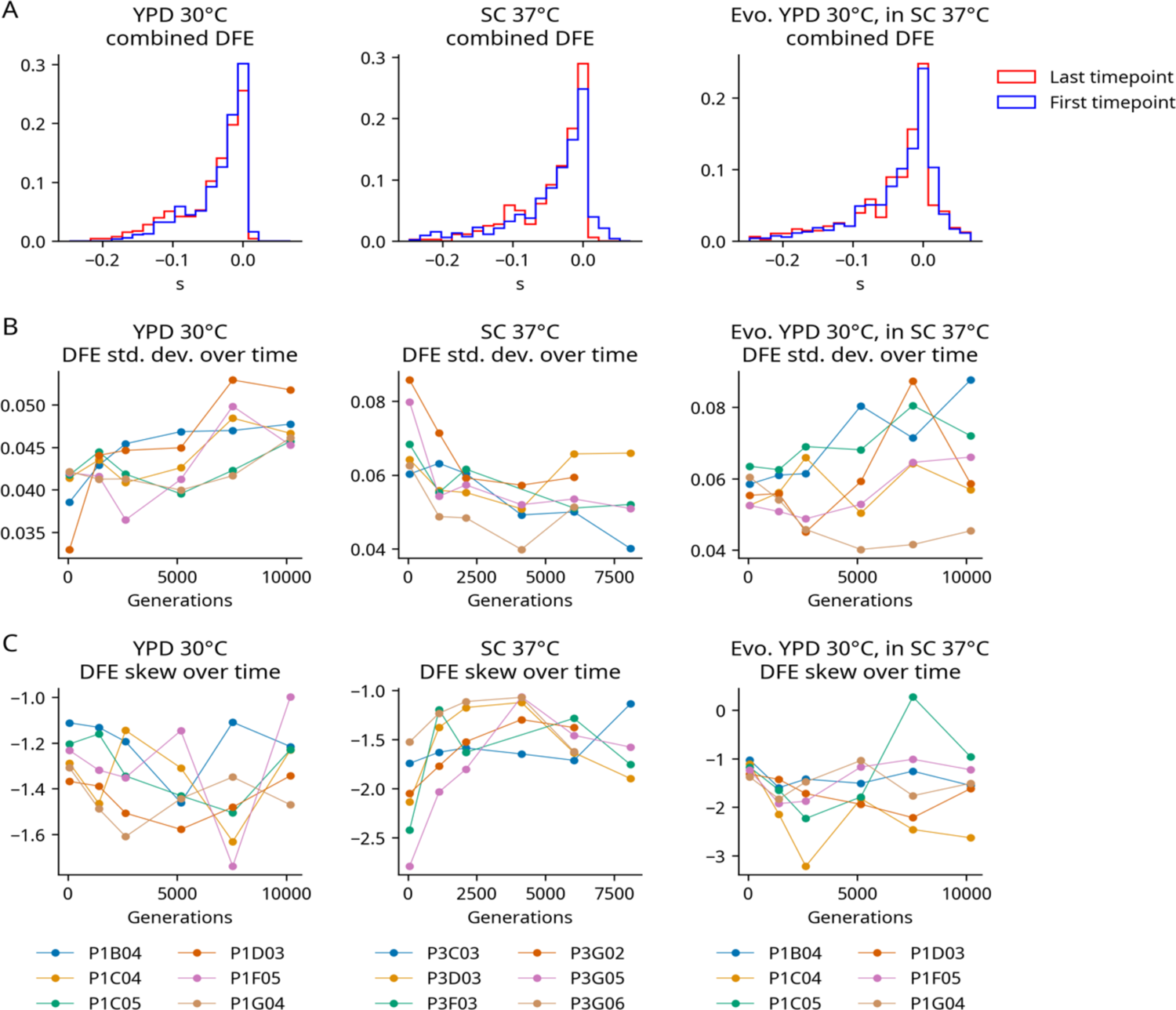
Additional DFE statistics. **(A)** The combined DFE of all populations at the first and last timepoint in each environment. **(B)** The standard deviation of the DFE over time in each population. **(C)** The skew of the DFE over time in each population.

**Figure 1 – figure supplement 5.**
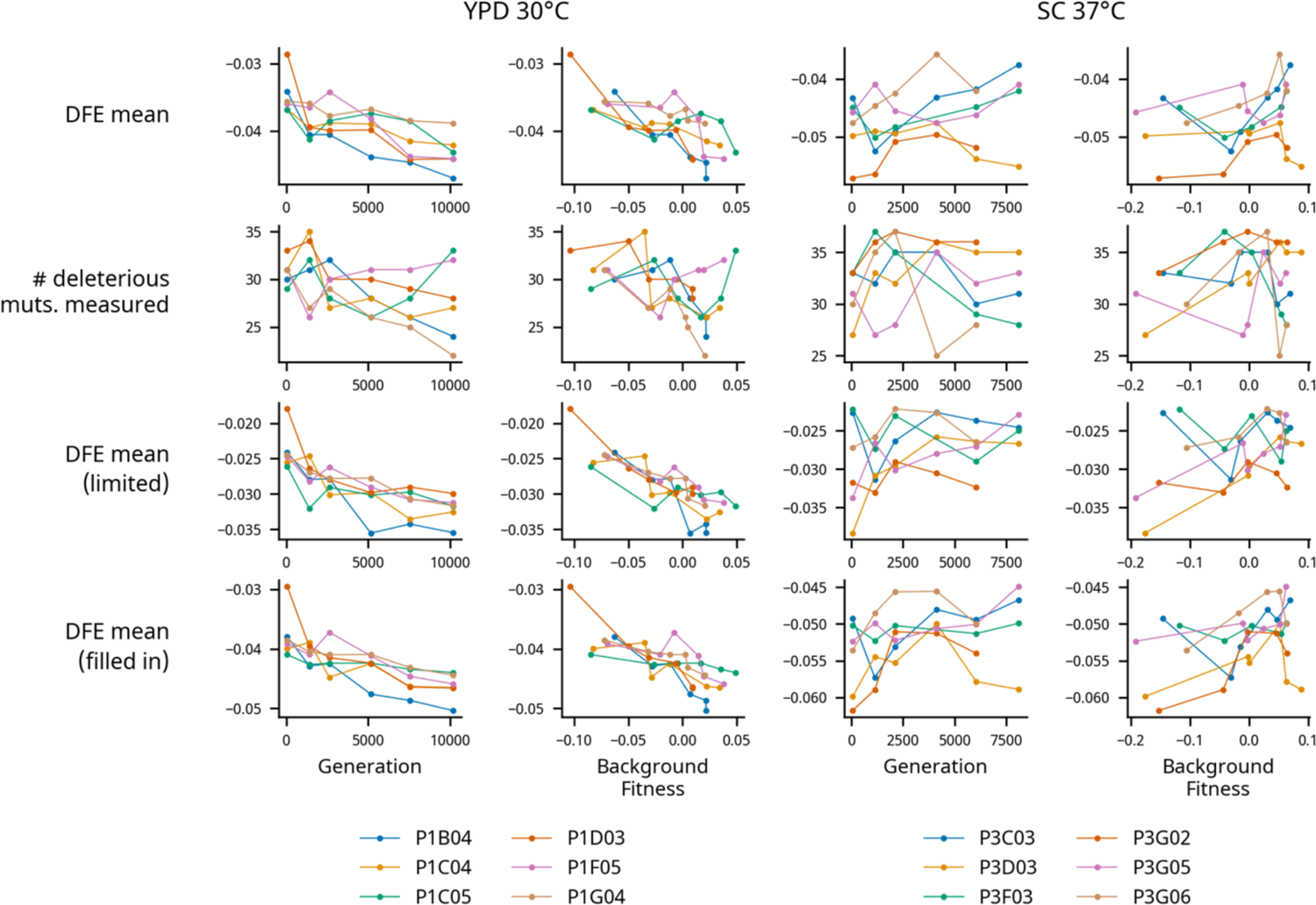
Accounting for missing fitness effect measurements. Relationships between generations evolved and background fitness and various DFE measurements in each environment. The first row shows the DFE mean. The second row shows the number of deleterious mutations (based on their average fitness effect across strains) that had their fitness effects successfully measured in each population-timepoint. The third row shows the mean of a DFE of mutations in which every mutation is measured in every population timepoint with usable data (68 mutations in all 36 population-timepoints successfully assayed in YPD 30°C, 45 mutations in 33 populations timepoints successfully assayed in SC 37°C). The fourth row shows the mean of a DFE in which missing fitness effect measurements are filled in with their average fitness effect across strains. These methods for accounting for missing fitness measurements strengthen our conclusion that the mean of the DFE declines during evolution in YPD 30°C.

**Figure 1 – figure supplement 6.**
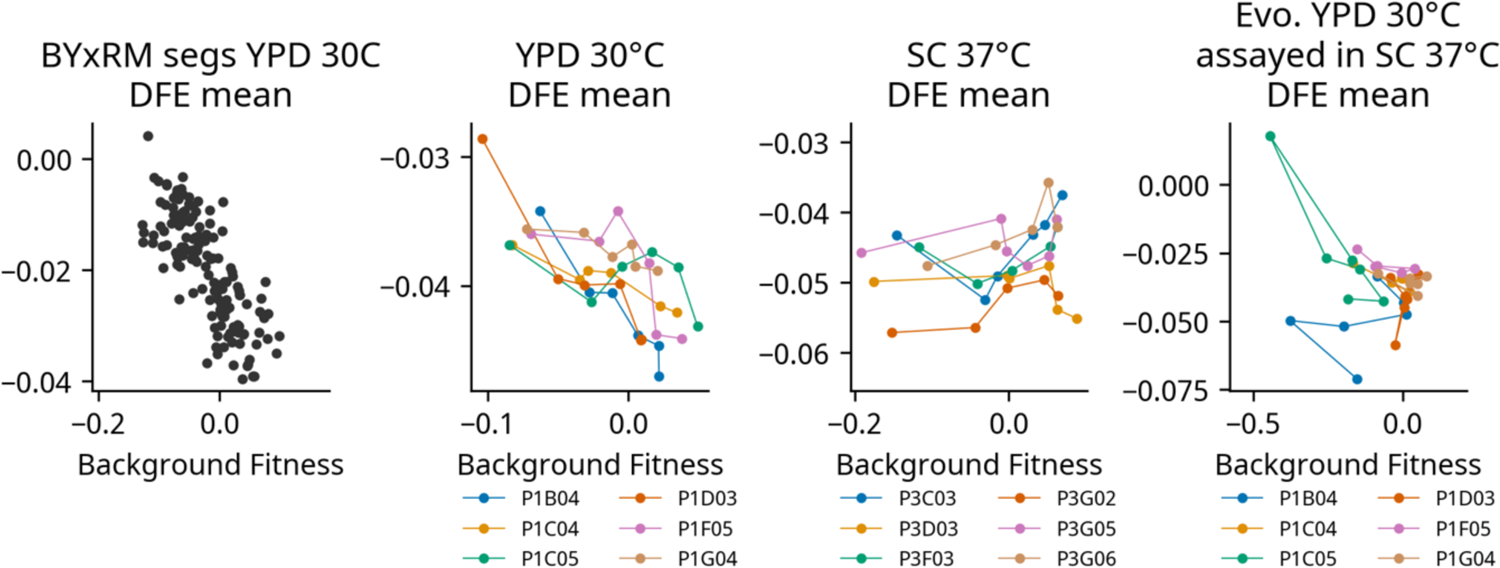
Comparison of our DFE mean vs. background fitness data with the data from Johnson et al. (2019). Each graph shows the relationship between the mean of the DFE and strain background. All graphs have the same aspect ratio.

**Figure 1 – figure supplement 7.**
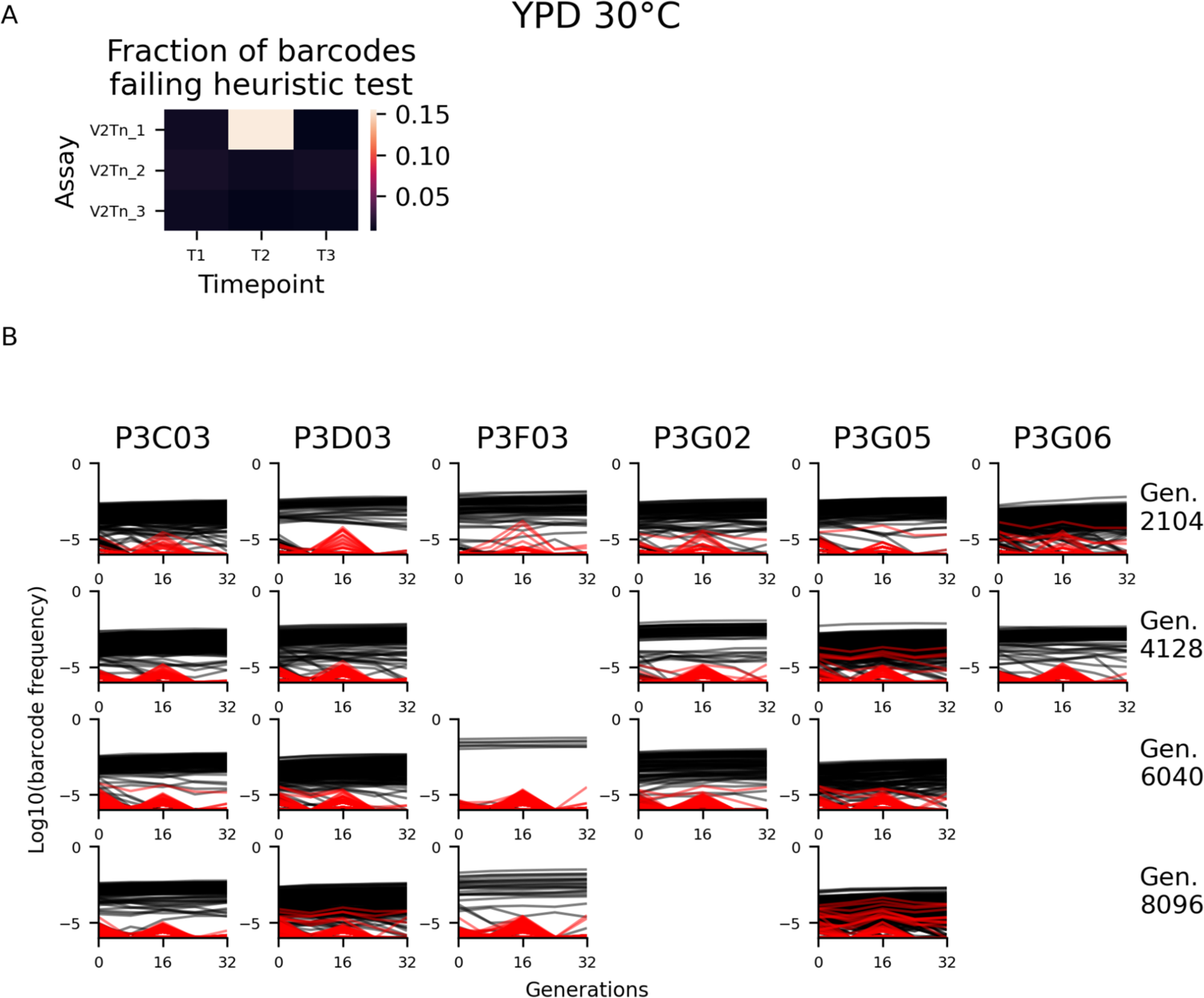
Excluding barcodes that experience sequencing cross contamination. **(A)** Heatmap showing the percentage of barcodes that failed our heuristic test for sequencing cross contamination at each timepoint in each assay in our SC 37°C experiment. Based on this data, we excluded barcodes that failed this test at timepoint 2 in the V2Tn_2 assay. Frequency trajectories of reference mutation barcodes are plotted in **(B)**, with excluded barcodes shown in red. This procedure excluded 0.4% of reads at timepoint 2 on average. See methods for more details.

**Figure 2 – figure supplement 1.**
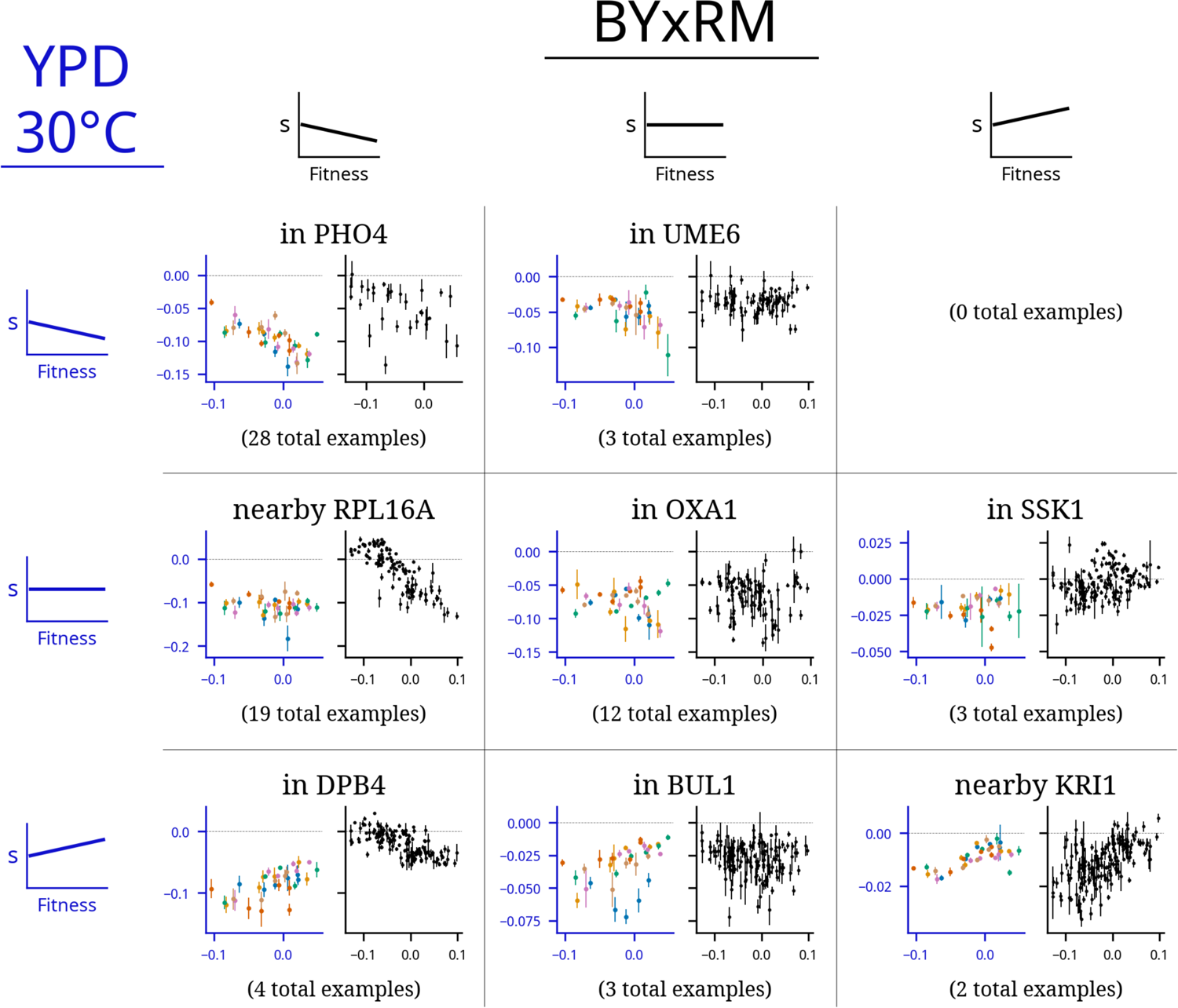
Comparison of patterns of fitness-correlated epistasis between YPD 30°C and a previous study. Each panel shows an example of a specific mutation with a particular combination of relationships (negative, positive, or non-significant correlation between fitness effect of the mutation, *s,* and background fitness) in the two environments; numbers indicate the total number of mutations displaying each pair of relationships. Graphs depict fitness effect (y-axis) as a function of background fitness (x-axis). The axes are colored to identify the environment: in each square the blue axes on the left is data from YPD 30°C and the black axes on the right is data from Johnson et al. (2019), in which each point represents a segregant from a yeast cross. Points are colored by populations, as in Figure 1. Each set of example plots is labeled by where the mutation is in the genome (what gene it disrupts).

**Figure 2 – figure supplement 2.**
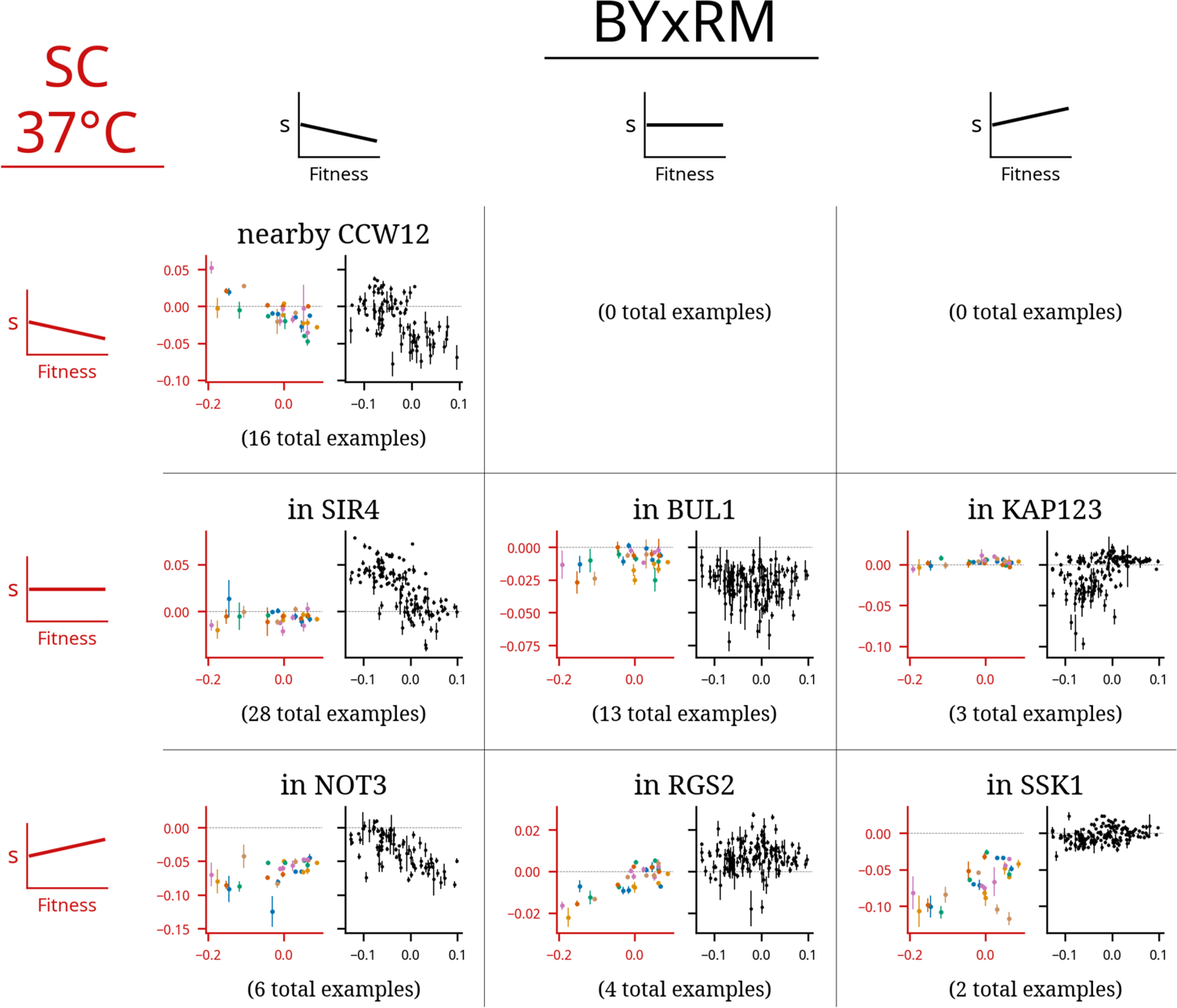
Comparison of patterns of fitness-correlated epistasis between SC 37°C and a previous study. Each panel shows an example of a specific mutation with a particular combination of relationships (negative, positive, or non-significant correlation between fitness effect of the mutation, *s,* and background fitness) in the two environments; numbers indicate the total number of mutations displaying each pair of relationships. Graphs depict fitness effect (y-axis) as a function of background fitness (x-axis). The axes are colored to identify the environment: in each square the red axes on the left is data from SC 37°C and the black axes on the right is data from Johnson et al. (2019), in which each point represents a segregant from a yeast cross. Points are colored by populations, as in Figure 1. Each set of example plots is labeled by where the mutation is in the genome (what gene it disrupts).

**Figure 2 – figure supplement 3.**
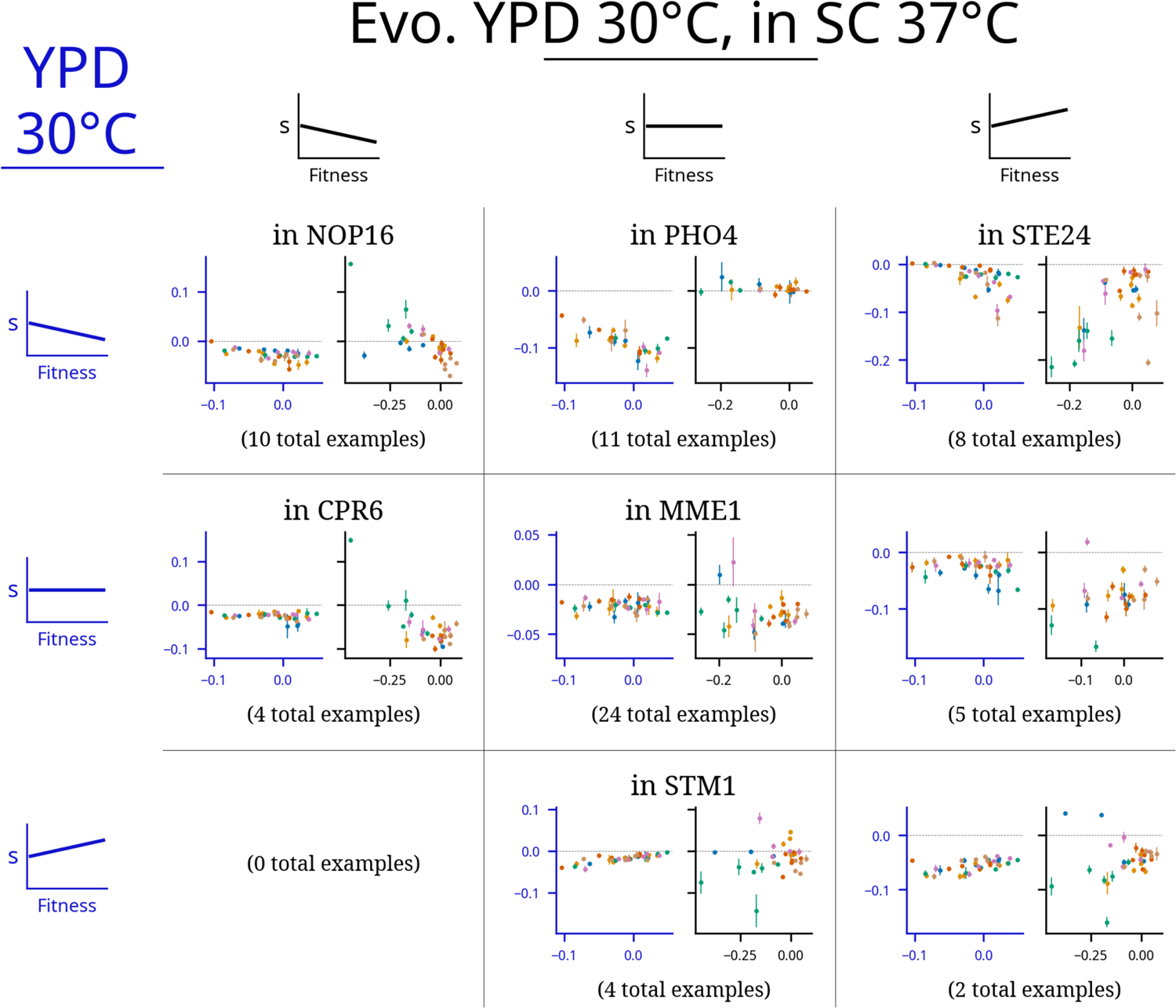
Comparison of patterns of fitness-correlated epistasis YPD 30°C and SC 37°C, in both cases using the set of clones isolated from evolution in YPD 30°C. Each panel shows an example of a specific mutation with a particular combination of relationships (negative, positive, or non-significant correlation between fitness effect of the mutation, *s,* and background fitness) in the two environments; numbers indicate the total number of mutations displaying each pair of relationships. Graphs depict fitness effect (y-axis) as a function of background fitness (x-axis). The axes are colored to identify the environment: in each square the blue axes on the left is data from YPD 30°C and the black axes on the right is data from the same clones isolated from evolution in YPD°C, but assayed in the SC 37°C environment. Points are colored by populations, as in Figure 1. Each set of example plots is labeled by where the mutation is in the genome (what gene it disrupts).

**Figure 2 – figure supplement 4.**
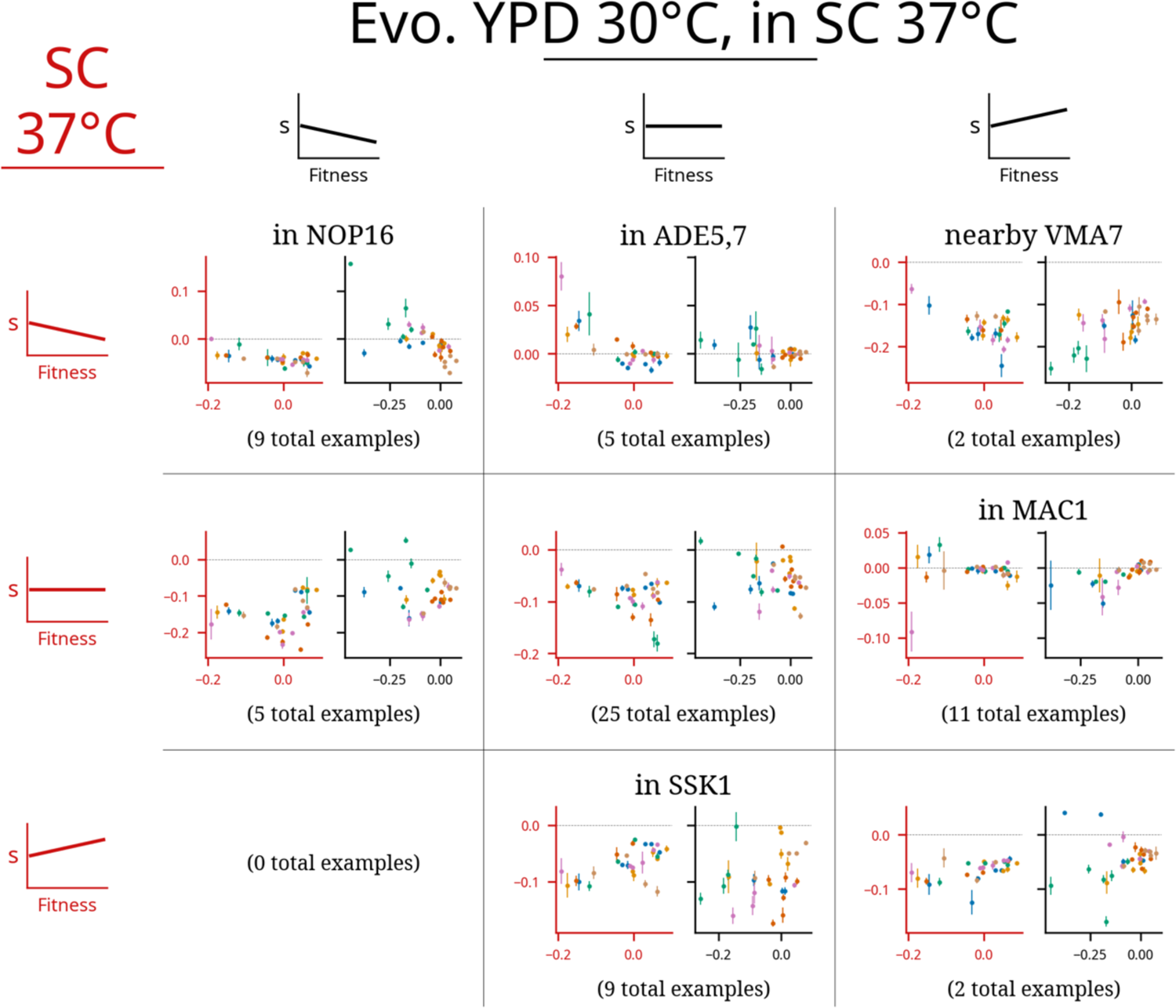
Comparison of patterns of fitness-correlated epistasis in the SC 37°C environment for clones isolated from evolution in either YPD 30°C or SC 37°C. Each panel shows an example of a specific mutation with a particular combination of relationships (negative, positive, or non-significant correlation between fitness effect of the mutation, *s,* and background fitness) in the two environments; numbers indicate the total number of mutations displaying each pair of relationships. Graphs depict fitness effect (y-axis) as a function of background fitness (x-axis). The axes are colored to identify the environment: in each square the red axes on the left is data from clones isolated from evolution in SC 37°C and the black axes on the right is data from clones isolated from evolution in YPD°C, both assayed in the SC 37°C environment. Points are colored by populations, as in Figure 1. Each set of example plots is labeled by where the mutation is in the genome (what gene it disrupts).

**Figure 3 – figure supplement 1.**
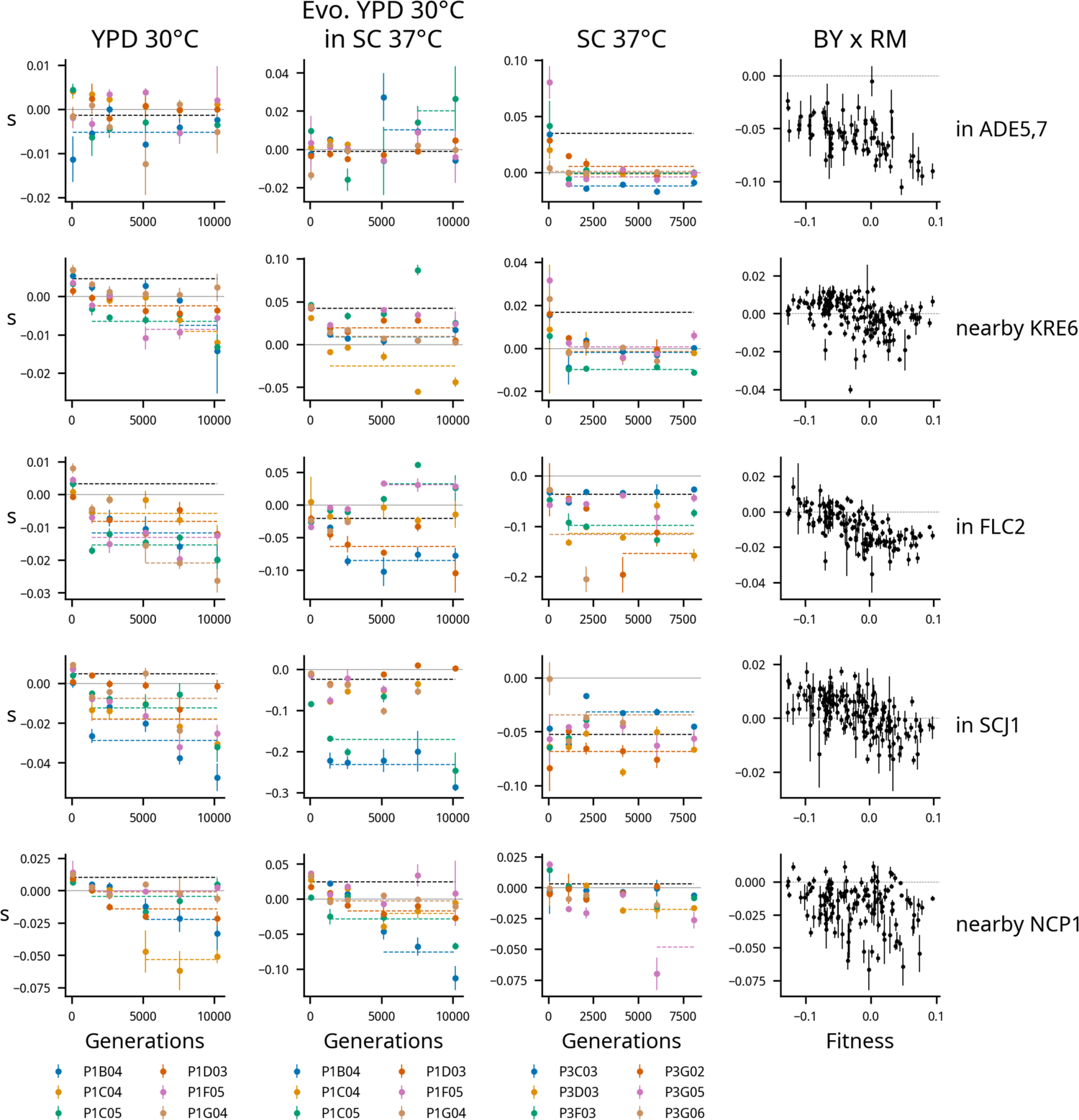
Epistasis in mutations that are beneficial on average at the first timepoint in at least one environment. Each row represents an insertion mutation, labeled with the gene it disrupts or a gene nearby its insertion site if it is intergenic. Each column represents a condition from our experiment or data from Johnson et al. (2019) (right). Each plot shows the fitness effects of mutations over time in each population, with idiosyncratic model fits shown, as in Figure 3. The plots in the right column show the fitness effect of each mutation as a function of background fitness. Each of the mutations displayed here has a beneficial fitness effect on average in at least one environment that becomes neutral or deleterious in that environment during evolution.

**Figure 3 – figure supplement 2.**
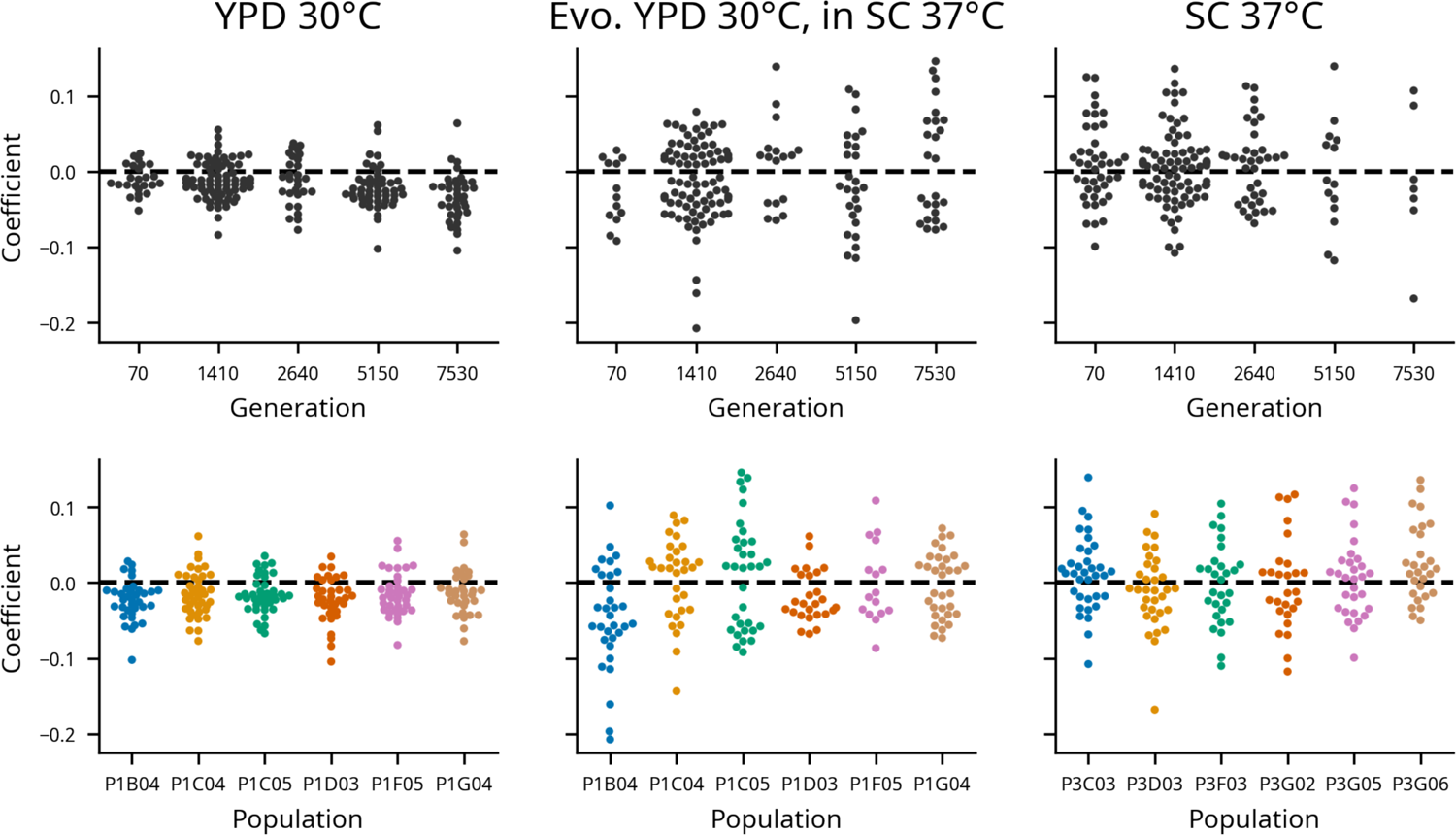
Idiosyncratic model coefficients, broken down by population and timepoint in each condition.

**Figure 3 – figure supplement 3.**
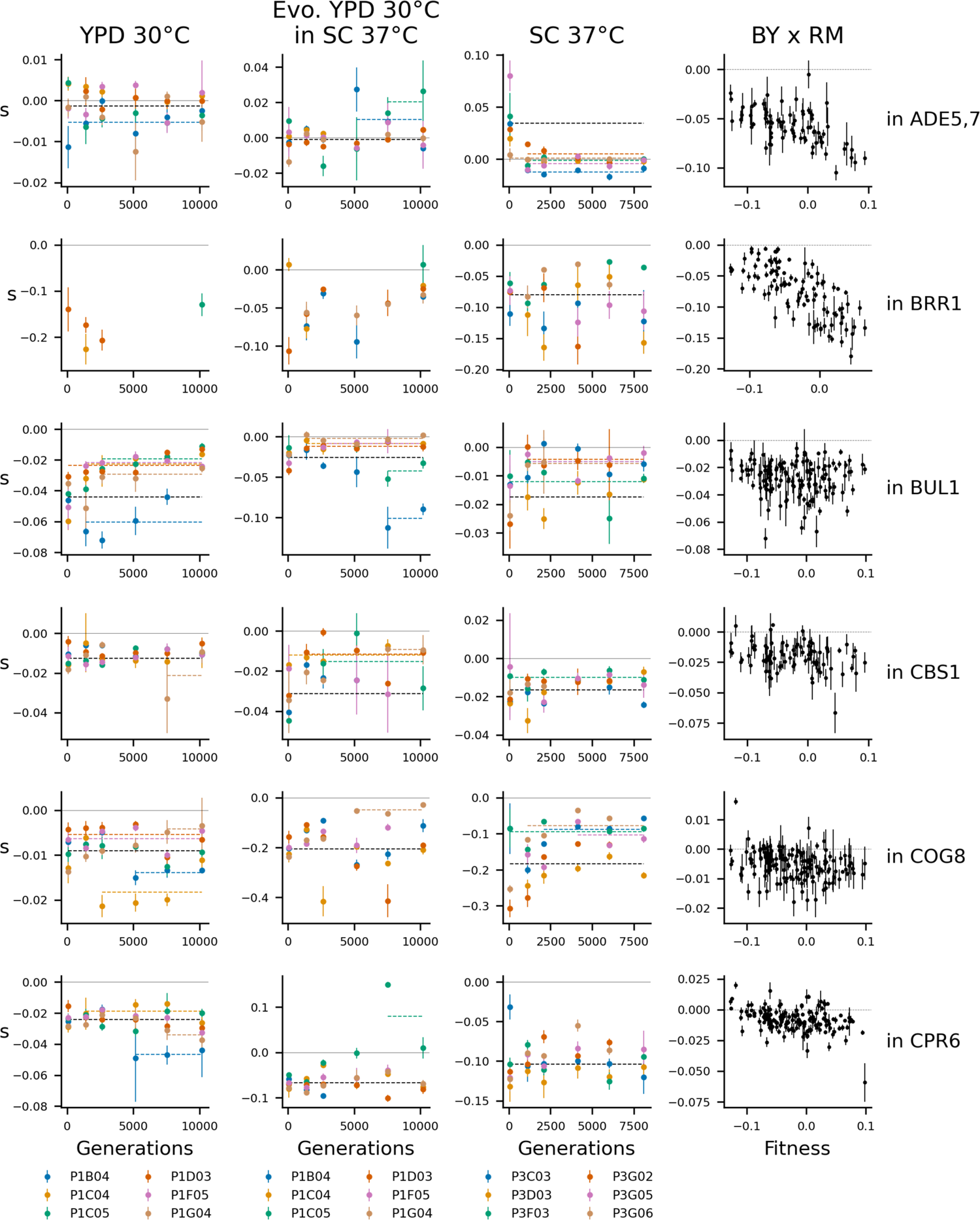
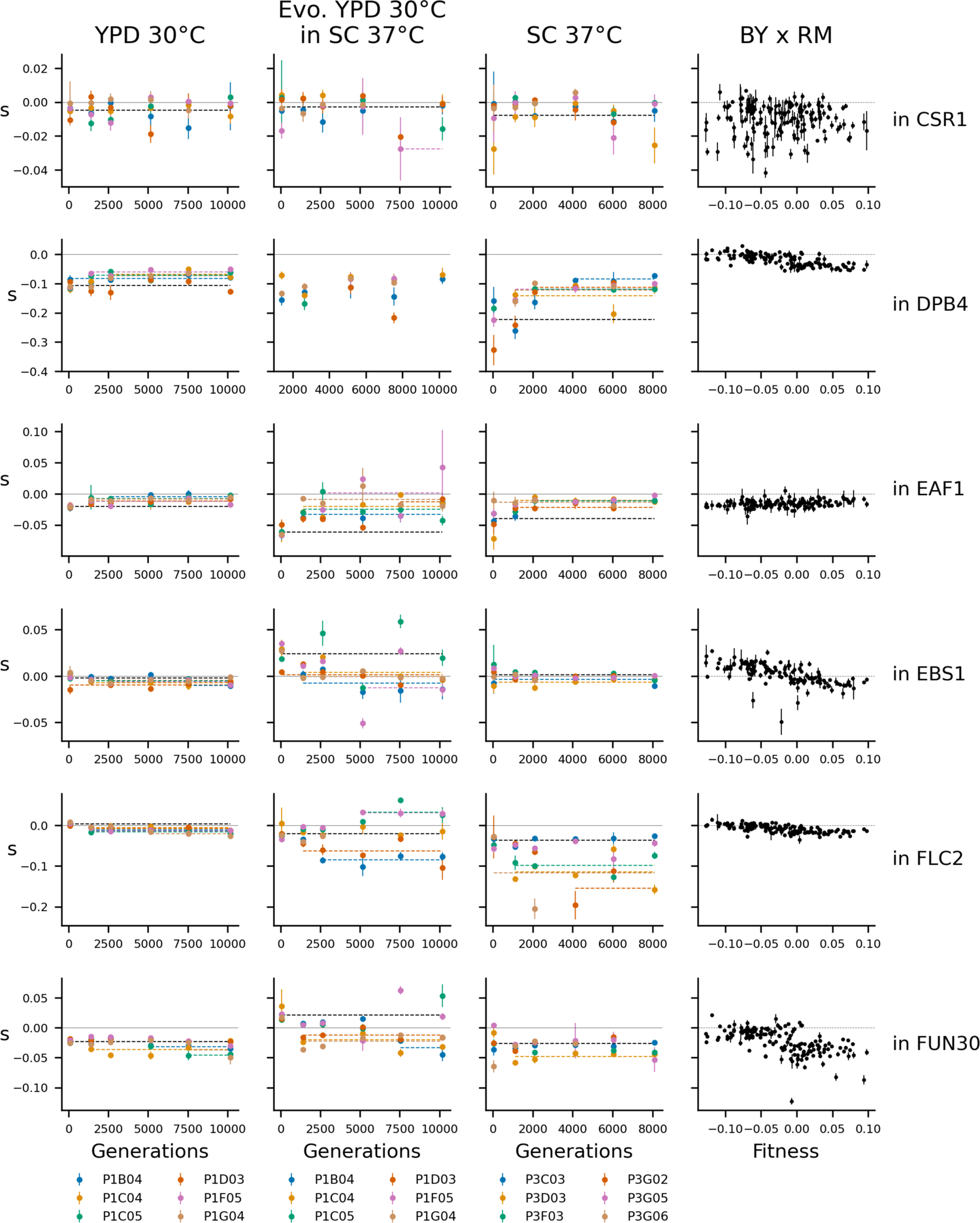
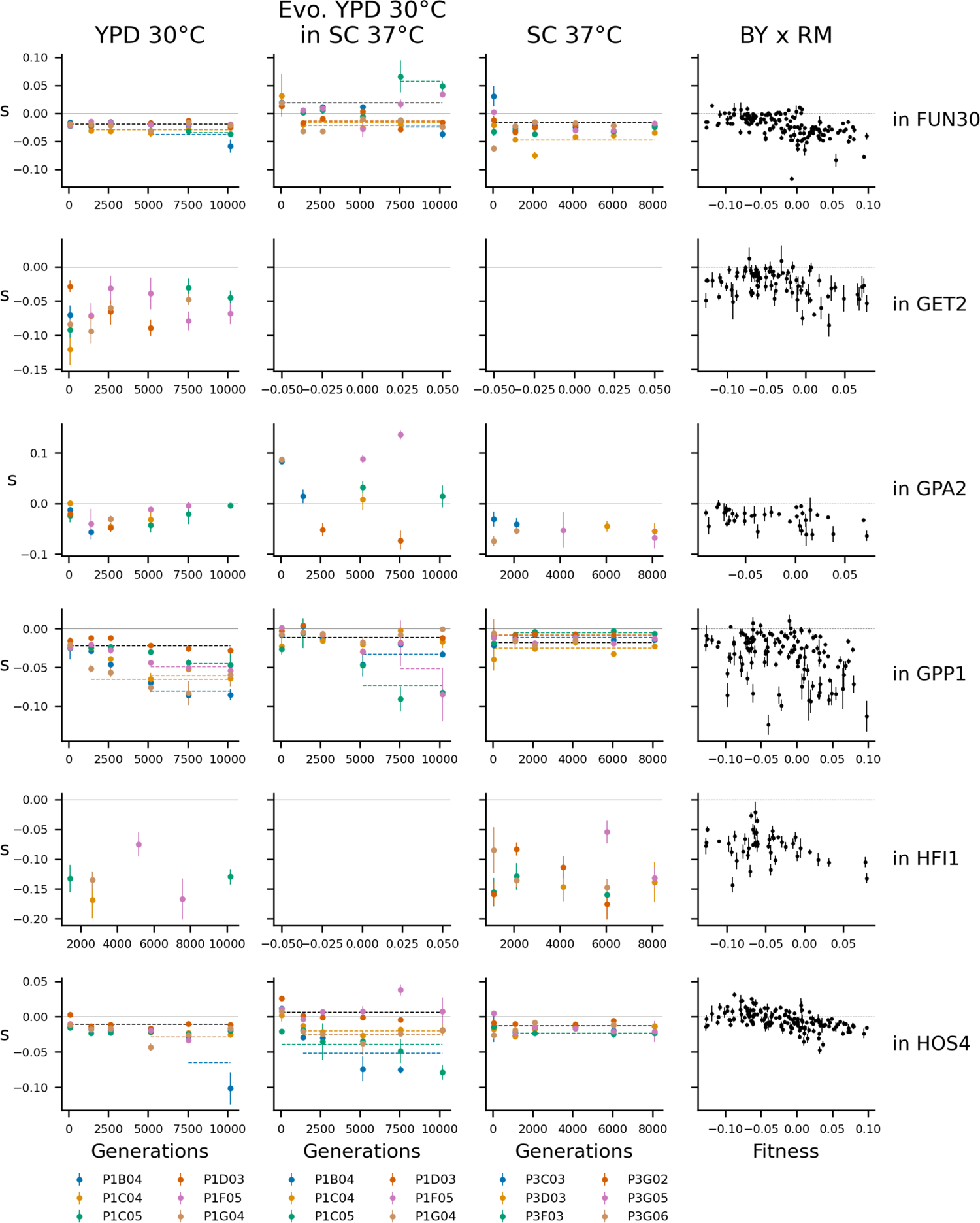
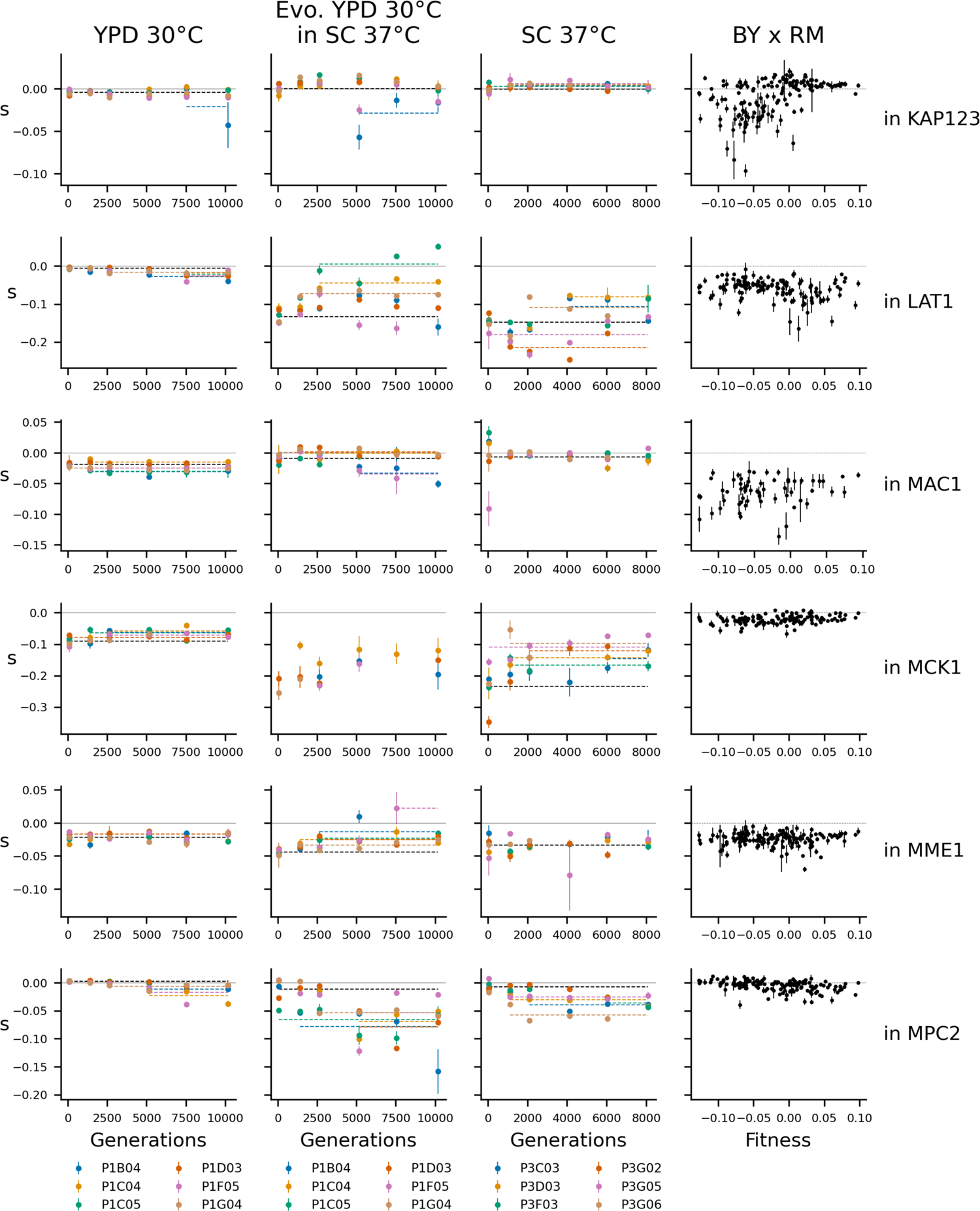
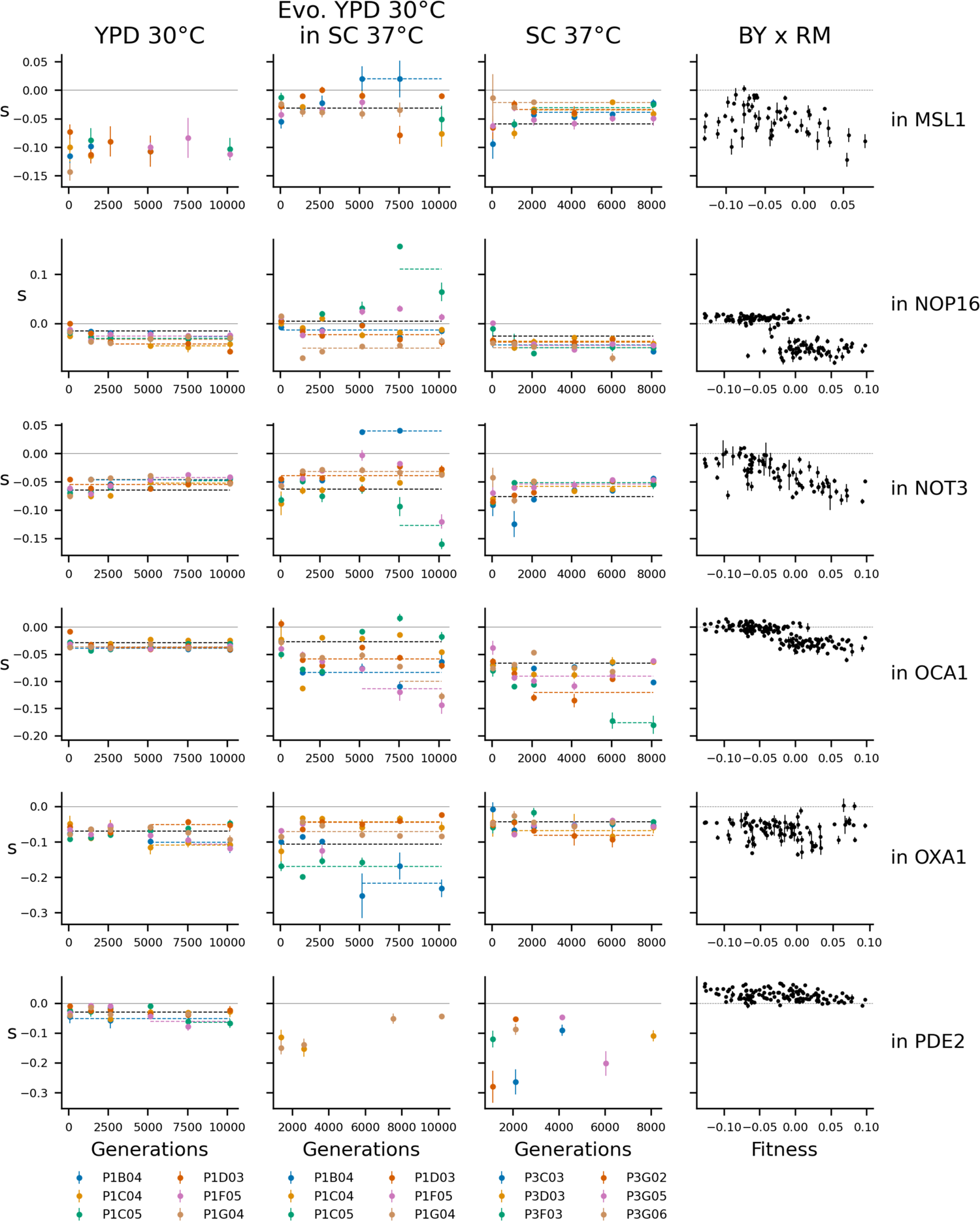
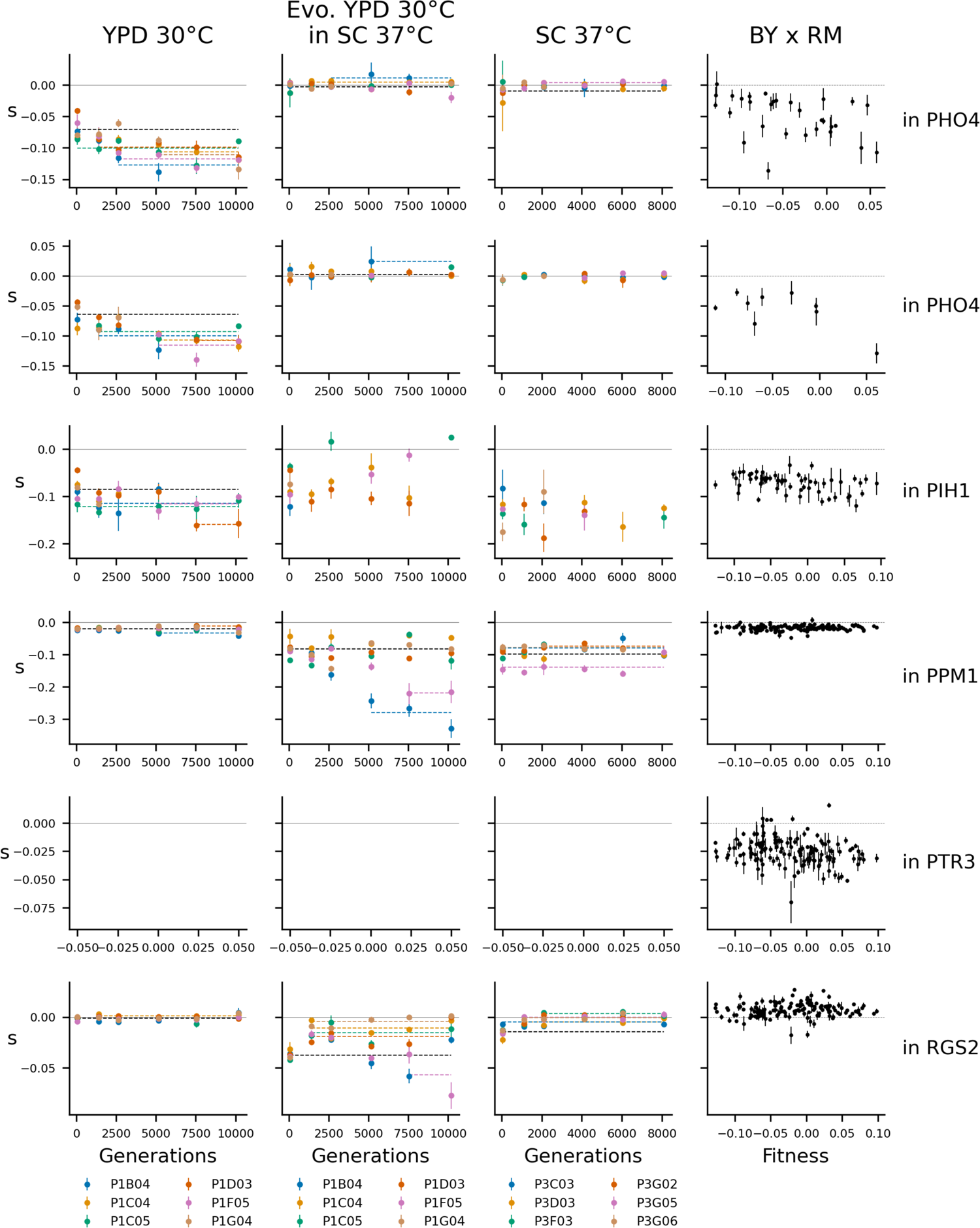
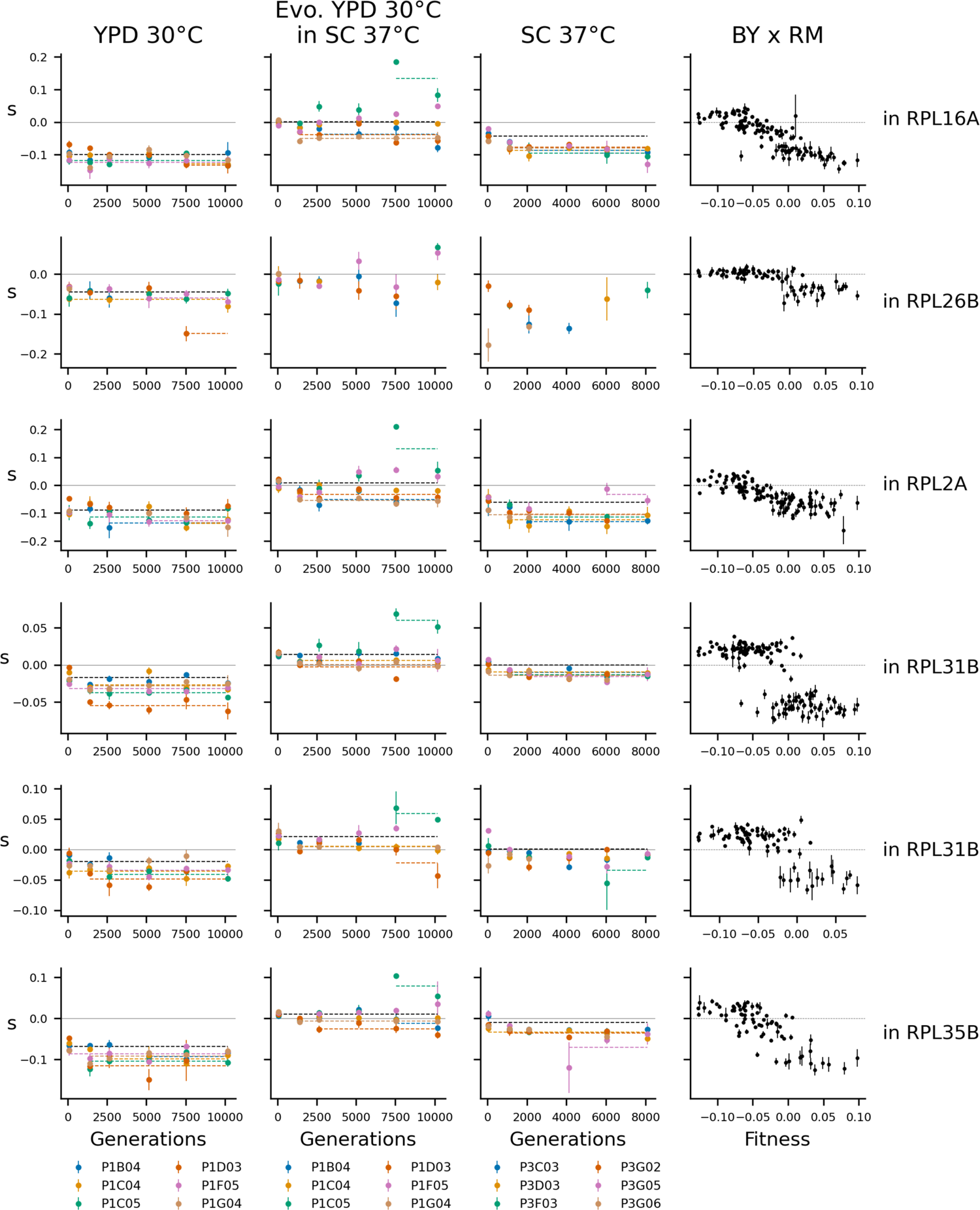
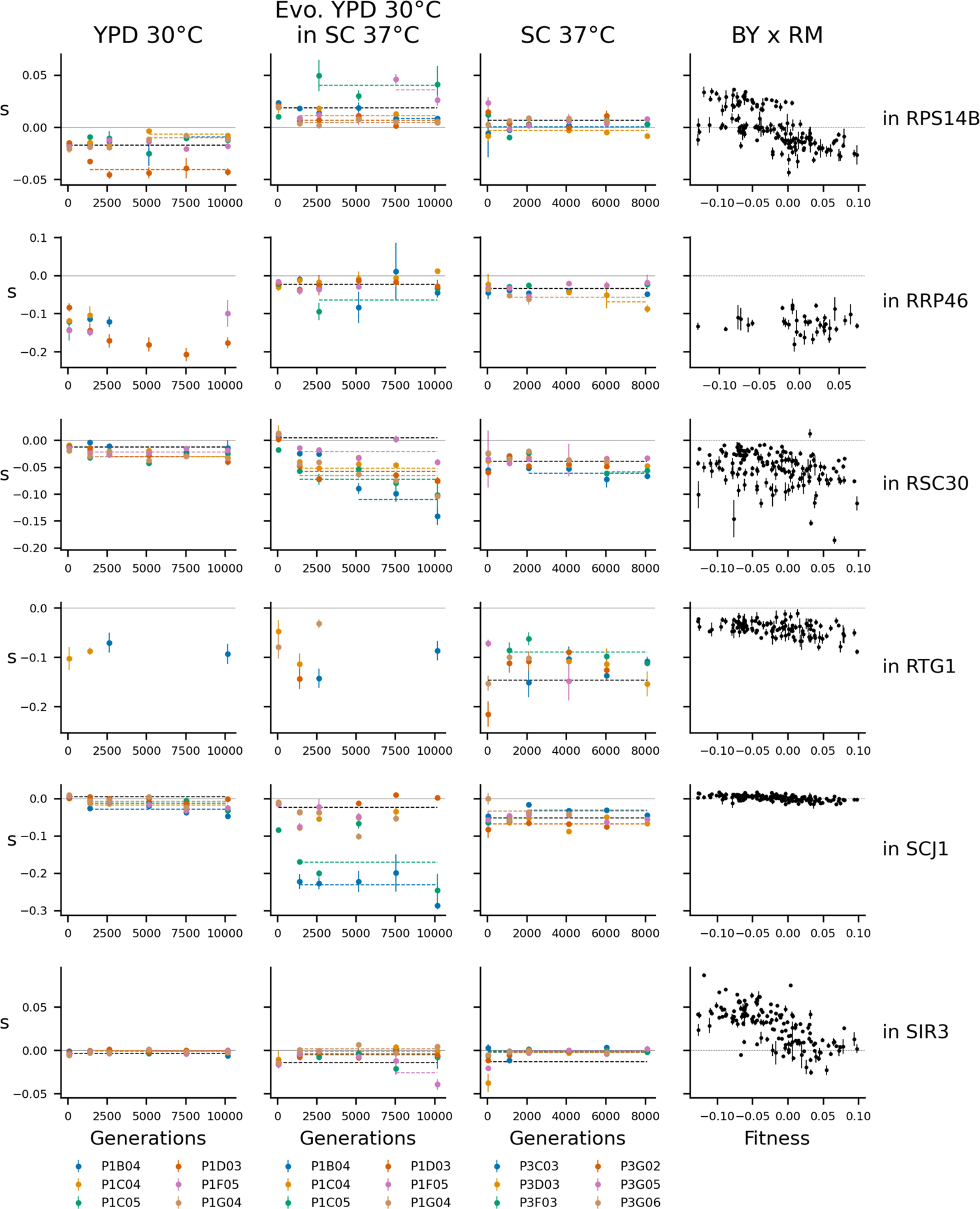
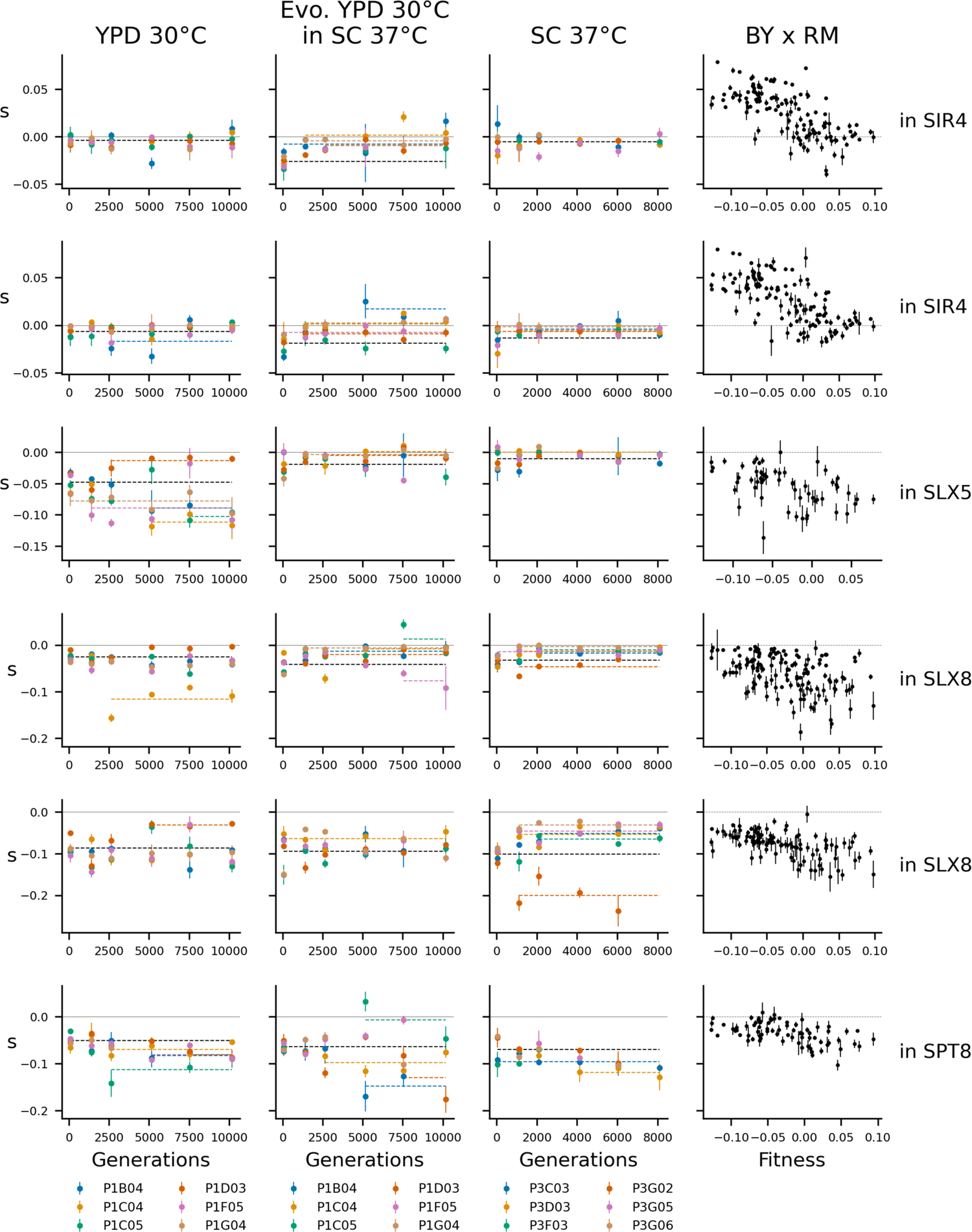
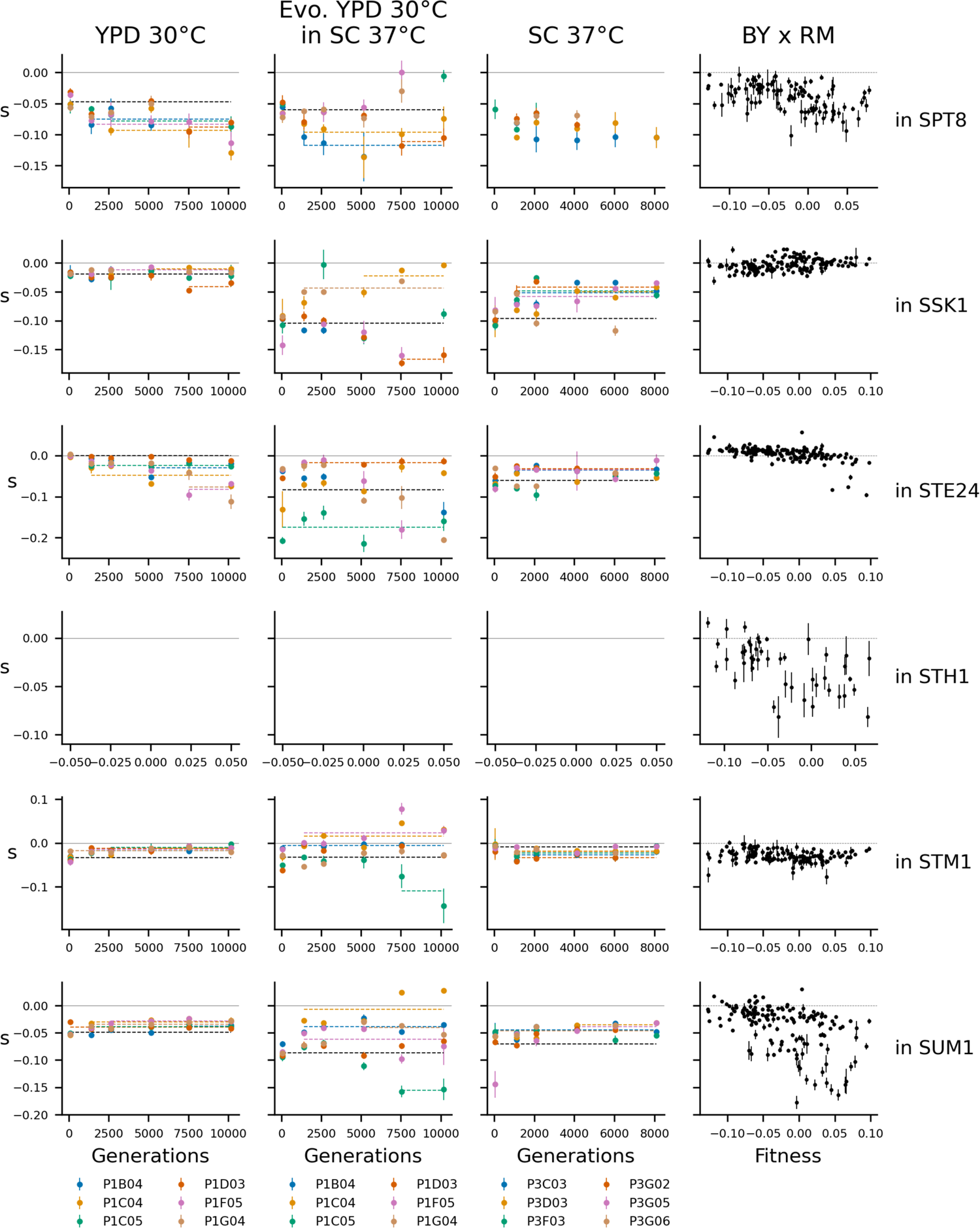
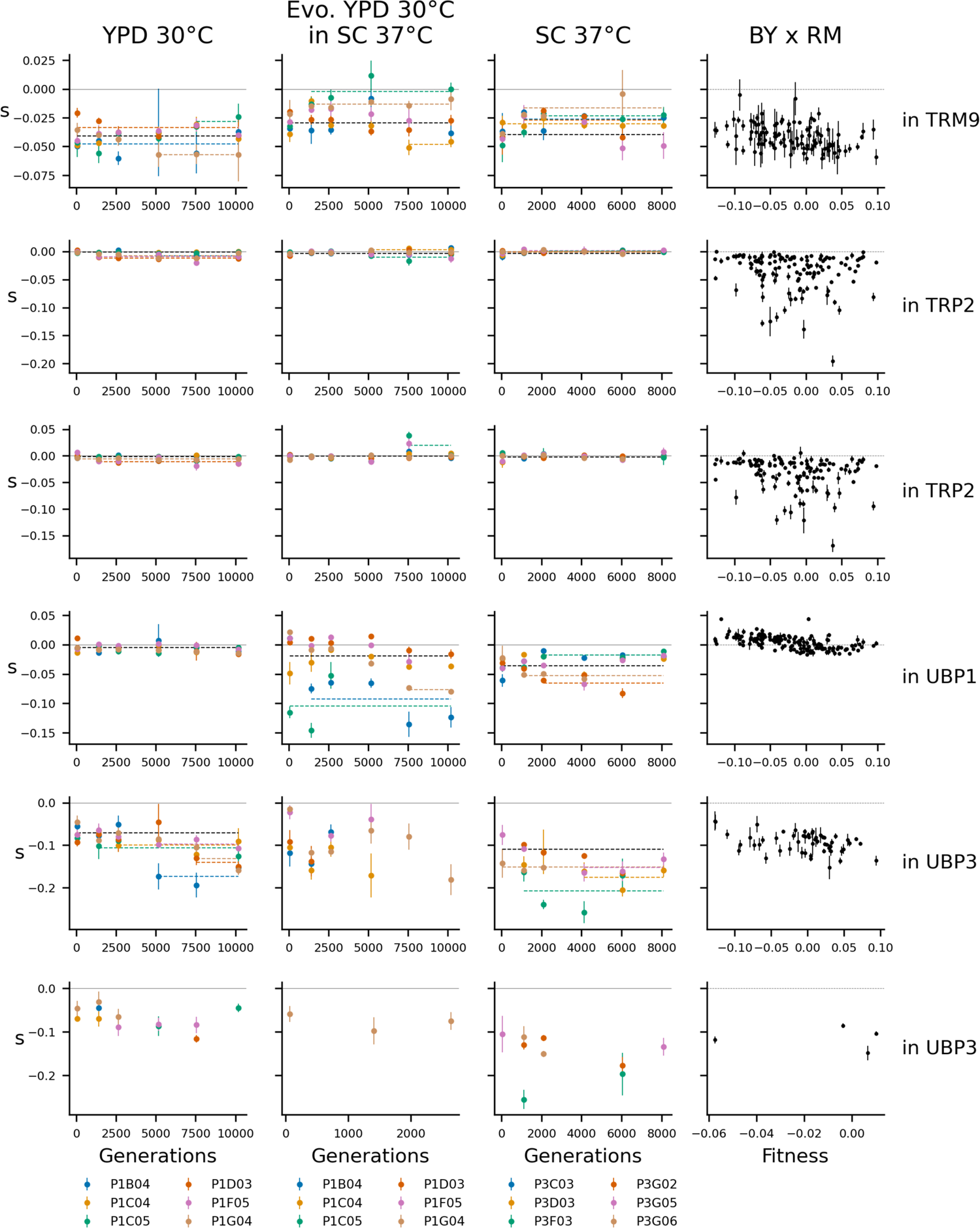
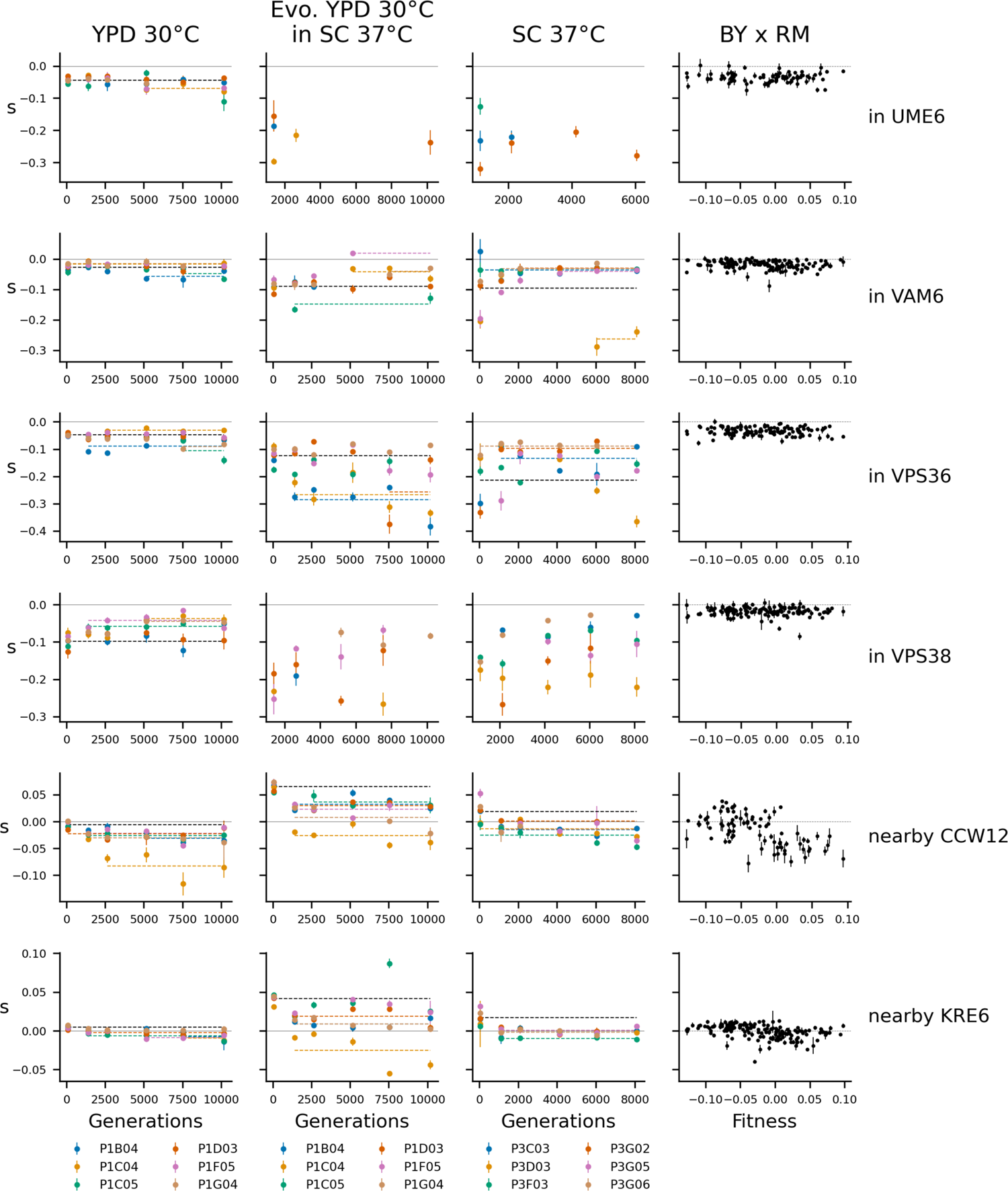
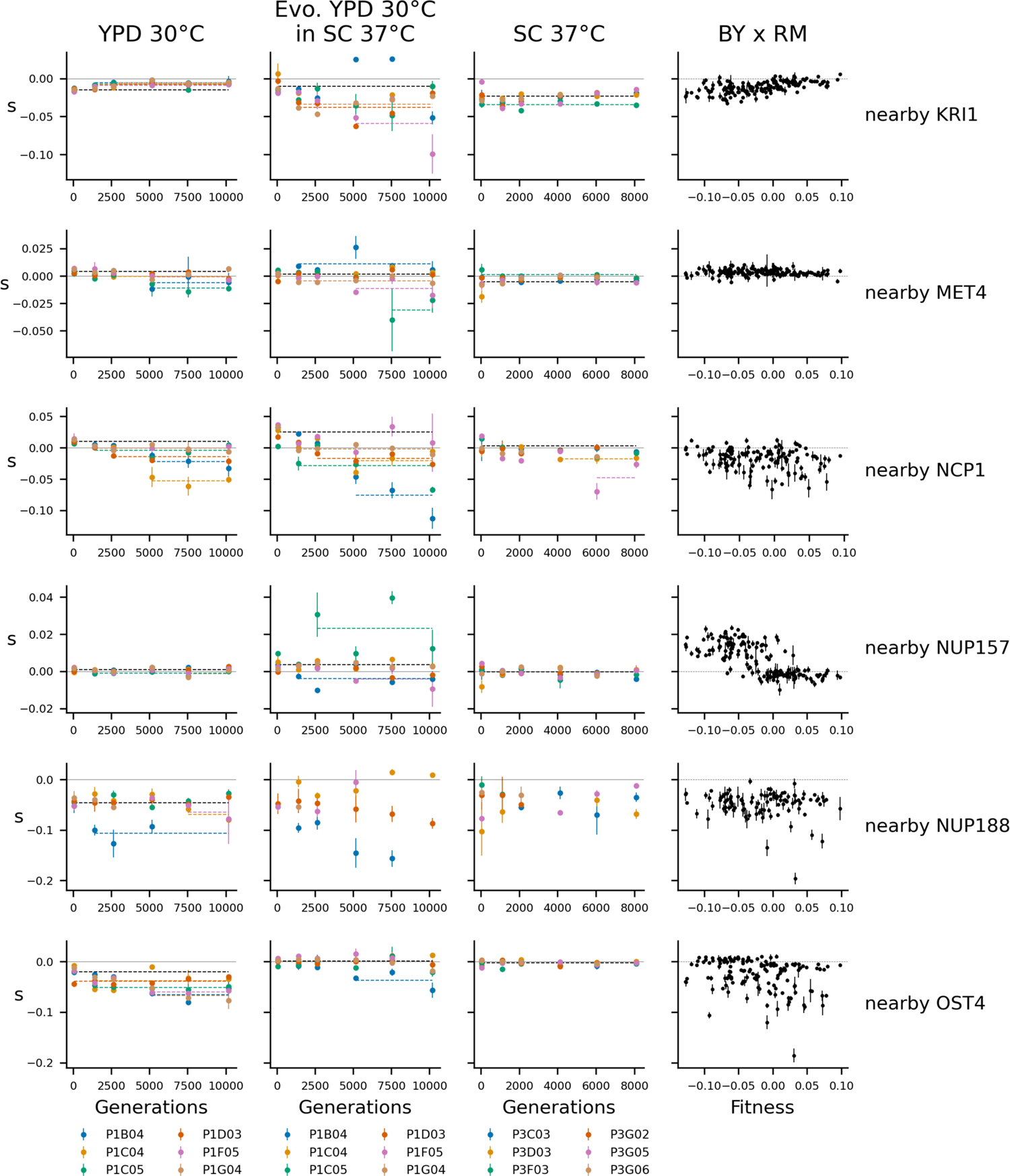
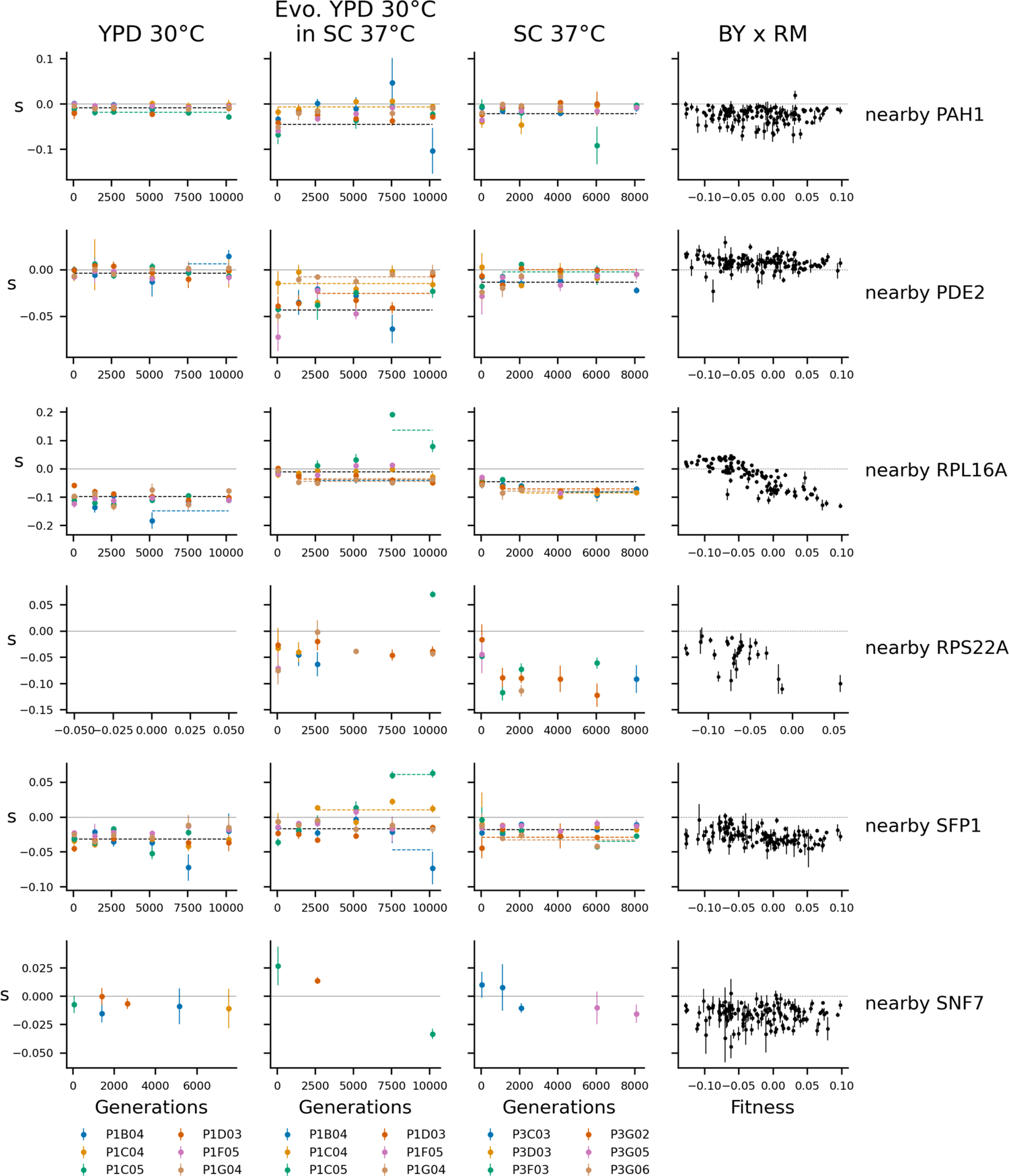
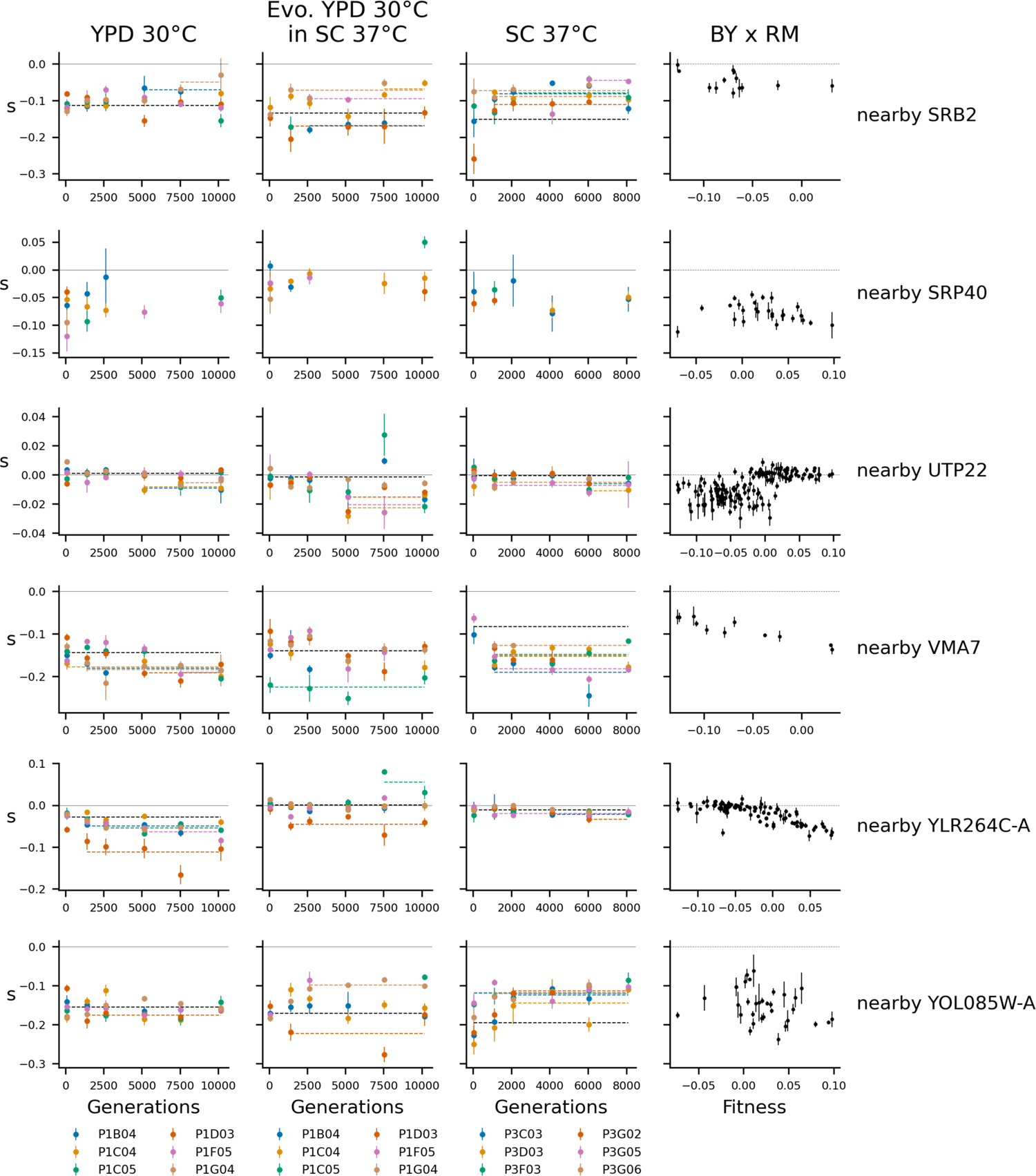
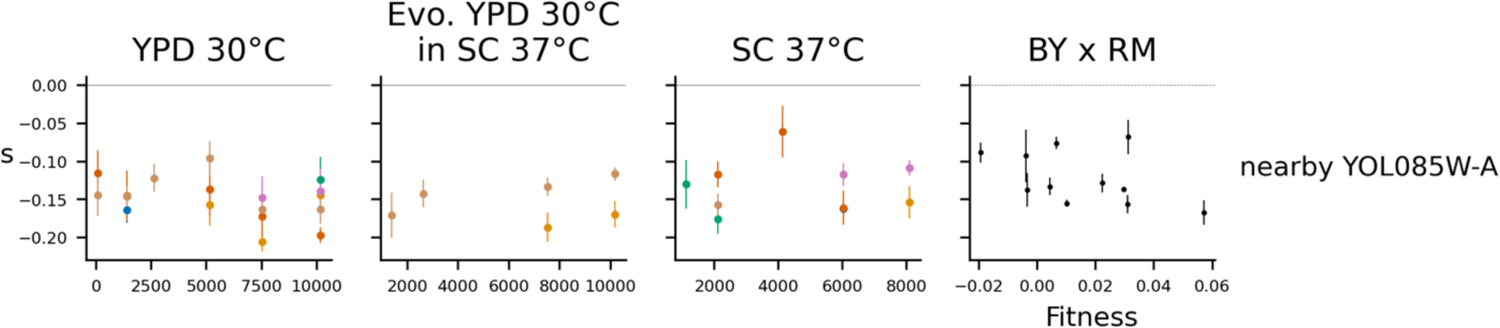
Determinants of fitness effects under the idiosyncratic model. Each row represents one mutation, labeled at right with by the gene it disrupts or the nearest start of a gene if it is intergenic. Each column represents a condition. The left three columns show the fitness effects of mutations over the course of evolution, with points colored by population. The right column shows the relationship between fitness and the fitness effect of the mutation in the segregants from a yeast cross (data from Johnson et al. (2019)). Model predictions are shown by dashed lines, and lines with contributions from indicator variables associated with a particular population are the same color as the points from that population.

**Figure 3 – figure supplement 4.**
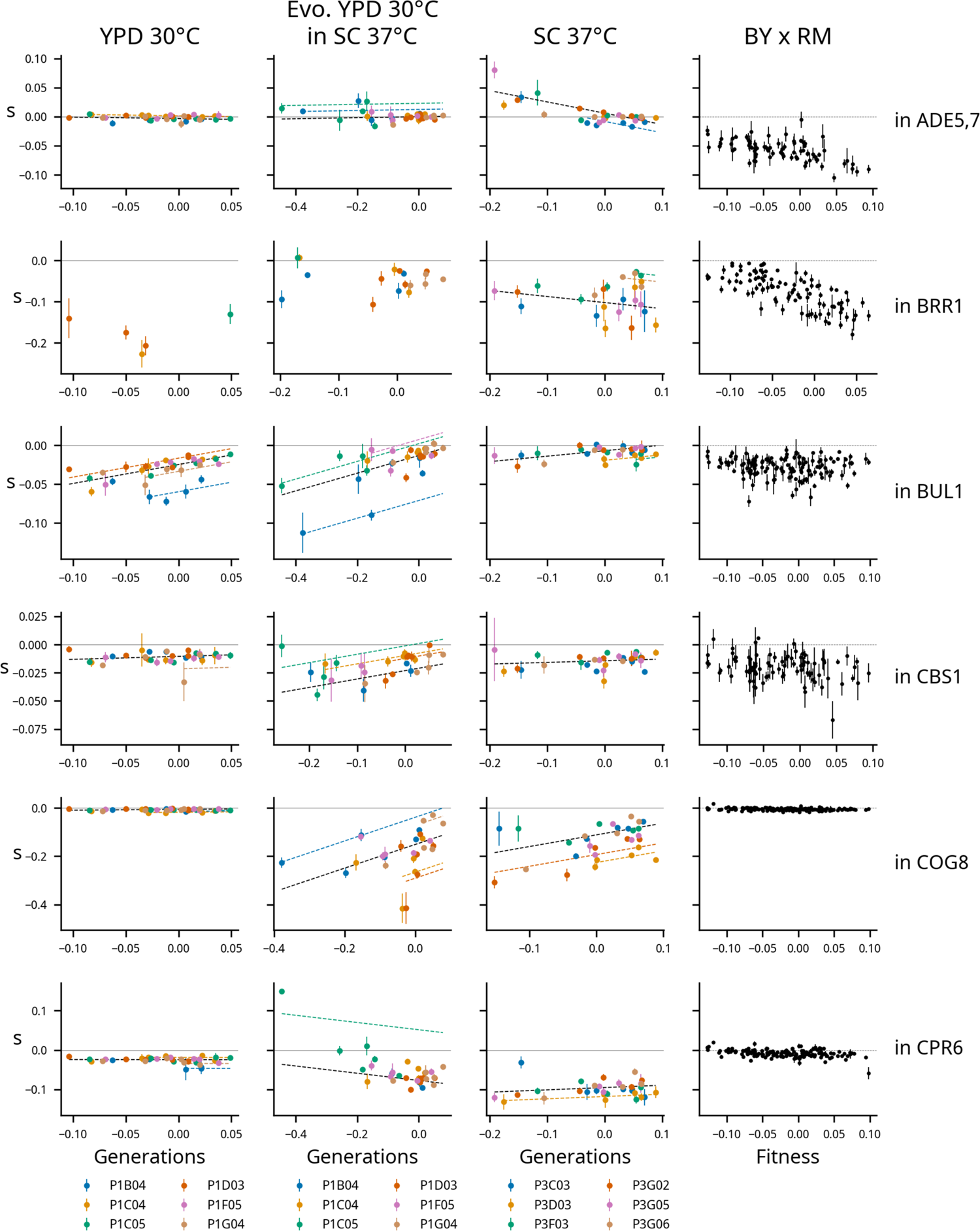
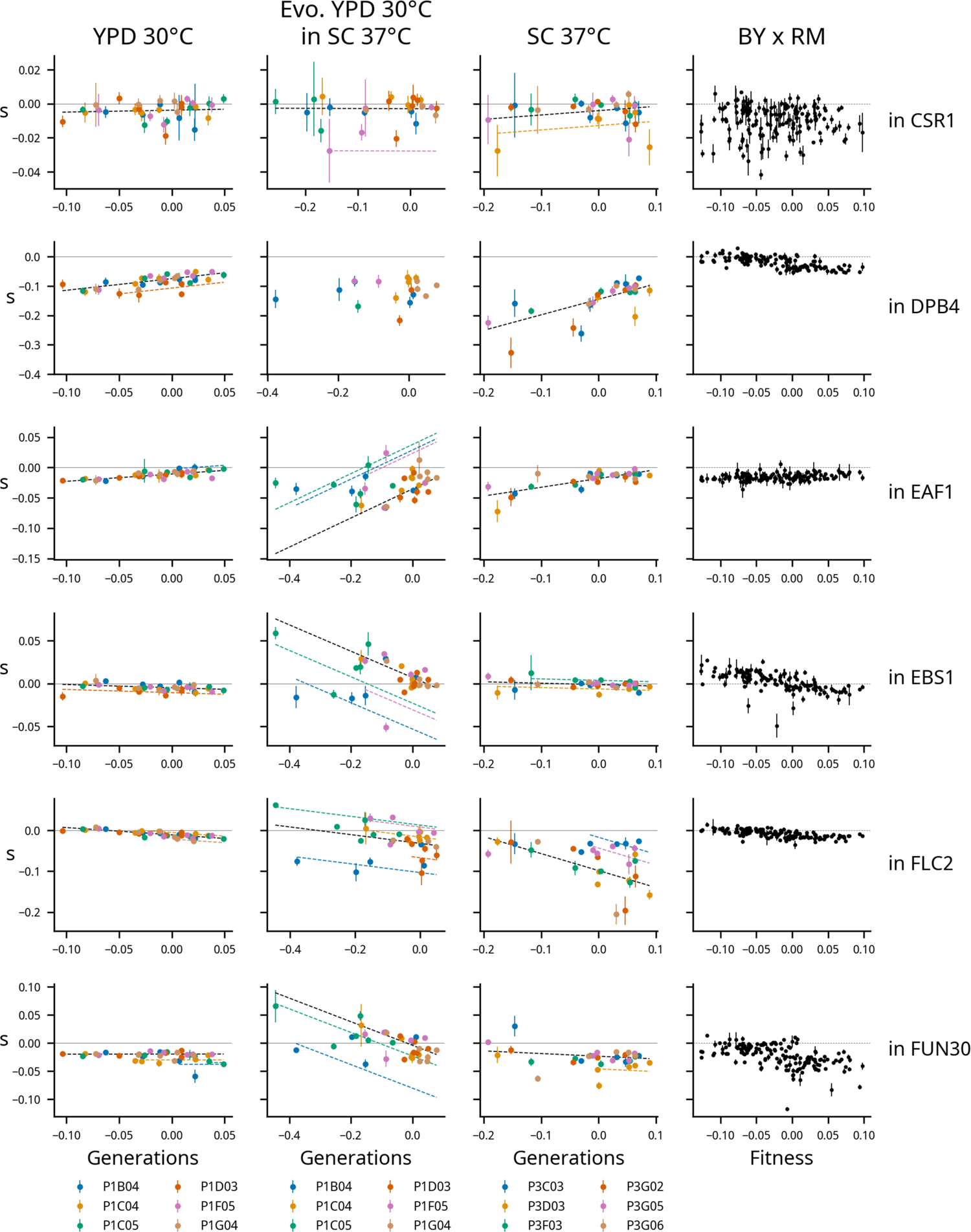
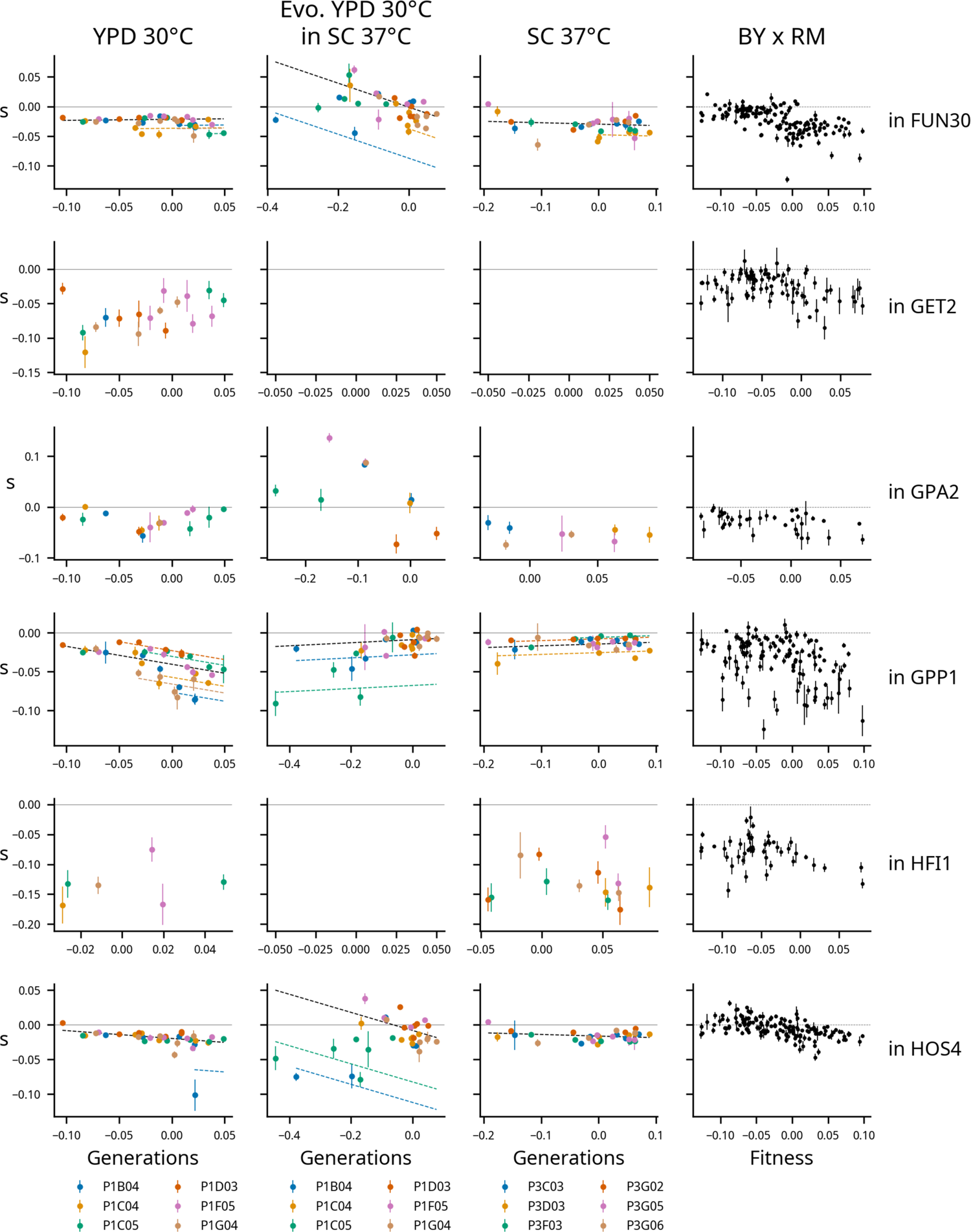
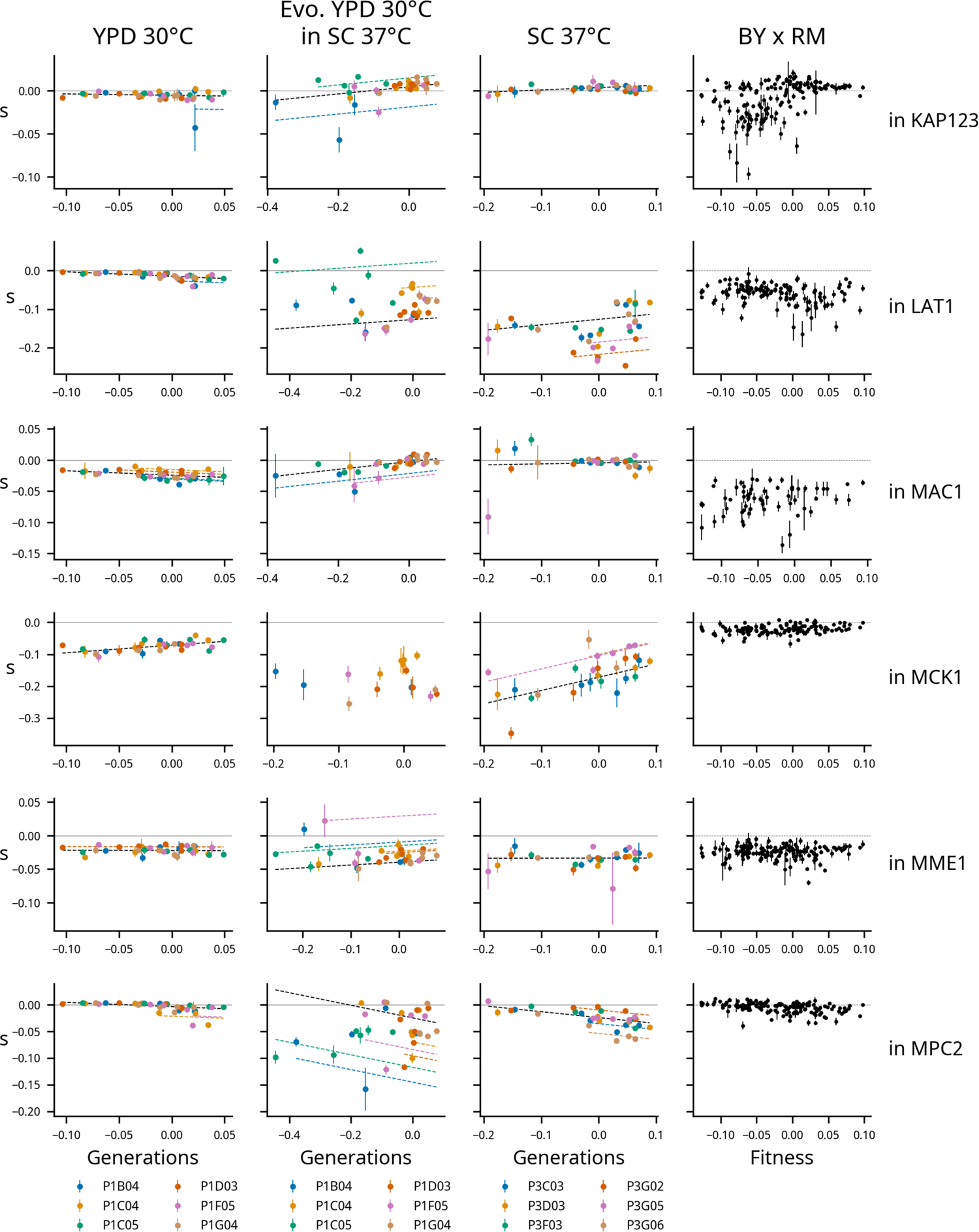
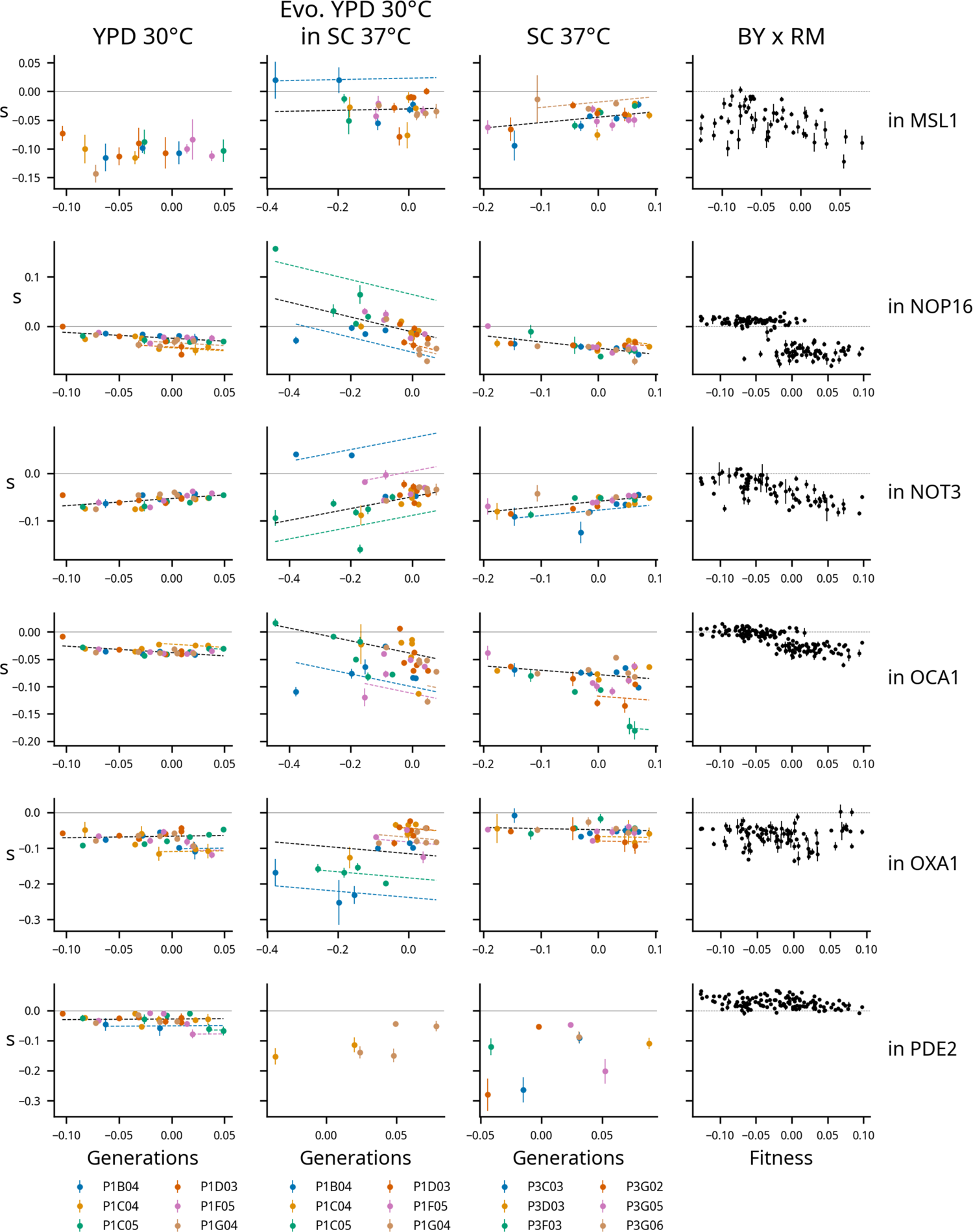
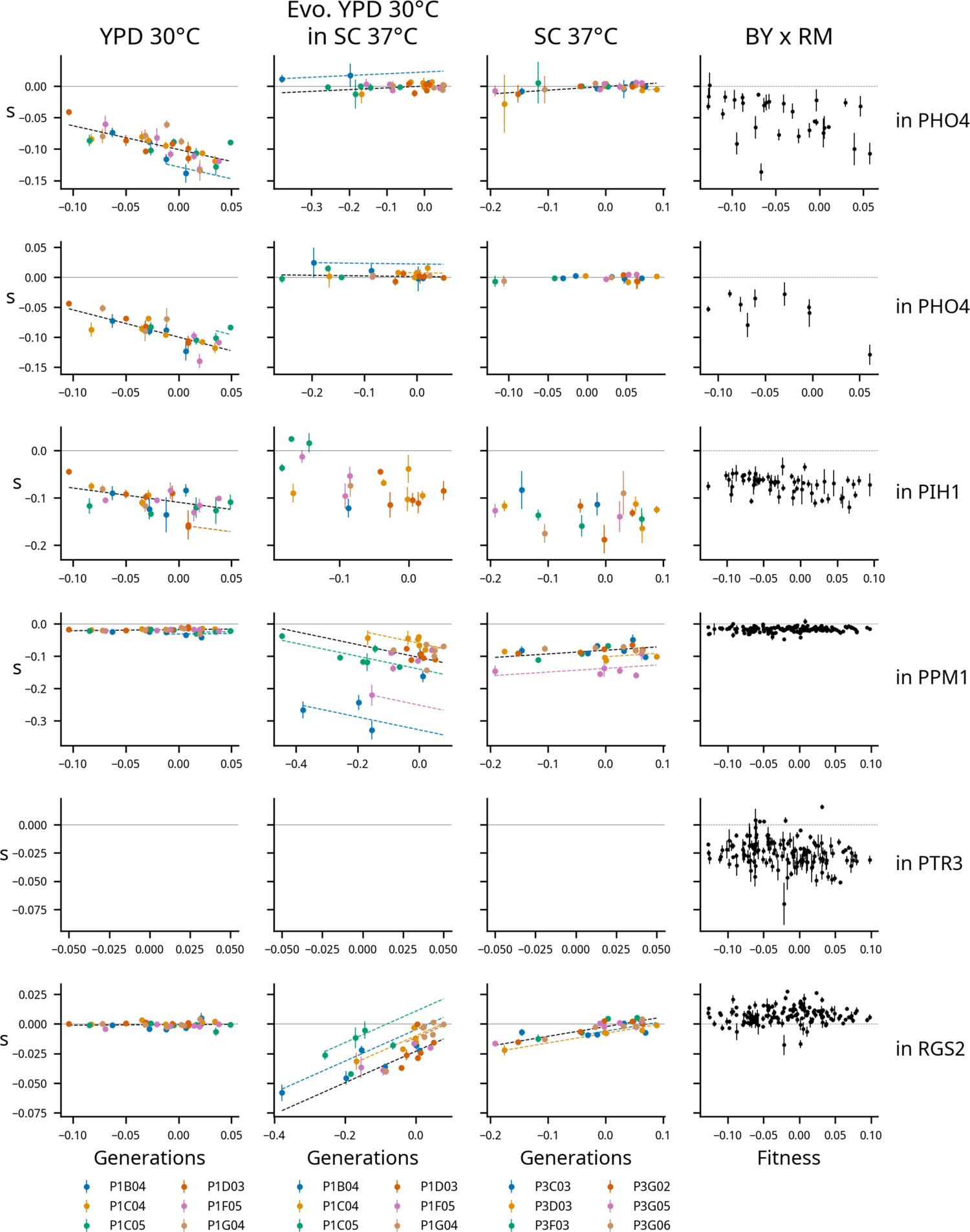
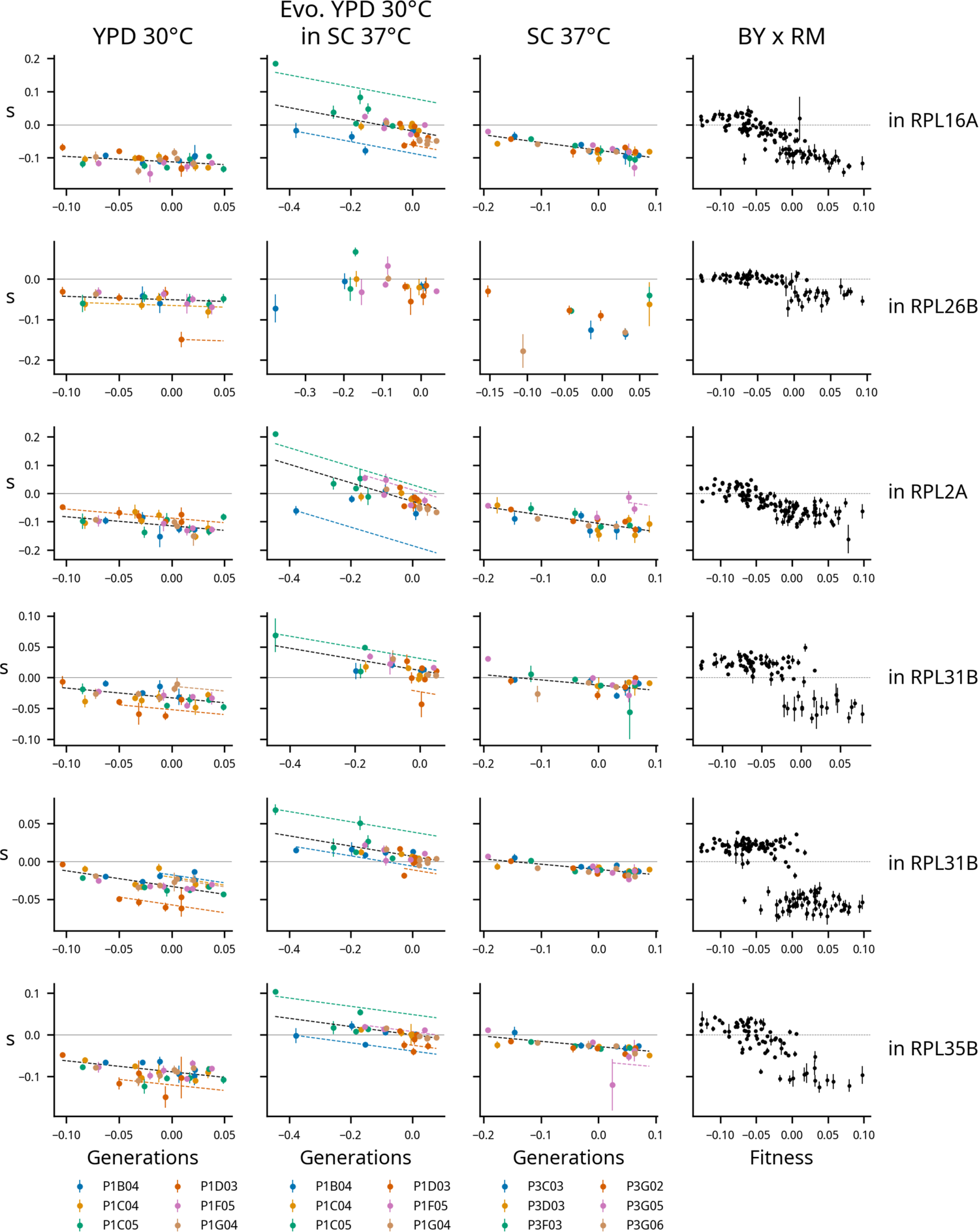
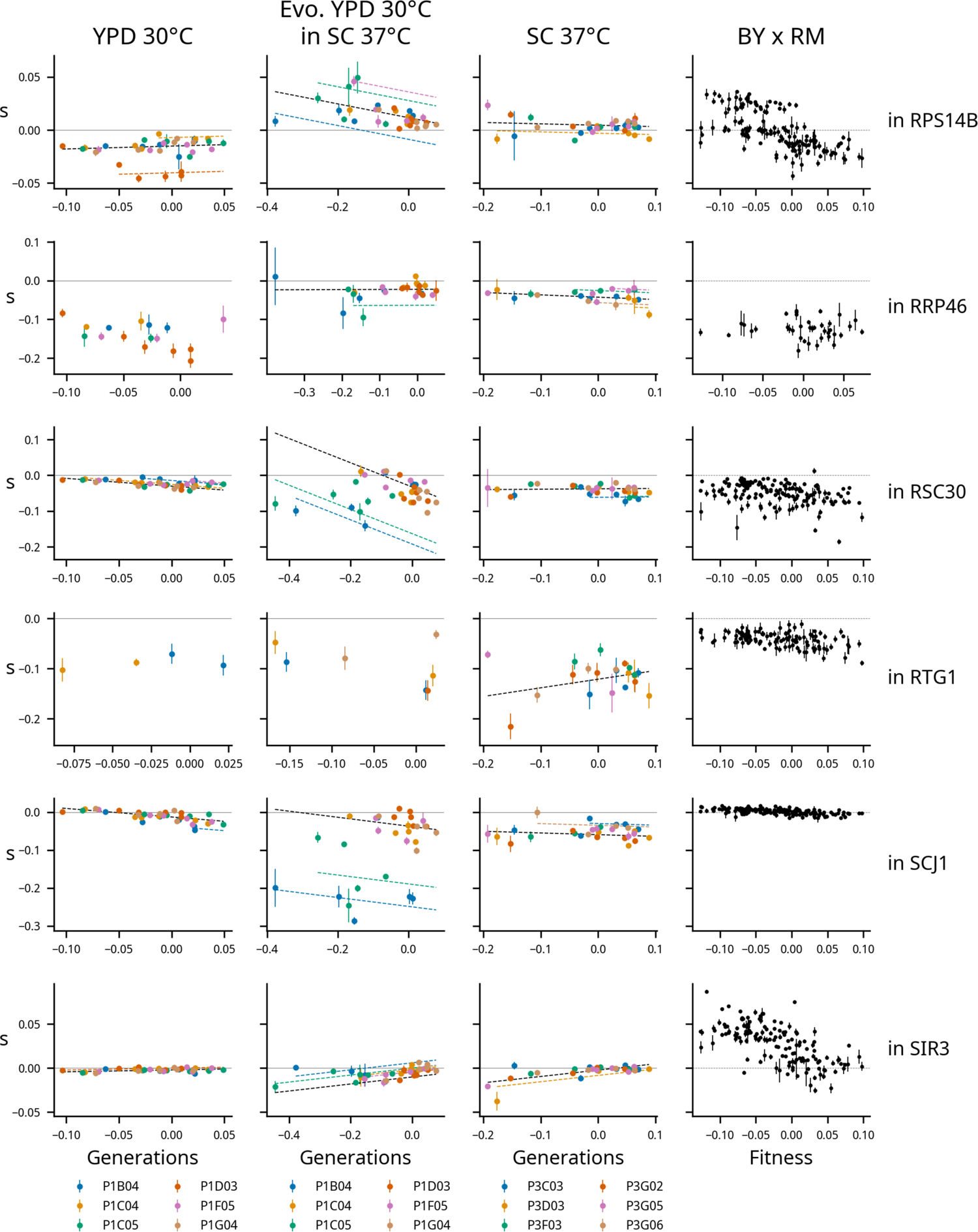
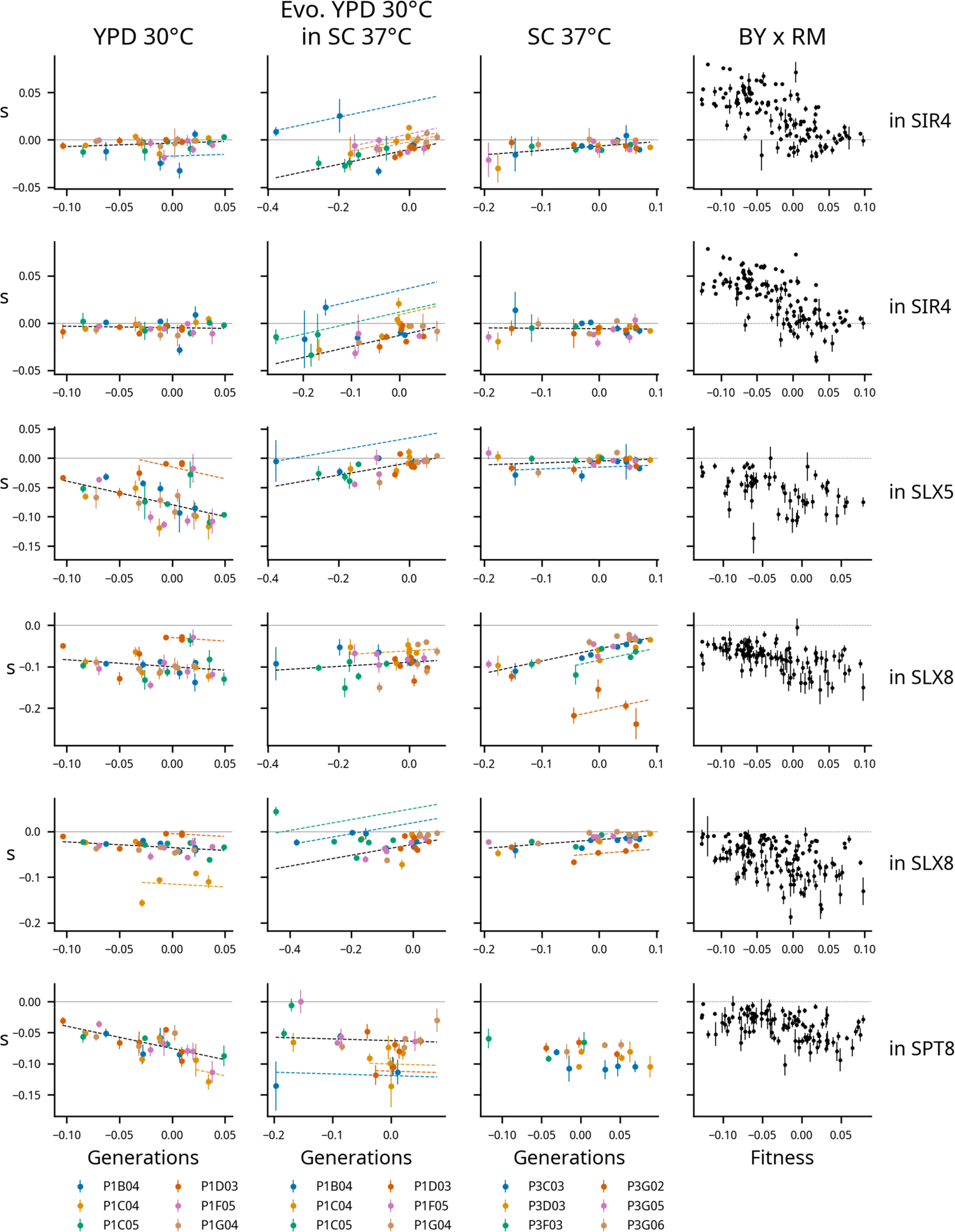
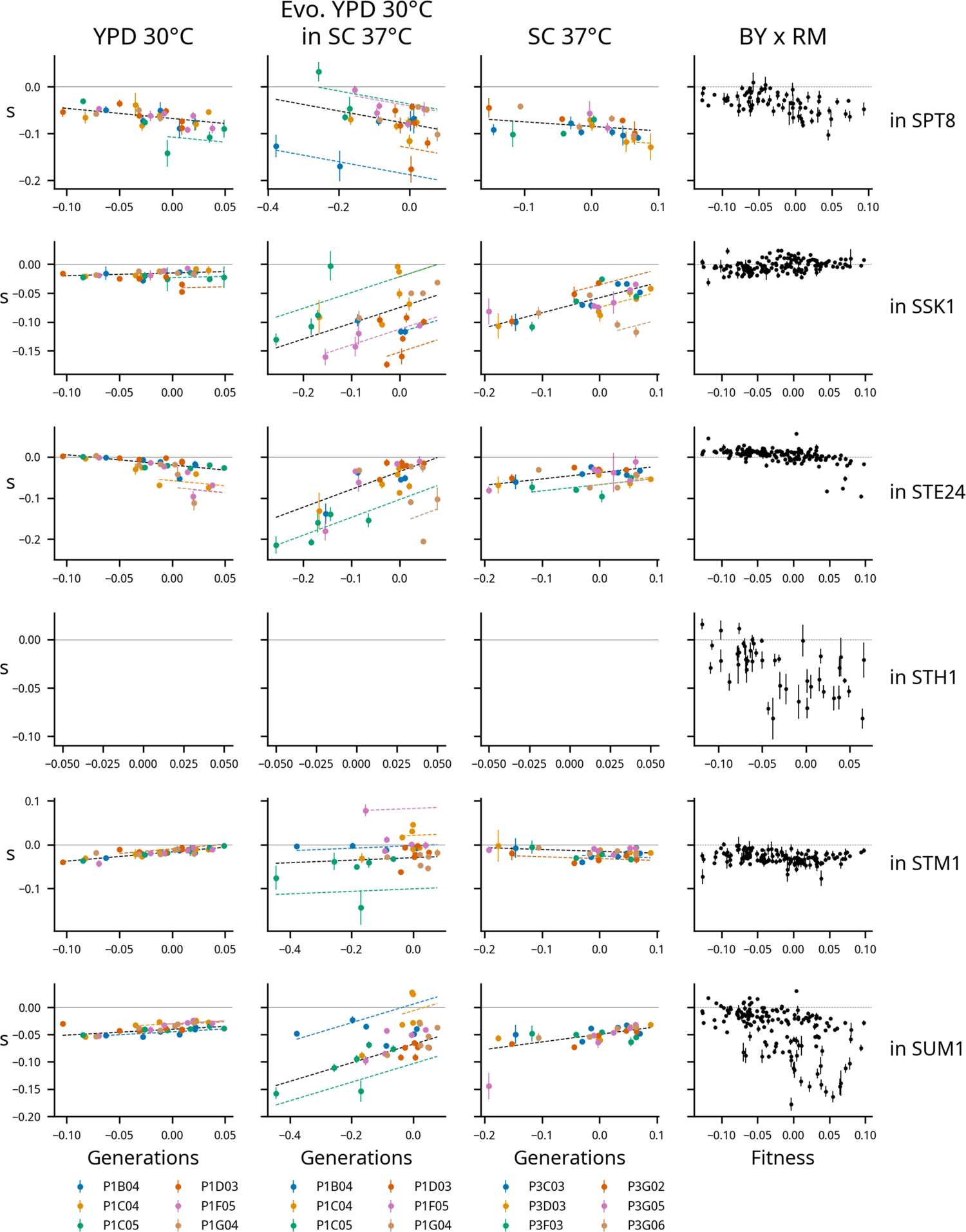
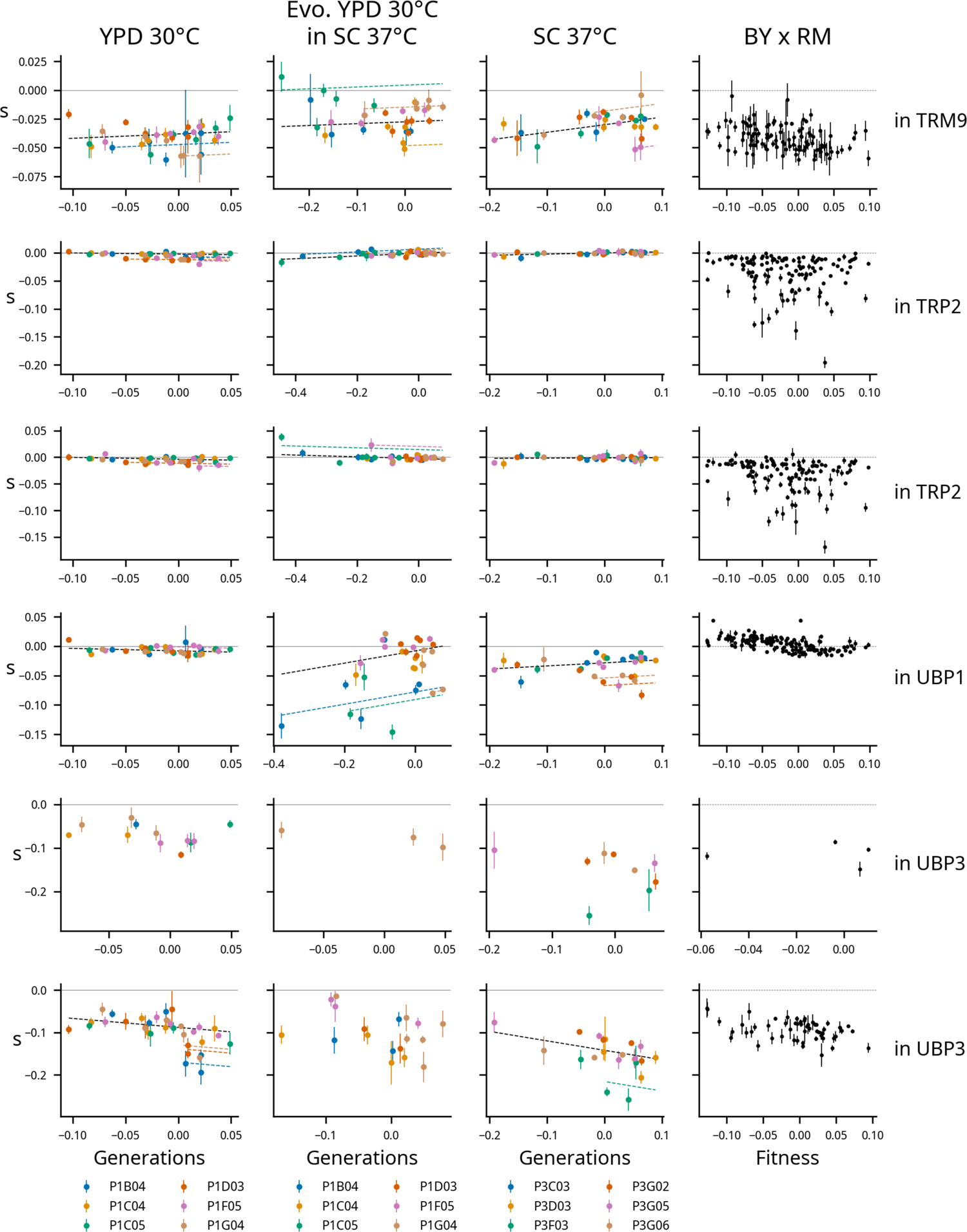
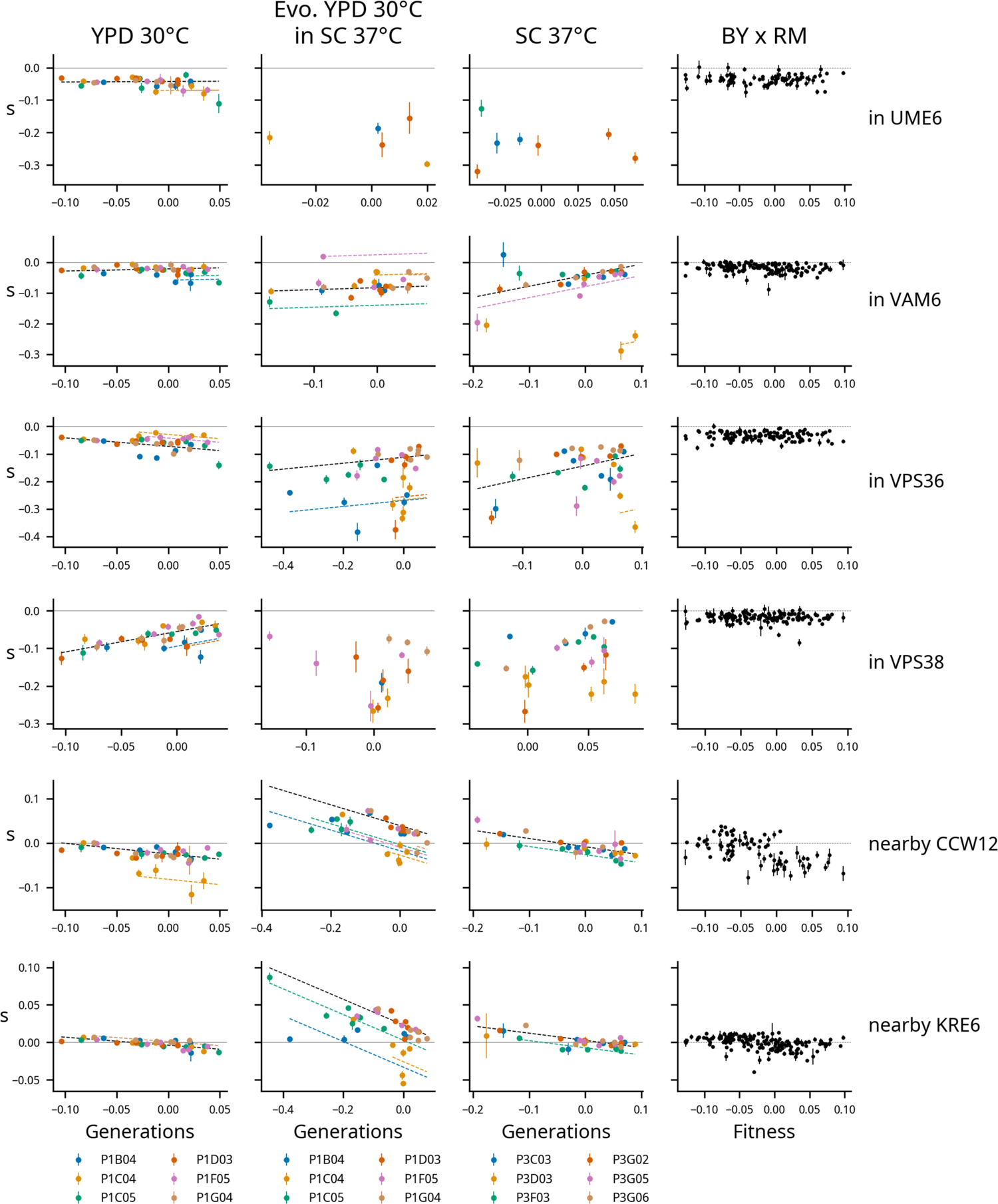
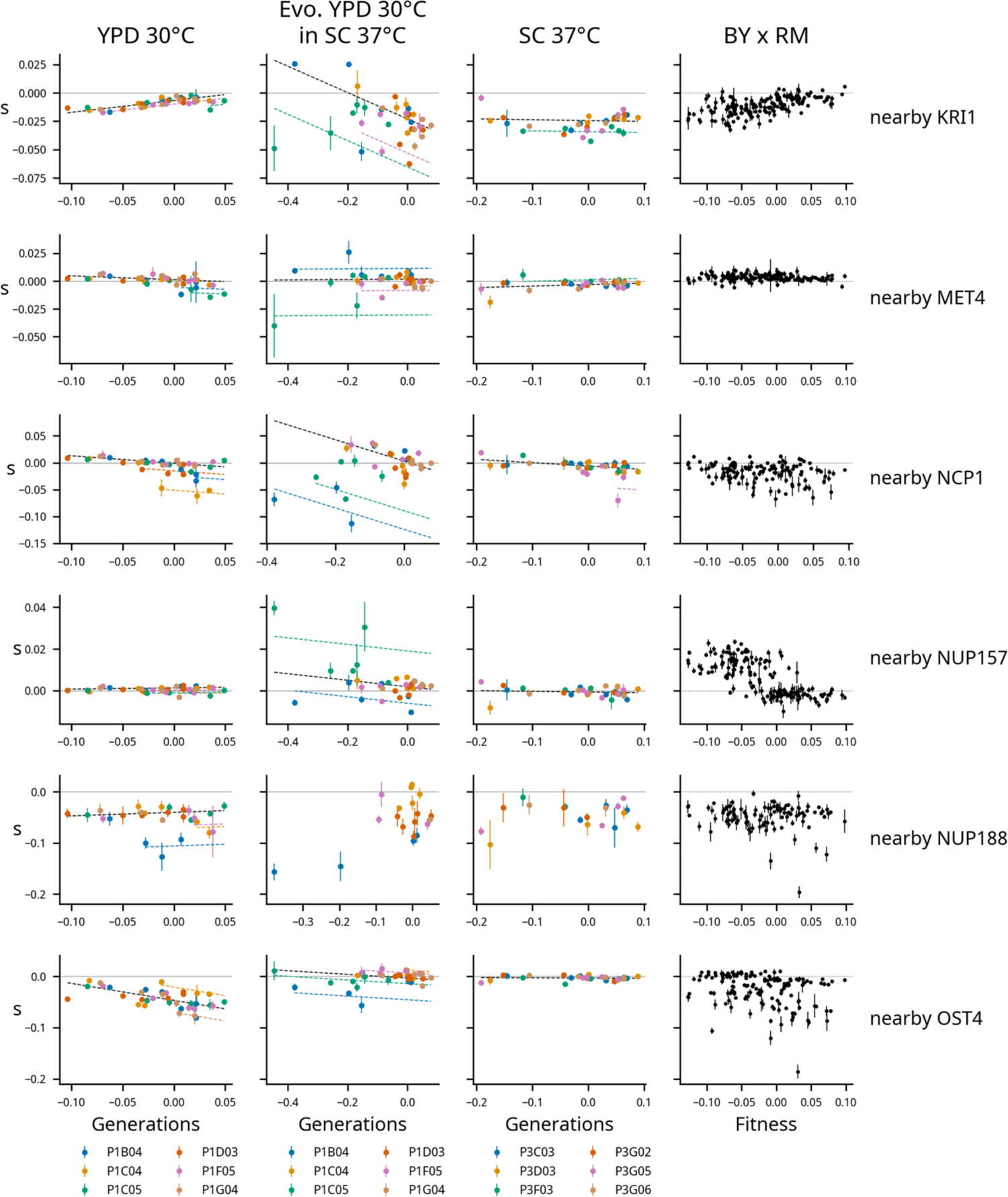
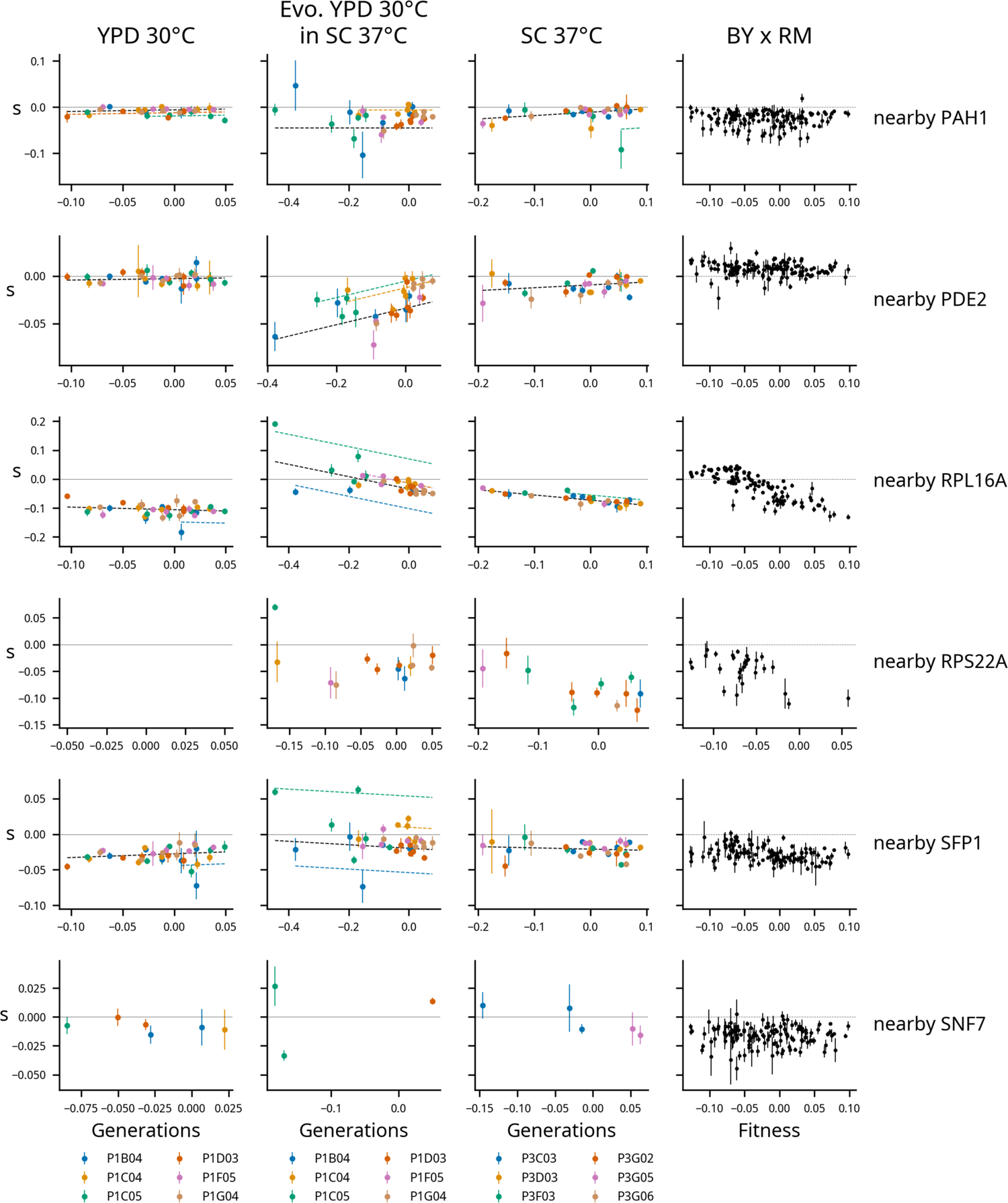
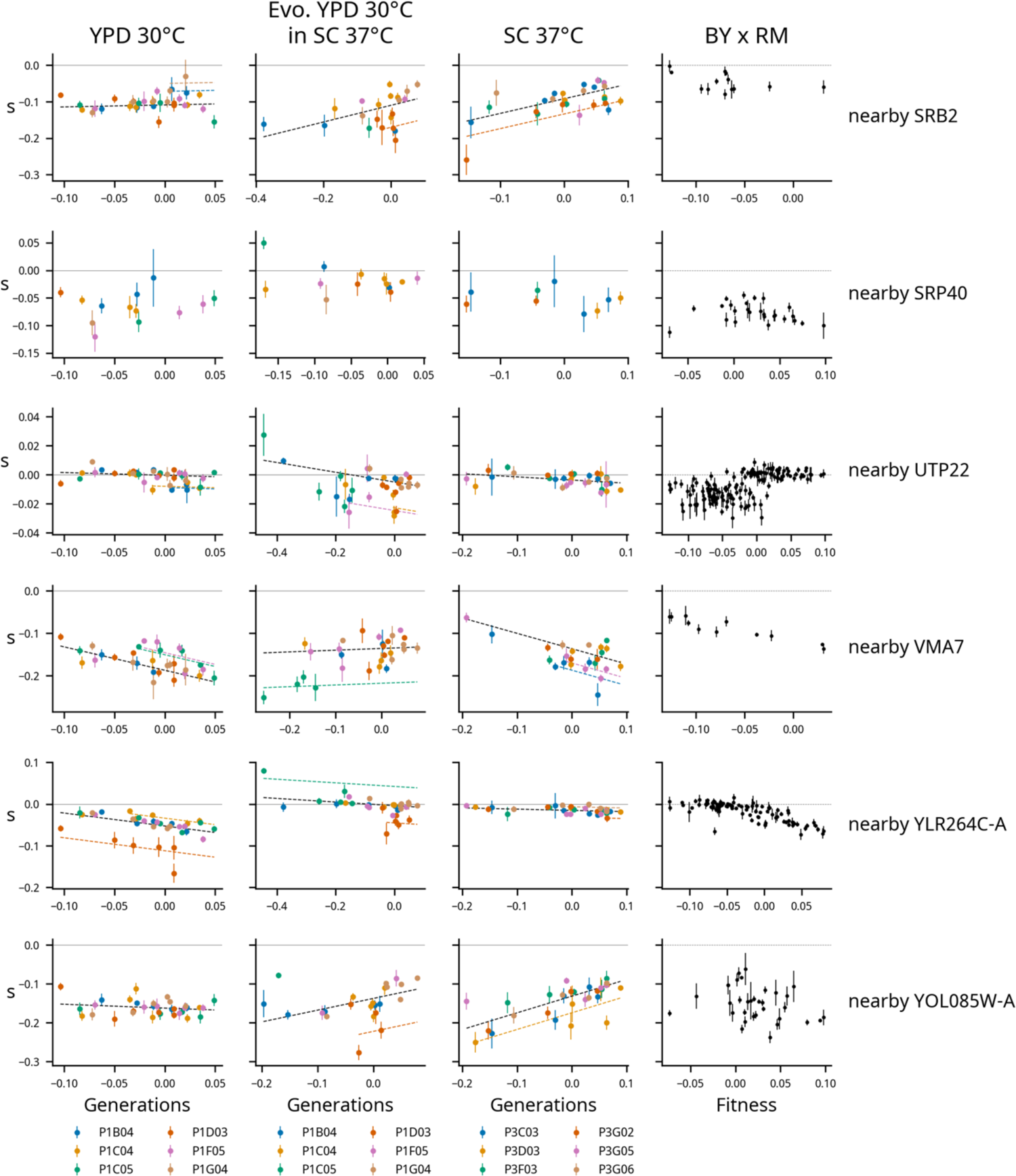
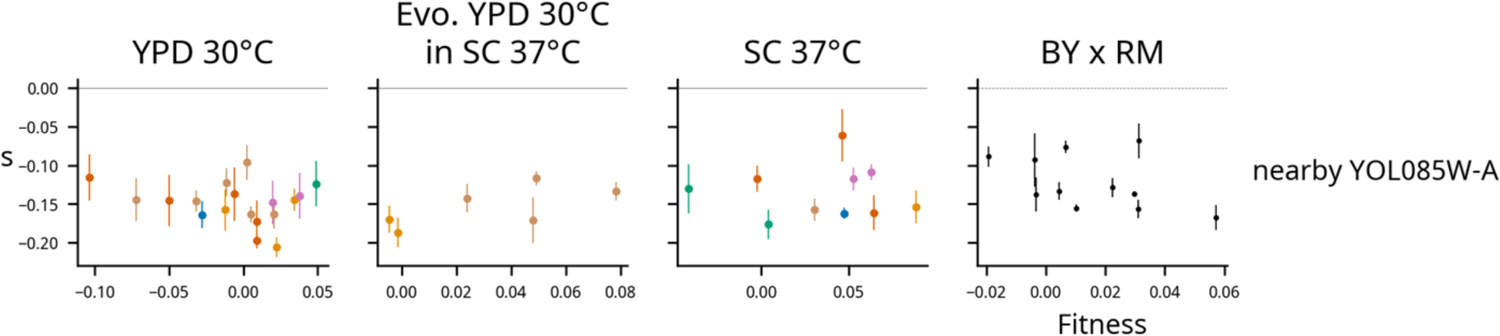
Determinants of fitness effects under the full model. Each row represents one mutation, labeled at right with by the gene it disrupts or the nearest start of a gene if it is intergenic. Each column represents a condition. The left three columns show the fitness effects of mutations as a function of fitness, with points colored by population. The right column shows the relationship between fitness and the fitness effect of the mutation in the segregants from a yeast cross (data from Johnson et al. (2019)). Model predictions are shown by dashed lines, and lines with contributions from indicator variables associated with a particular population are the same color as the points from that population.

